# Early neuroadaptations to an obesogenic diet identify the schizophrenia-related ErbB4 receptor in obesity-induced hippocampal abnormalities

**DOI:** 10.1101/2021.06.30.450398

**Authors:** Julio David Vega-Torres, Perla Ontiveros-Angel, Esmeralda Terrones, Erwin C. Stuffle, Sara Solak, Emma Tyner, Marie Oropeza, Ike Dela Peña, Andre Obenaus, Byron D. Ford, Johnny D. Figueroa

## Abstract

Childhood obesity leads to hippocampal atrophy and altered cognition. However, the molecular mechanisms underlying these impairments are poorly understood. The neurotrophic factor neuregulin-1 (NRG1) and its cognate ErbB4 receptor play critical roles in hippocampal maturation and function. This study aimed to determine whether altered NRG1-ErbB4 activities may partly explain hippocampal abnormalities in rats exposed to an obesogenic Western-like diet (WD). Lewis rats were randomly divided into four groups (12 rats/group): **1)** control diet+vehicle *(CDV)*; **2)** CD+NRG1 *(CDN)* (daily intraperitoneal injections: 5 μg/kg/day; between postnatal day, PND 21-PND 41); **3)** WD+VEH *(WDV)*; **4)** WD+NRG1 *(WDN)*. Neurobehavioral assessments were performed at PND 43-49. Brains were harvested for MRI and molecular analyses at PND 49. We found that NRG1 administration reduced hippocampal volume (7%) and attenuated hippocampal-dependent cued fear conditioning in CD rats (56%). NRG1 administration reduced PSD-95 protein expression (30%) and selectively reduced hippocampal cytokine levels (IL-33, GM-CSF, CCL-2, IFN-γ) while significantly impacting microglia morphology (increased span ratio and reduced circularity). WD rats exhibited reduced right hippocampal volume (7%), altered microglia morphology (reduced density and increased lacunarity), and increased levels of cytokines implicated in neuroinflammation (IL-1α, TNF-α, IL-6). Notably, NRG1 synergized with the WD to increase hippocampal ErbB4 phosphorylation and the tumor necrosis alpha converting enzyme (TACE/ADAM17) protein levels. Together, these data suggest a novel interaction between obesogenic diet exposure and TACE/ADAM17-NRG1-ErbB4 signaling during hippocampal maturation. Our results indicate that supraoptimal ErbB4 activities may contribute to the abnormal hippocampal structure and cognitive vulnerabilities observed in obese individuals.

**Highlights:**

- Obesogenic diet consumption during adolescence induces anxiety-like behaviors before the onset of obesity-related changes in metabolism.
- Obesogenic diet-driven abnormal behaviors co-occurred with alterations in hippocampal pro-inflammatory cytokine profiles.
- Obesogenic diet consumption attenuates exogenous NRG1 effects on hippocampal-related behaviors and structure.
- Exogenous NRG1 administration during adolescence resulted in reduced hippocampal volumes and domain-specific cognitive impairments.
- Exogenous NRG1 administration has potent immunomodulatory actions and alters hippocampal microglia morphology.

## Introduction

Childhood obesity is a severe medical condition affecting more than 340 million children and adolescents worldwide (World Health Organization, 2016). While obesity is a complex disease, poor diet quality is a significant factor contributing to the global obesity epidemic in children and adolescents (Moreno et al., 2010). Obesity and the consumption of imbalanced obesogenic diets rich in saturated fats and simple sugars have emerged as risk factors for the development of anxiety and other stress-related disorders (Scott et al., 2008; Perkonigg et al., 2009; Johannessen and Berntsen, 2013; Maguen et al., 2013; Mitchell et al., 2013; Kubzansky et al., 2014; Bartoli et al., 2015; Smith et al., 2015; Salari-Moghaddam et al., 2018). Evidence demonstrates that the relationship between obesogenic and anxiogenic phenotypes is bidirectional (Lopresti and Drummond, 2013; Michopoulos et al., 2016; Hruby et al., 2021). In other words, the presence of one increases the risk of developing the other. Thus, it has become crucial to understand better the specific substrates responsible for the intertwined pathophysiology associated with both conditions.

While the neural substrates impacted by obesity are very complex, hippocampal volume deficits have emerged as a prominent anatomical endophenotype (Raji et al., 2010; Cherbuin et al., 2015; Jacka et al., 2015; Mestre et al., 2017). Studies in animals and humans indicate that the hippocampus is essential for reducing energy intake (Kanoski et al., 2011; Stevenson and Francis, 2017; Liu and Kanoski, 2018) and stress reactivity (Jacobson and Sapolsky, 1991; Jimenez et al., 2018). As such, alterations in this brain region may contribute to the broad range of metabolic, cognitive, and emotional disturbances associated with obesity. We reported unique hippocampal vulnerabilities to the detrimental effects of obesogenic diets (Kalyan-Masih et al., 2016) consistent with human data (Bauer et al., 2015). However, the molecular and cellular basis underlying these vulnerabilities are poorly understood.

The hippocampal volumetric deficits observed in obese individuals could result from dysregulated neurotrophic factors and inflammatory signals. Neuregulin 1 (NRG1) is a trophic factor containing an epidermal growth factor (EGF)-like domain and acting via their cognate ErbB receptor tyrosine kinases. ErbB4, the only autonomous NRG1-specific activated tyrosine kinase, plays a critical role in regulating brain development and homeostasis. NRG1-ErbB4 signaling has been heavily implicated in neuronal migration, synaptic plasticity, and cognition (Mei and Xiong, 2008). NRG1-ErbB4 signaling regulates hippocampal function through increasing neuronal arborization (Gerecke et al., 2004), increasing gamma oscillations in the CA3 region of the hippocampus (Fisahn et al., 2009), modulating long-term potentiation at CA1 hippocampal synapses (Kwon et al., 2008), and modifying stress reactivity and behavior (Dang et al., 2016; Mahar et al., 2017; Clarke et al., 2018). Exogenous NRG1 administration increases hippocampal neurogenesis (Mahar et al., 2011, 2016) and improves learning and memory while rescuing dendritic and synaptic abnormalities in a mouse model of Alzheimer’s disease (AD) (Ryu et al., 2016). Similar neurorestorative properties were reported in a rat model of AD (Jalilzad et al., 2019).

Genetic studies have demonstrated strong associations between single-nucleotide polymorphisms in the ErbB4 gene and obesity (Locke et al., 2015; Salinas et al., 2016). Notably, while ErbB4 deletion predisposes mice to metabolic alterations implicated in obesity, NRG1 administration is known to improve metabolic health in obese mice by lowering blood glucose, improving insulin sensitivity, and reducing caloric intake (Ennequin et al., 2015a; Zeng et al., 2018; Zhang et al., 2018). Recent studies have reported an interplay between NRG1 signaling and high-fat diet (HFD) consumption in mice (Holm-Hansen et al., 2016; Zieba et al., 2019a; b). Zieba et al., demonstrated that HFD consumption during late adolescence has sex-specific effects on open-field (OF) exploration and fear conditioning (Zieba et al., 2019a). Female mice exposed to the HFD during late adolescence exhibited reduced OF exploration and increased cue freezing behaviors (Zieba et al., 2019a). Notably, female heterozygous transmembrane domain NRG1 mutant mice conferred protection against the effects of the HFD (body weight, OF distance traveled, cued freezing) (Zieba et al., 2019a).

Exogenous NRG1 administration confers potent anti-inflammatory effects. We and others have demonstrated that NRG1 anti-inflammatory actions extend to models of neuroinflammation, including cerebral malaria (Liu et al., 2018), nerve agent intoxication (Li et al., 2015), and ischemia (Simmons et al., 2016). Studies demonstrate that ErbB4 inhibition attenuates the beneficial effects of NRG1, supporting an essential role for ErbB4 in NRG1 actions (Tan et al., 2012; Guan et al., 2015; Gao et al., 2016; Zhang et al., 2017).

In this study, we hypothesized that NRG1 administration would confer protection against the detrimental effects of an obesogenic diet on neuroinflammation, hippocampal maturation, and anxiety-like behaviors. We reasoned that examining the responses of the hippocampal NRG1-ErbB4 system to obesogenic diets during adolescence may provide valuable insights into the potential contribution of this pathway to obesity-induced hippocampal deficits and anxiety. This study is the first to investigate the therapeutic potential of exogenous NRG1 to prevent the early consequences of an obesogenic environment during adolescence.

## Materials and Methods

### Animals

All the experiments were performed following protocols approved by the Institutional Animal Care and Use Committee (IACUC) at the Loma Linda University Health School of Medicine. Female Lewis rats with 8-10 male pups (postnatal day, PND 15) were purchased from Charles River Laboratories (Portage, MI, USA). Upon arrival, female rats were housed with their pups with free access to food and water. At PND 21, adolescent male pups were weaned, housed in groups (2 per cage), and experimental manipulations commenced. Animals were kept in customary housing conditions (21 ± 2 C, relative humidity of 45%, and 12-hour light/dark cycle with lights on at 7:00 AM). The body weights were recorded once a week or daily during the week of behavioral testing. Food consumption was quantified at least twice per week. The rats were never food or water restricted.

### Study Design

We used a 2 x 2 design with four groups: **(1)** Control diet, injected with vehicle (CDV); **(2)** Control diet, injected with neuregulin-1 (CDN); **(3)** Western diet, injected with vehicle (WDV); **(4)** Western diet, injected with neuregulin-1 (WDN). **Figure 1** summarizes the timeline of experimental procedures and behavioral tests. The matched low-fat purified control diet (CD, 5-gm% fat, product *#F7463*) and Western-like high-saturated fat diet (WD, 20-gm% fat, product *#F7462*) were obtained from Bio-Serv (Frenchtown, NJ, USA). The macronutrient composition and fatty acid profiles are detailed in a previous study and summarized in **Supplemental Table 1** (Vega-Torres et al., 2020b). Weight-matched adolescent rats (PND 21) were divided into 4 study groups: CDV, CDN, WDV, and WDN. NRG1 (5 micrograms/kilogram/day) or saline (vehicle) intraperitoneal injections were performed daily between 12:00-1:00 PM. The rationale for injection dosage and route was based on our previous reports describing the neuroprotective effects of this intervention (Li et al., 2012, 2015), NRG1 biodistribution, kinetics, and short half-life in plasma (Rösler et al., 2011; Zhang et al., 2018). Recombinant human NRG1-*β* 1/HRG1-*β* 1 EGF domain (NRG1) was provided by Dr. Byron D. Ford (manufactured by R&D Systems, catalog #*396-HB-050*). **Study 1 (behavioral and anatomical assessment):** All behavior testing sessions involved a 20-30-minute acclimation period to the testing facility. The rats were allowed to consume the custom diets until the completion of the study (PND 21-49). The initial group size was *n* = 18 for each group (divided into four cohorts). Twenty-four rats were randomly selected for MRI studies (*n* = 6 per group). **Study 2 (molecular markers investigation):** To minimize potential carryover effects between behavioral tests and molecular outcomes, we used a separate group of animals *without behavioral manipulations* for all the molecular outcomes reported in this study (RT-PCR, Western blot, and flow cytometry). For this group, an additional injection of NRG1 or saline was administered 1 h before euthanizing the animals at PND 42 (Rösler et al., 2011). The initial group size for Study 2 was *n* = 5-6 for each group (CDV, *n* = 5; CDN, *n* = 5; WDN; *n* = 6; WDV, *n* = 6).

**Figure 1.**
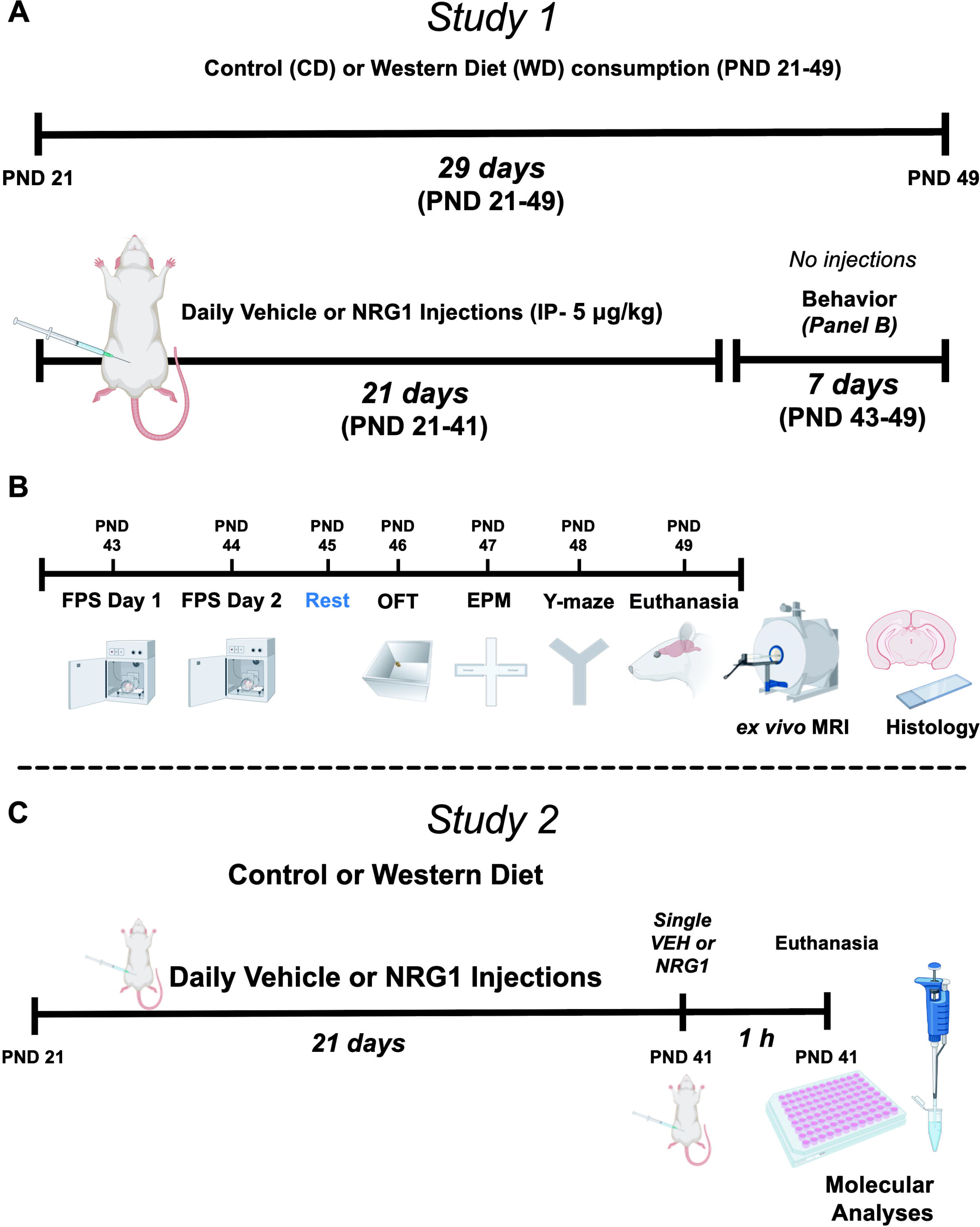
Experimental timeline and procedures. **(A)** Study 1 timeline and study groups. **(B)** Study 1 timeline of behavioral procedures, MRI, and histology. **(C)** Study 2 timeline of molecular analyses. Abbreviations: **CD**, control diet; **EPM**, elevated plus maze; **FPS**, fear-potentiated startle; **MRI**, magnetic resonance imaging; **NRG1**, neuregulin-1; **OFT**, open field test; **PND**, postnatal day; **VEH**, vehicle; **WD**, Western diet.

### Study 1

#### Fear-Potentiated Startle (FPS)

We used a *trace* FPS paradigm to evaluate hippocampal-dependent cued fear learning. Each FPS session started with a 5-min acclimation period (background noise = 60 dB). During the first session of the paradigm (fear training), the rats were trained to associate a light stimulus (conditioned stimulus, CS) with a 0.6 mA footshock (unconditioned stimulus, US). The conditioning session involved 10 CS + US pairing presentations. During each CS + US presentation, the onset of the shock (500 ms duration) occurred 3 sec after the offset of the 4 sec light. Light-shock pairings were presented in a quasi-random manner (ITI = 3-5 min). The cued fear acquisition was measured 24 h later. During cued fear acquisition (fear learning testing), the rats were first presented with 15 startle-inducing tones (leaders; 5 each at 90 dB, 95 dB, and 105 dB) delivered alone at 30 sec ITI. Subsequently, the rats were presented with 60 test trials. For half of these test trials, a 20 ms tone was presented alone (tone alone trials; 10 trials for each tone: 90 dB, 95 dB, and 105 dB). For the other half, the onset of the tone (20 ms duration) occurred 3 sec after the offset of the 4 sec light; 10 trials for each pairing: 90 dB, 95 dB, and 105 dB. The 60 trials were divided into 10 blocks of 6 test trials each, which included 3 tone alone trials and 3 light + tone trials. To conclude the testing session, the rats were presented with 15 startle-inducing tones (trailers; 5 each at 90 dB, 95 dB, and 105 dB) delivered at 30 sec ITI. The trials in this session were presented in a quasi-random order (ITI = 30 sec). The startle-eliciting tones had a 20 ms duration. We assessed fear learning by comparing the startle amplitude from light + tone trials (conditioned + unconditioned stimulus, CS + US) relative to tone alone trials (unconditioned stimulus, US). FPS data was reported as the change (delta) between US and CS + US [FPS = (Light + Tone Startle) − (Tone Alone Startle)].

#### Open Field Test (OFT)

The OFT was performed in an opaque acrylic open field maze (43.2 cm length × 43.2 cm width × 30.5 cm height). The rats were placed in the center of the field and recorded for 5 min using Ethovision XT (Noldus Information Technology). Behaviors were recorded at normal room lighting conditions (269 lux illumination). We used Ethovision to subdivide the arena into nine zones, which allowed for the evaluation of inner (center) and outer arena exploration. The chamber was cleaned with 70% ethanol between trials. The following equation was used to calculate the anxiety index: Anxiety index = 1 – [([center cumulative duration/total test duration] + [center total entries/total number of entries to center + corners])/2.

#### Elevated Plus Maze (EPM)

The near-infrared (NIR)-backlit EPM apparatus (catalog #ENV-564A, MedAssociates) consisted of two opposite open arms (50.8 × 10.2 cm) and two enclosed arms (50.8 × 10.2 × 40.6 cm) that were elevated 72.4 cm above the floor. Behaviors were recorded under low light conditions (< 10 lx). The rat was placed on the central platform facing an open arm and allowed to explore the maze for 5 min. The apparatus was thoroughly cleaned after each test session. Behaviors were recorded via a monochrome video camera equipped with a NIR filter. The time spent on both types of arms, the number of entries and duration in both types of arms, the latency to the first entry into any of the open arms (OA), closed arms (CA) entries, frequency in the head-dipping zone, and stretch attend postures (SAPs) were determined using Ethovision XT tracking software (Noldus Information Technology). From these data, we calculated the anxiety index according to our group and others (Cohen et al., 2012; Contreras et al., 2014; Kalyan-Masih et al., 2016; Vega-Torres et al., 2020b). Anxiety index = 1 – [([OA cumulative duration/total test duration] + [OA entries/total number of entries to CA + OA])/2.

#### Y-maze Spontaneous Alternation

The Y-maze apparatus was constructed of black Plexiglass with 3 identical arms spaced at an angle of 120 degrees (arms dimensions: 22” L × 4.5” W × 11.5” H). Each rat was placed at the end of the start arm and allowed to move freely through the maze during a 9-min session. The total number of arm entries and alternations from visited to unvisited arms was recorded using a video camera, tracked using Ethovision XT (Noldus Information Technology), and scored manually by two blinded investigators. Entries occur when all four limbs are within the arm. Each arm choice was recorded, and spontaneous alternation behaviors were scored as three successive choices that include one instance of each arm (a triad). The percentage of spontaneous alternations was calculated as the total number of alternations / (total arm entries – 2) x 100. This measure is considered to reflect short-term memory in rats. The alternate arm returns and same arm returns were also scored for each animal to investigate attention within spontaneous working memory. The number of total alternations was calculated as an index of locomotor activity, and rats exhibiting 0 entries were excluded from analyses. Behaviors were recorded at normal room lighting conditions (269 lx). The maze was cleaned with 70% ethanol between trials.

#### Magnetic Resonance Imaging

All fixed brain tissues underwent *ex vivo* high-resolution imaging using a 9.4 T Bruker Avance Imager (Bruker Biospin, Billerica, MA). All the data were collected with a 2.2-cm field of view, 0.55-mm slice thickness, and a 200x180 acquisition zero-filled to a final 256x256 matrix. T2-weighted RARE (rapid acquisition with relaxation enhancement; T2 RARE) were collected with the following parameters: repetition time (TR) / echo time (TE) = 6482 msec / 49.3 msec, RARE factor of 4, for a final resolution of 110x111x550 microns. The total imaging time was 39 min.

The experimenters were blinded to the group designation during MRI analysis. MRI scans were eddy corrected, and the cranium was removed using MatLab. The Waxholm rat brain atlas was fit to each animal, and regional labels were applied with Advanced Normalization Tools (ANTs). Regional volumes were extracted for 72 regions (144 bilateral regions). All data were extracted and summarized in Excel. Given our interest in hippocampal atrophy, we focused on hippocampal volumes and subfields.

### Neurohistology

#### Tissue Preparation and Immunohistochemistry

Before euthanasia, the animals were deeply anesthetized with an intraperitoneal overdose injection of Euthasol (150 mg/kg; Virbac, Fort Worth, TX) and sacrificed via transcardiac perfusion with 10% sucrose (isotonic at 9.25% in distilled water; prewash) and 4% paraformaldehyde (PFA; fixative) using the Perfusion Two™ Automated Pressure Perfusion System (Leica Biosystems, Buffalo Grove, IL) (*n* = 12; 3 rats per group). The brains were removed from the cranial vault after MRI imaging and washed with phosphate-buffered saline (PBS) and cryoprotected with sucrose (30%) for 12–16 h at 4°C before embedding in Tissue-Tek^®^ O.C.T.™ compound (Sakura, Torrance, CA, United States). Three (3) brains per group were later sectioned with a cryostat (Cryocut 1800 Cryostat, Leica Biosystems, Vista, CA, USA) at 25 μm thickness in the coronal plane. Regions of interest included dorsal and ventral areas of the CA1 region of the hippocampus of both hemispheres with a distance from Bregma between -4.6 and -5.6. Sections were mounted in glass slides, and immunohistochemistry staining was performed with the microglia/macrophage marker, ionized calcium-binding adaptor protein molecule 1 (Iba-1). First, sections were incubated in a quenching buffer (10% methanol, 1% hydrogen peroxide in PBS) to block endogenous peroxide activity. To block the signal from endogenous avidin, biotin, and biotin-binding proteins in tissue, we used Avidin/Biotin blocking kit (Abcam, USA) followed by incubation with normal serum (5% Donkey and 5% Goat serum) with 1% Triton-X in PBS. Sections were then incubated overnight with a polyclonal rabbit anti-Iba1 primary antibody (1:750, rabbit anti-Iba1, catalog #019-19741, Wako, USA) followed by secondary biotinylated antibody incubation (1:200, goat anti-rabbit IgG, Vector Labs Inc.). The Vectastain Elite ABC HRP kit and DAB peroxidase substrate kit with nickel (Vector Labs Inc., Burlingame, CA, USA) were used to visualize staining according to manufacturer instructions. Following staining, slides were dehydrated in ethanol (79, 95, and 100%), rinsed in Xylene, sealed with Permount Mounting Medium (Fisher Chemical), and coverslips applied for subsequent visualization.

#### Image Acquisition and Segmentation Processing for Microglial Morphological Characterization

Blinded investigators carried out image acquisition and analyses. Images of tissue sections that were DAB-stained with the Iba-1 antibody were obtained using a Keyence BZ-9000 series microscope (Keyence Corporation, Osaka, Japan) in brightfield at a 40× magnification. To obtain a single high-quality full-focused composite image that showed detailed magnification of ramified processes of the cells and facilitate image segmentation, a multi-plane virtual-Z mode image with 10 images (1 μm thickness) in 20 μm depth of tissue section was captured and later combined using public domain software *ImageJ* (https://imagej.nih.gov/ij/). Hippocampal dorsal and ventral CA1 areas were scanned to obtain at least 3 images per region of interest per hemisphere covering a field of view area of 1.2 mm. Each acquired image was saved as a TIFF file and included at least six (6) Iba-1+ cells.

Images were processed in an unbiased and systematic way to obtain a filled image. For this purpose, a series of steps were performed using ImageJ with the Fiji plugin (https://imagej.net/Fiji). First, a single high-quality image was obtained by merging all the single images and processed under a minimum threshold feature to soften the background and enhance the contrast. Subsequently, the image was transformed to 8-bit grayscale and binarized to obtain a black and white image by applying a previously established threshold. To obtain a cell image consisting of continuous pixels, manual editing of each image was performed. A set threshold of pixel number addition (no more than 4 pixels) to join processes belonging to a selected cell was established and performed systematically. A 2-pixel set threshold was applied to separate ramification to neighboring cells when applicable. This step was carefully performed under the view of the original z-stack images with special care to avoid bias. The final filled shape image was saved and later analyzed with *FracLac* V2.5 for *ImageJ* (Karperien, A., FracLac for ImageJ; http://rsb.info.nih.gov/ij/plugins/fraclac/FLHelp/Introduction.htm) (Karperien et al., 2013) to quantify the morphological changes of microglial cells in each experimental group. Foreground pixel quantification of the filled binary images per object (cell) was done with the Box Counting (BC) feature, which counts the number of boxes containing any foreground pixels along with the successively smaller caliber grids. BC box size scale was obtained as a Power Series where the base is raised to the exponent added to it to make successive sizes. Finally, the slope for each object was the average of 12 measurements with different and random placements of the grid. Box Counting and Convex and Hull Area measurements were exported using Excel (Microsoft, Redmond, WA, USA) for record and tabulation. Graphpad Prism version 9.0 (Graphpad Software, La Jolla, CA, USA) was used for statistical analyses.

### Study 2

#### Western Blot

We used a separate group of animals *without behavioral manipulations* for Western blot experiments to minimize potential carryover effects of behavioral tests on molecular outcomes. For this group, an additional injection of NRG1 or saline was administered 1 h before euthanizing the animals to investigate the acute activation of NRG1-ErbB4 signaling pathways. The rats were euthanized with intraperitoneal administration of Euthasol (Virbac) and perfused transcardially with PBS. The rats were rapidly decapitated, and the brains were isolated, and the hippocampus was dissected out. The tissue was immediately homogenized in 600 μL MPER Extraction Buffer (Thermo, 78501) (supplemented with protease and phosphatase inhibitors) using Bullet Blender. Homogenates were centrifuged for 10 min at 14,000 x g at 4°C. The supernatant was removed, aliquoted, and stored at -80°C until the day of the experiment. Protein concentration was determined using the Pierce BCA Protein Assay (Thermo, 23225). Homogenates were then mixed with 4x Protein Sample Loading Buffer (LI-COR Biosciences, 928-40004), NuPAGE Sample Reducing Agent (10x) (Thermo, NP0009), and subsequently heated at 95°C for 5 min. Fifty micrograms (50 µg) of protein were separated on NuPAGE 4-12% Bis-Tris gels (Invitrogen). Proteins were transferred to iBlot Gel Transfer Stacks Nitrocellulose Membranes (Invitrogen) using the iBlot 2 System (Thermo). The blots were washed and blocked for 1 h in 5% dry milk or Intercept (TBS) blocking buffer (LI-COR, 927-60001) and incubated with primary antibodies overnight at 4°C (diluted in 1x TBS with 0.1% Tween® 20). Primary antibody dilutions are described in **Supplemental Table 2**. Blots were subsequently washed in 1x TBS with 0.05% Tween® 20 and incubated with appropriate fluorescent secondary antibodies (LI-COR; diluted at 1:30,000 in 1x TBS with 0.1% Tween® 20) for 90 min at room temperature. Membranes were washed with 1x TBS before imaging. Images were obtained using Odyssey® Fc Imager (LI-COR).

### Bead-based Cytokine Detection and Metabolic Status Measurements

To examine neuroinflammation from hippocampal homogenates, we performed a multianalyte bead-based immunoassay (LEGENDplex™ Custom Panel of Rat Inflammation markers; Biolegend, San Diego, CA, USA) to provide fast and quantitative information on the WD and NRG1 treatment effects in levels of relevant pro and anti-inflammatory cytokines and chemokines. Rats were euthanized with an intraperitoneal administration of Euthasol (Virbac, Fort Worth, TX, USA) and perfused transcardially with ice-cold PBS. Animals were rapidly decapitated, and the brains were isolated. For total protein extraction, hippocampal brain tissue was homogenized in a buffer containing 20 mmol/L Tris-HCl (pH 7.5), 150 mmol/L NaCl, 1 mmol/L PMSF, 0.05% Tween-20, and a Protease Inhibitor Cocktail (cOmplete™ Mini Protease Inhibitor Cocktail, Roche Diagnostic, USA). Samples were then agitated for 30 min at 4°C and centrifuged at 12,000 rpm for 20 min at 4°C, the supernatant was then removed. Protein concentration was quantified in triplicates using the DC™ Protein Assay (Bio-Rad Laboratories, Hercules, CA, USA) according to the manufacturer’s instructions, and absorbance was read at 655 nm using Spectramax i3x detection platform (Molecular Devices, Sunnyvale, CA, USA).

Cytokine evaluation of hippocampal IL-10, IL-1α, IL-6, IL-1β, TNF-α, IL-33, GM-CSF, CXCL-1, CCL2, IL-18, IL-12p70, IL-17a, and INF-γ were quantified in triplicates using the bead-based Assay LEGENDplex™ protocol for V bottom plate, according to the manufacturer’s instructions. Samples were diluted at 1:4 with Assay Buffer. Cytokine levels were determined using antibodies for each analyte covalently conjugated to a set of microspheres in a carbodiimide crosslinking reaction and detected by a cocktail of biotinylated antibodies. Following binding of streptavidin–phycoerythrin conjugate, the fluorescent reporter signal was measured using a MACSQuant® Analyzer 10 (Miltenyi Biotech, *Bergisch Gladbach*, Germany). FCS files were evaluated with the LEGENDplex™ Data Analysis Software Suite using the Qognit cloud base platform (Biolegend, San Diego, CA, USA). A five-parameter logistic curve-fitting method was used to calculate concentrations from Median Fluorescence Intensity and normalized to the amount of protein in each sample. The results are reported as pg/mg of protein.

### Metabolic and Inflammatory Assessments

Please refer to published work from our lab for detailed protocols on the metabolic biomarker measures used in this study (Kalyan-Masih et al., 2016).

#### Blood glucose

post-prandial blood glucose levels were measured in anesthetized rats by cutting the tail tip right before inducing euthanasia. Glucose levels were measured using a glucometer (OneTouch UltraMini® manufactured by LifeScan Inc, Milpitas, CA, USA) and reported as mg per dL. Study 2 rats were used for the molecular experiments (same rats used for Western blot analyses). Plasma was collected and stored at -80 °C.

#### Corticosterone

Circulating plasma levels of corticosterone (CORT) were measured by ELISA (Enzo Life Sciences, ADI-900-097 Ann Arbor, MI, USA) according to the manufacturer’s instructions. Plasma samples were diluted with kit assay buffer (1:5 dilutions) and ran in triplicates. The absorbance was measured at 405 nm with 570 nm correction using the SpectraMax i3X detection platform (Molecular Devices, Sunnyvale, CA). Corticosterone levels were determined as the percentage bound using a standard curve with a detection range between 20,000 - 31.25 pg/mL. Values are reported as picograms per milliliter.

#### Leptin

Plasma samples were used to evaluate leptin levels, samples were diluted with kit assay buffer (1:3 dilutions), ran in triplicates, and measurements were made using an enzyme-linked immunosorbent assay ELISA kit (Leptin Rat ELISA Kit Abcam, ab100773; Cambridge, MA, USA) according to manufacturer’s instructions. Plates were read at 450 nm using the SpectraMax i3X (Molecular Devices, Sunnyvale, CA USA). Specific concentrations for each sample were determined as mean absorbance using a standard curve of samples ranging from 0 to 8,000 pg/mL. Values were then calculated and reported as picograms of leptin per mL.

#### Insulin

Plasma samples were assayed for insulin content using an insulin ELISA kit (Abcam, ab100578, Cambridge, MA, USA) following the manufacturer’s instructions. Samples were diluted with Assay Diluent (1:5 dilution) in triplicates, and levels measured against a standard curve from 0-300 uIU. Samples were read at 450 nm using the SpectraMax i3X (Molecular Devices, Sunnyvale, CA, USA). Insulin concentration is reported as uIU per mL.

#### Triglycerides

Plasma samples were used to determine triglycerides levels with the TG assay kit for quantification (Abcam, ab65336; Cambridge, MA, USA) as previously described (Kalyan-Masih et al., 2016). Samples were diluted in with kit assay buffer (1:3 dilution) and ran in triplicates. Triglyceride levels were determined using the SpectraMax i3X plate reader (Molecular Devices, Sunnyvale, CA USA). TG concentration was determined as the mean absorbance at 570 nm using a standard curve of samples ranging from 0 to 10,000 pmol. TG concentration was calculated and reported as milligrams of TG per dL

#### FGF-21

Plasma samples were assayed for FGF-21 levels using an ELISA kit from R&D Systems (MF2100, R&D Systems, Minneapolis MN, USA) according to the manufacturer’s instructions. Samples were diluted with Assay Diluent (1:2 dilution) in triplicates and levels measured against a standard curve from 0-2,000 pg/mL. Samples were read at 450 nm and corrected at 540nm using the SpectraMax i3X (Molecular Devices, Sunnyvale, CA USA). FGF-21 concentration is reported as ng/mL.

#### NRG1

Hippocampal homogenate samples were used to determine NRG1 levels with the Human NRG1-β1 ELISA (DY377, R&D Systems, Minneapolis MN, USA). Samples were diluted in kit assay buffer (1:6 dilution) in triplicates. NRG1 levels were determined using the SpectraMax i3X plate reader (Molecular Devices, Sunnyvale, CA USA) as the mean absorbance at 450nm with correction at 540 nm using a standard curve of samples ranging from 0 to 4,000 pg/mL. NRG1 concentration was calculated and reported as nanograms of NRG1 per mL.

### Statistics

We analyzed the data using GraphPad *Prism* version 9.0 (Graphpad Software, La Jolla, CA, USA). Shapiro-Wilk statistical analyses were used to determine sample distribution. The Brown-Forsythe test was used to test for the equality of group variances. Two-way analysis of variance (ANOVA) was used to examine the effect of the diet type, treatment, and interaction between factors on outcome measures. Three-way ANOVA was used to investigate the impact of the diet type, treatment, hippocampal regions, and interactions between factors on microglial morphology. Mixed-effects model analyses were used to examine the effect of the diet type, treatment, ErbB4 isoform type, and interactions on hippocampal ErbB4 mRNA levels. Multiple comparisons were made using Sidak’s (for one significant main effect; within-group) or Tukey’s test (for more than one significant main effect or significant interaction effect; between-group). The ROUT method was used to determine outliers. Principal component analyses (PCA) were performed to reduce the microglia morphological descriptors’ dimensionality by using centering/scaling methods. We considered differences significant when *p* < .05. The data is shown as the mean +/-standard error of the mean (SEM).

### Images and Figures

Illustrations were prepared using BioRender (www.Biorender.com), OmniGraffle Pro (The Omni Group; Seattle, WA), and Prism (Graphpad Software; La Jolla, CA).

## Results

### Early impact of macronutrient profile of diet on biometric parameters

Following the reports indicating that exogenous NRG1 improves metabolic outcomes implicated with obesity (Ennequin et al., 2015a; b; Caillaud et al., 2016; López-Soldado et al., 2016; Zhang et al., 2018), we tested whether exogenous NRG1 administration prevents/delays the onset of obesity symptoms. We postulated that exogenous NRG1 administration would attenuate the WD-induced hippocampal structural impairments and alterations in cued learning, spatial working memory, inflammatory signals, and synaptic integrity. **Figure 1** details the experimental timeline and experimental groups (*CDV*, control diet + vehicle injections; *CDN*, control diet + NRG1 injections; *WDN*, Western diet + vehicle injections; *WDN*, Western diet + NRG1 injections). Here, we found that adolescent rats exposed to the Western-like high-fat diet (WD) for three weeks exhibited significantly increased body weight, caloric intake, and post-prandial blood glucose levels relative to the rats that consumed the control diet (CD) (*p* < .05; **Table 1**). However, three weeks of consuming the obesogenic WD was not sufficient to significantly alter critical peripheral obesity biomarkers, including leptin, insulin, triglycerides, and corticosterone (*p* > .05; **Table 1**). Subchronic exogenous NRG1 administration had no significant effects on the biometric parameters assessed in this study (*p* > .05; **Table 1**).

**Table 1.**
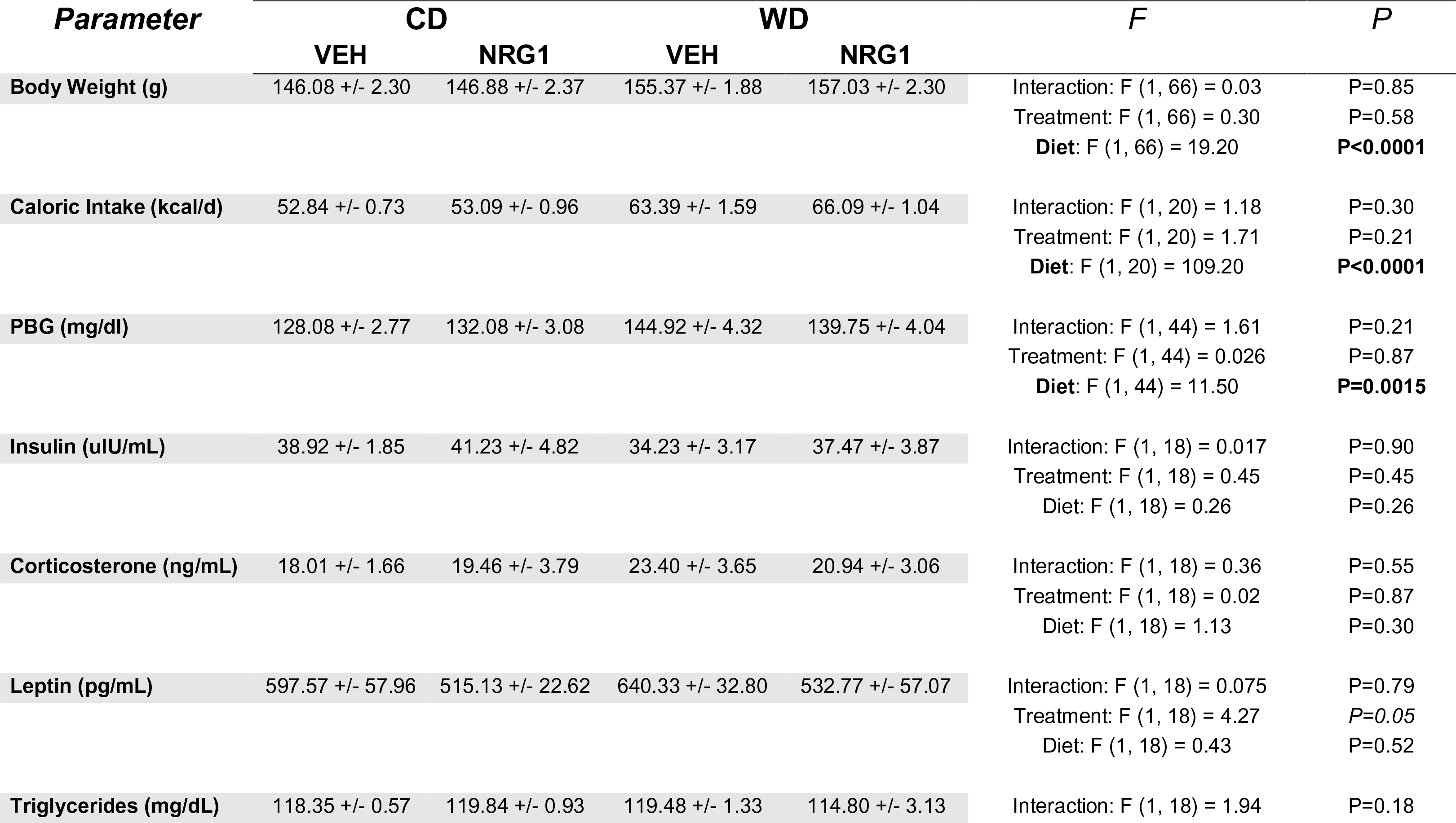
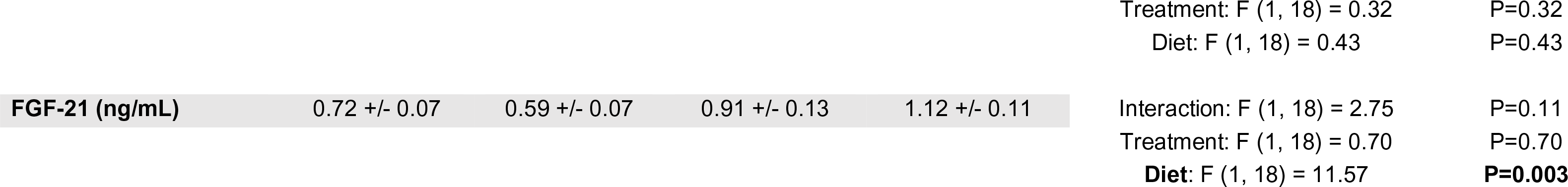
Detailed biometric parameters findings and statistics. The adolescent rats that consumed the WD exhibited an increase in body weight, food consumption, circulating FGF-21 levels, and post-prandial blood glucose levels relative to animals consuming the control diet at postnatal day 41. Three weeks of consuming the WD was not sufficient to alter plasma insulin, leptin, corticosterone, and triglyceride levels in adolescent rats. Exogenous NRG-1 administration did not have a significant effect on any of the biometric parameters measured.

### Exogenous NRG1 administration during adolescence reduces trace cued fear learning

Recent work from our lab has focused on identifying early (mal)adaptive responses to obesogenic environments (Kalyan-Masih et al., 2016; Vega-Torres et al., 2018, 2019a, 2020b; Santana et al., 2021). Having demonstrated that exposure to an obesogenic diet during adolescence alters the neural and behavioral substrates implicated with emotional regulation, we asked whether these impairments can be attenuated by administering neuregulin-1 (NRG1), a neurotrophic factor with potent neuroprotective and anti-inflammatory properties (Li et al., 2012; Depboylu et al., 2015; Guan et al., 2015; Simmons et al., 2016).

Trace conditioning is thought to reflect hippocampal-dependent declarative memory. Hippocampal lesions impair acquisition and expression of trace conditioning, while having little effect on the acquisition of amygdalar-dependent delay conditioning (McGlinchey-Berroth et al., 1997; Trivedi and Coover, 2006). Based on our findings reporting structural impairments in hippocampus of rats that consumed an obesogenic diet during adolescence (Kalyan-Masih et al., 2016), we decided to investigate the effect of the WD and exogenous NRG1 on trace conditioning. We found no significant interaction [*F*_(1, 43)_ = .79, *p* = .38], treatment [*F*_(1, 43)_ = .05, *p*= .82], or diet [*F*_(1, 43)_ = 1.82, *p* = .19] effects on foot shock reactivity (**Figure 2A**). We found that daily NRG1 administration (5 μg/kg/day for 21 days between postnatal day, PND21-PND41) led to a significant reduction in fear-potentiated startle (FPS) responses at 24 h following trace cued fear conditioning. Analyses showed a significant interaction [*F*_(1, 43)_ = 4.57, *p* = .038] effect in FPS responses, with no significant treatment [*F*_(1, 43)_ = 2.81, *p* = .10] or diet [*F*_(1,43)_ = .40, *p* = .53] (**Figure 2B**). Post hoc testing revealed significant differences between the CDV and CDN rats (*p* = .021). When analyzing each separate tone intensity, we found a stimulus effect (tone alone vs. light + tone) for 90 dB [*F*_(1, 40)_ = 64.82, *p* < .0001], 95 dB [*F*_(1, 44)_ = 27.71, *p* < .0001], and 105 dB [*F*_(1, 43)_ = 37.61, *p* < .0001]. Interestingly, only CDV rats exhibited significant differences between the stimulus type (**Supplemental Figure 1**). This phenotype is consistent with previous reports from our lab proposing that obesity and the consumption of obesogenic diets saturates fear and emotional circuits (Vega-Torres et al., 2018).

**Figure 2.**
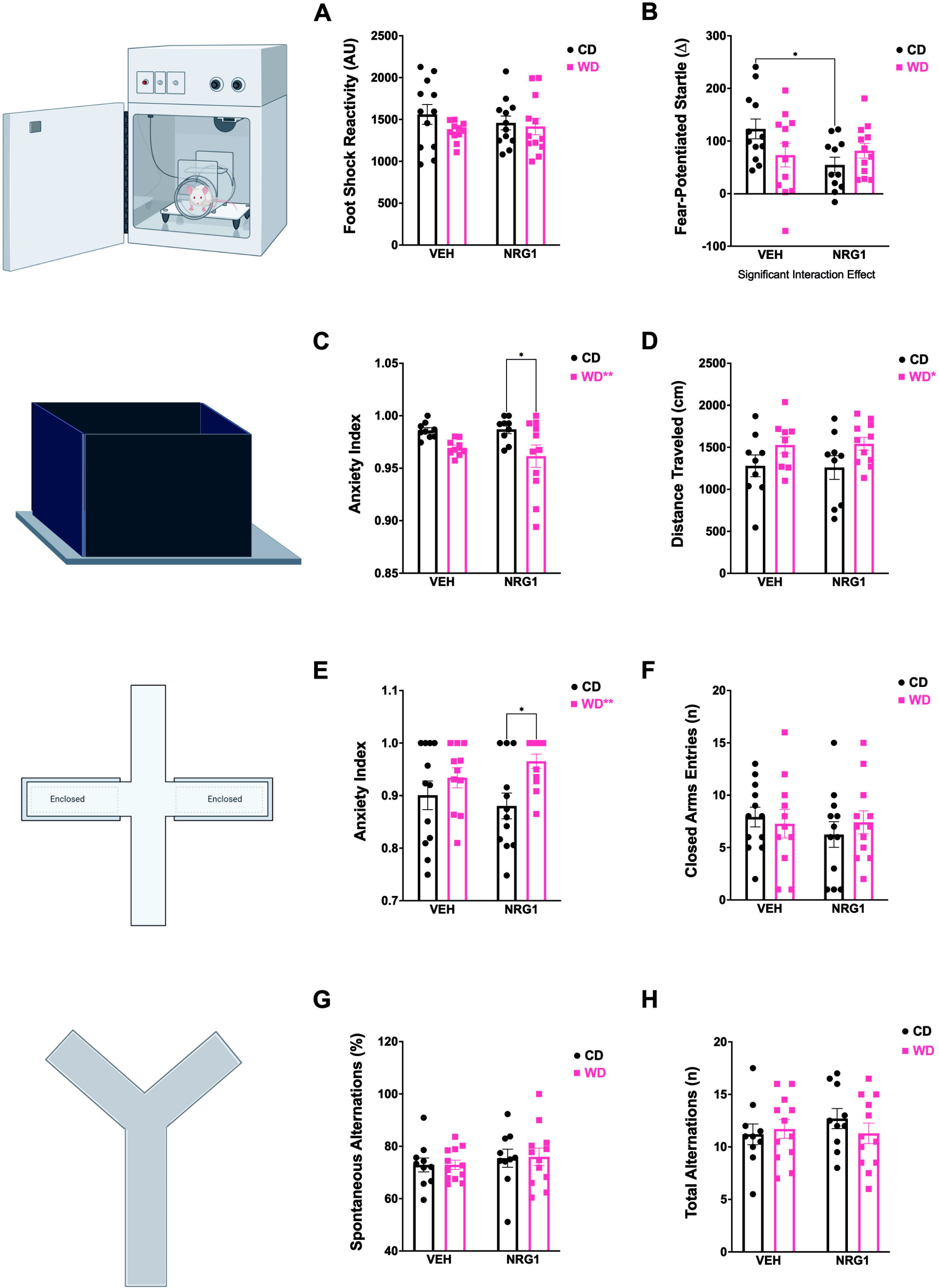
Behavioral phenotypes associated with early access to an obesogenic diet and prolonged NRG1 administration. **(A)**Startle reactivity to the foot shocks was not affected by treatment [*F*_(1, 43)_ = .05, *p*= .82] or diet [*F*_(1, 43)_ = 1.82, *p* = .19]. **(B)** Fear potentiated-startle (FPS) responses were determined from trace conditioning protocol and showed a decrease in FPS (interaction: [*F*_(1, 43)_ = 4.57, *p* = .038]). CDN group had a significant decrease in FPS responses when compared to CDV (*p* = .021). **(C)** Open-field test (OFT) anxiety index was affected by the diet [*F*_(1, 34)_ = 9.69, *p* = .004]. WDN rats exhibited reduced indices of anxiety in the OFT relative to CDN rats (*p* = .019). **(D)** The diet had a significant effect on total distanced traveled in the OFT [*F*_(1, 34)_ = 5.70, *p* = .023]. **(E)** Elevated-plus maze (EPM) anxiety index was affected by the diet [*F*_(1, 43)_ = 7.27, *p* = .009]. WDN rats demonstrated higher anxiety indices in the EPM when compared to CDN (*p* = .017). **(F)** Number of closed arms entries in the EPM was not affected by treatment [*F*_(1, 43)_ = .434, *p* = .514] or diet [*F*_(1, 43)_ = .051, *p* = .822]. **(G)** Spontaneous alternations in the Y-maze was not affected by treatment [*F*_(1, 40)_ = .97, *p* = .33] or diet [*F*_(1, 40)_ = .011, *p*= .92]. **(H)** Treatment [*F*_(1, 40)_ = .32, *p* = .58] or diet [*F*_(1, 40)_ = .22, *p* = .64] had no effects on total number of alternations in Y-maze. For all behaviors: sample size = 12 rats / group (before outlier testing).

#### WD consumption during adolescence increases indices of anxiety in the EPM

Open-field test (OFT) behavior analyses revealed a significant diet [*F*_(1, 34)_ = 9.69, *p* = .004], and no significant interaction [*F*_(1, 34)_ = .43, *p* = .52] or treatment [*F*_(1, 34)_ = .27, *p* = .61] effects on the OFT anxiety index (**Figure 2C**). Post hoc testing revealed that WDN rats exhibited reduced indices of anxiety in the OFT relative to CDN rats (*p* = .019). These behavioral alterations were accompanied by a significant diet [*F*_(1, 34)_ = 5.70, *p* = .023], and no significant interaction [*F*_(1, 34)_ = .024, *p* = .88] or treatment [*F*_(1, 34)_ = .0007, *p* = .98] effects on the total distance traveled during the 5-min testing session (**Figure 2D**). Interestingly, when the rats were exposed to a more anxiogenic behavioral setting, the elevated-plus maze (EPM), the same groups showed opposite anxiety index results. Analyses revealed a significant diet [*F*_(1, 43)_ = 7.27, *p* = .009], and no significant interaction [*F*_(1, 43)_ = 1.38, *p* = .25] or treatment [*F*_(1, 43)_ = .066, *p* = .80] effects on the EPM anxiety index (**Figure 2E**). WDN rats demonstrated higher anxiety indices in the EPM when compared to CDN (*p* = .017). We used the number of closed arm entries as a proxy for locomotion in the EPM. We found no significant interaction [*F*_(1, 43)_ = .613, *p* = .438], treatment [*F*_(1, 43)_ = .434, *p* = .514], or diet [*F*_(1, 43)_ = .051, *p* = .822] effects in the number of closed arm entries (**Figure 2F**).

Spontaneous alternations in the Y-maze revealed no significant interaction [*F*_(1, 40)_ = .007, *p*= .93], treatment [*F*_(1, 40)_ = .97, *p*= .33], or diet [*F*_(1, 40)_ = .011, *p* = .92] effects (**Figure 2G**). Analyses for the total number of alternations revealed no significant interaction [*F*_(1, 40)_ = 1.00, *p*= .32], treatment [*F*_(1, 40)_ = .32, *p*= .58], or diet [*F*_(1, 40)_ = .22, *p* = .64] effects (**Figure 2H**).

### Exogenous subchronic NRG1 reduces the hippocampal volume in adolescent rats

Exogenous NRG1 confers robust anti-inflammatory (Li et al., 2015; Liu et al., 2018), neuroprotective (Li et al., 2012; Depboylu et al., 2015; Guan et al., 2015), and neurogenic effects (Mahar et al., 2011, 2016). In this respect, we asked whether NRG1 administration prevents hippocampal atrophy in rats that consume obesogenic diets. We examined hippocampal volumetric measurements using magnetic resonance imaging (MRI). We found a significant interaction [*F*_(1, 19)_ = 6.61, *p* = .019] and treatment [*F*_(1, 19)_ = 9.48, *p* = .006] effects on total hippocampal volume (both left and right hippocampi combined), with no diet [*F*_(1, 19)_ = .052, *p* = .82] effects (**Figure 3A-B**). The rats that received prolonged NRG1 administration during early adolescence exhibited smaller hippocampi (CDV vs. CDN, *p* = .003), an effect that was dependent on the diet type. The volumetric differences were not related to changes in total brain volume. Analyses revealed no significant interaction [*F*_(1, 19)_ = .14, *p* = .71], treatment [*F*_(1, 19)_ = .18, *p* = .67], or diet [*F*_(1, 19)_ = .25, *p* = .62] effects on total brain volume (**Supplemental Figure 2A**).

**Figure 3.**
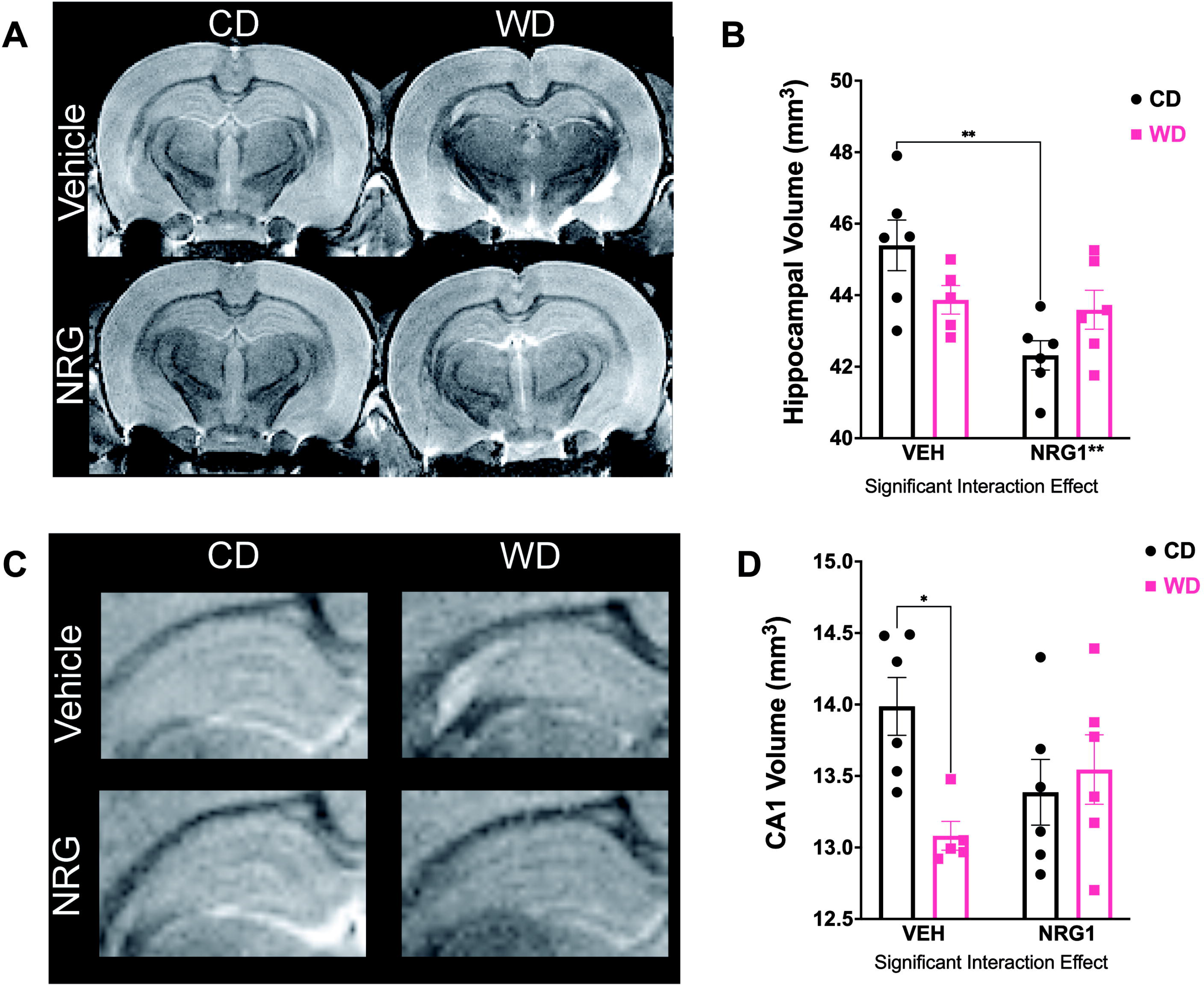
Obesogenic diet and prolonged NRG1 administration interact to reduce hippocampal volumes. **(A-B)** Total hippocampal volume (both left and right hippocampi combined) were affected by the treatment [*F*_(1, 19)_ = 9.48, *p* = .006]. Subchronic NRG1 administration during adolescence resulted in significant total hippocampal volume reduction (CDV vs. CDN, *p* = .003). **(C-D)** Hippocampal subfield segmentation showed a significant interaction [*F*_(1, 19)_ = 6.44, *p* = .020] effects on CA1 region. WDV rats had a significantly smaller CA1 volume than CDV rats (*p* = .015). Sample size = 6 rats / group.

Hippocampal subfield segmentation revealed unique vulnerabilities to the experimental manipulations in the CA1 region. Analyses revealed a significant interaction [*F*_(1, 19)_ = 6.44, *p* = .020] and no significant treatment [*F*_(1, 19)_ = .11, *p* = .75] or diet [*F*_(1, 19)_ = 3.16, *p* = .092] effects (**Figure 3C-D**), indicating a crossover interaction between the main factors. WDV rats had a significantly smaller CA1 volume than CDV rats (*p* = .035). **Table 2** details hippocampal volumetric data and statistics. The left and right rodent hippocampi exhibit striking anatomical and functional lateralization. We dissected the effects of the WD and hippocampal hemispheric lateralization. We found significant hippocampal hemisphere lateralization [*F*_(1, 19)_ = 6.08, *p* = .023] and diet type [*F*_(1, 19)_ = 4.71, *p* = .043] effects, with no significant interaction [*F*_(1, 19)_ = .053, *p* = .82] effects on hippocampal volumetrics (**Supplemental Figure 2B**). WD rats exhibited a reduced right hippocampal volume relative to the left hippocampal volume in CD rats (post hoc *p* = .019).

**Table 2.**
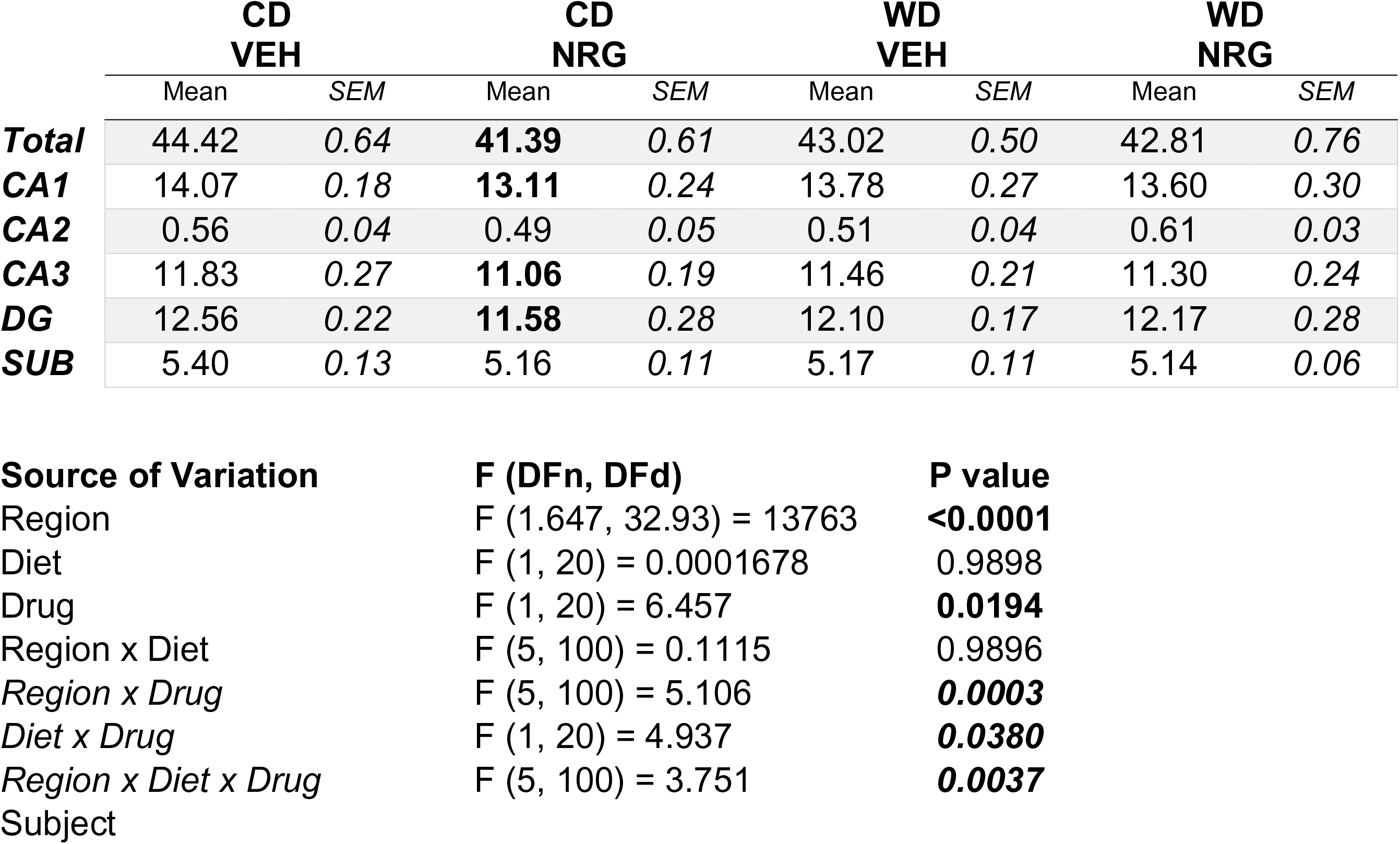
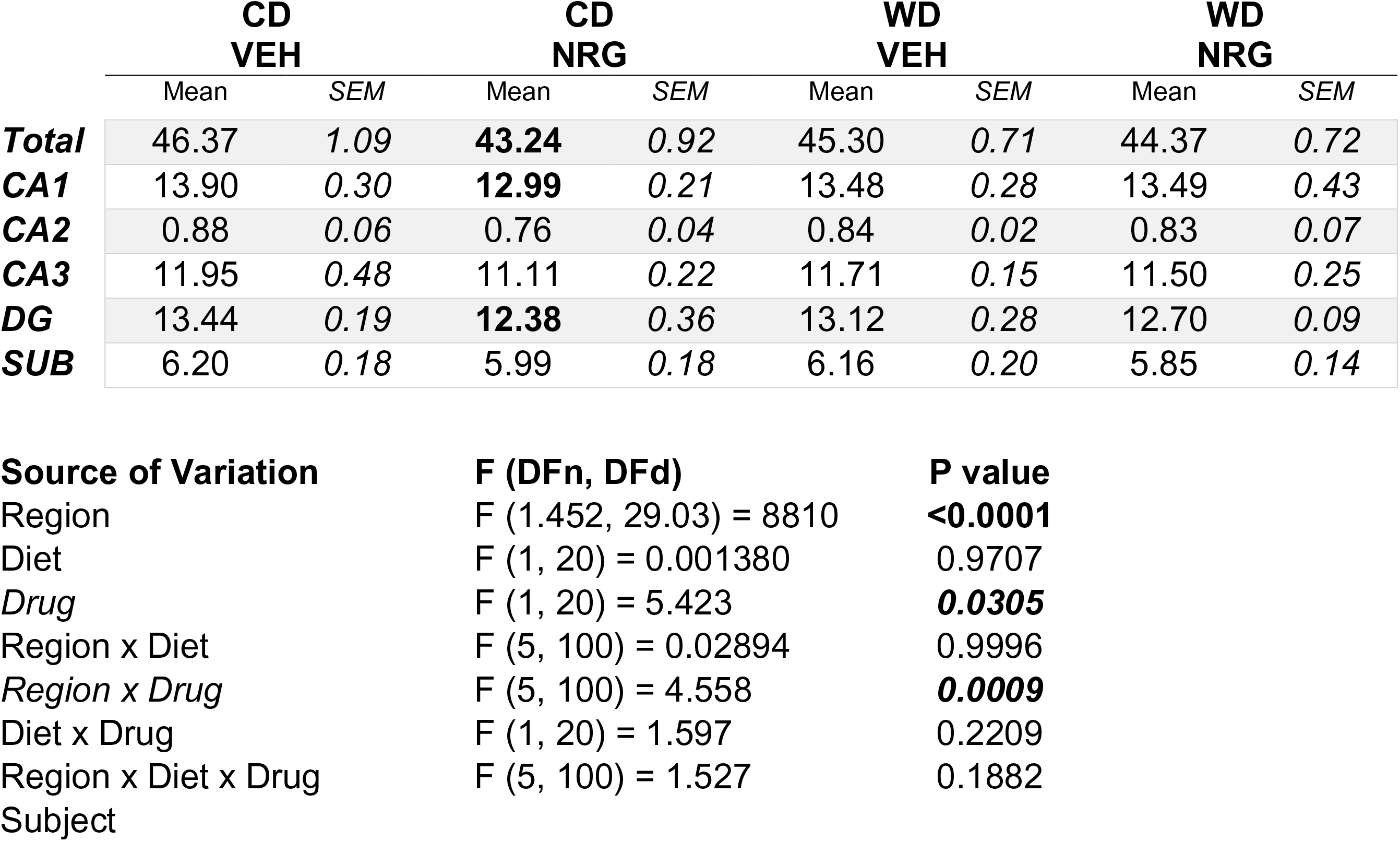
Detailed hippocampal subfield volumetrics and statistics. (A) Right hippocampal subfield volumes and three-way ANOVA statistics. **(B)** Left hippocampal subfield volumes and three-way ANOVA statistics.

#### Quantitative morphometric analysis shows a diverse morphologic microglia response to an obesogenic diet and exogenous NRG1 treatment in the CA1 region of the hippocampus

Microglia are the innate immune cells of the brain. Notably, microglia have been shown to play a significant role in axonal remodeling and synaptic pruning (Schafer and Stevens, 2013) and hippocampal tissue volume (Nelson et al., 2021). Microglia are highly responsive to obesogenic diets (Hao et al., 2016; Cope et al., 2018) and NRG1 (Simmons et al., 2016; Alizadeh et al., 2017). Given that microglial function is closely related to its morphology (Fernández-Arjona et al., 2017), we measured an array of 18 key morphometric parameters using fractal analysis (**Supplemental Table 2**). We focused on the CA1 hippocampal subfield based on its role on hippocampal-dependent behavior, abundant ErbB4 expression, and the observed volumetric changes and vulnerabilities of this region to several environmental insults. We performed morphological analyses in 1,981 Iba-1-positive cells (approximately 500 microglia/group). We applied Principal Component (PC) Analysis on the morphometric parameters to trace the possible differences in microglia driving hippocampal structure and behavior changes. PCs with eigenvalues greater than 1 were used (*Kaiser* rule). For all groups, we had three PCs with eigenvalues greater than 1. These PCs explained more than 80% of the accumulated variance between cells (PC1: ∼60%; PC2: ∼20%; PC3: ∼5%; **Supplemental Figure 4**). The analyses allowed for the extraction of presumably independent (non-overlapping) factors reflecting different parameters constituting microglial shape and function in our rat model. We identified several morphometric features with variable contribution values higher than .06 (1/18; 18 total number of features). These morphometric parameters were subjected to PCA. PCA revealed distinctive morphological profiles between groups; PC1 contributed to explaining diet effects, and PC2 helped understand the differences between NRG1-treated groups (PC1 = 58, PC2 = 35%; **Supplemental Figure 5**). **Figure 4A** shows representative microglia morphologies from hippocampal dorsal and ventral CA1 regions in each study group. Evaluation of microglial density (**Figure 4B**) as a measure of cell size based on area demonstrated that the WD significantly decreased microglial density in the right CA1 subfield (diet: [*F*_(1,8)_ = 6.82, *p* < .031]; complete *F* stats summarized in **Table 5**). This finding indicates an amoeboid and thus phagocytic microglia phenotype, supporting the reduced CA1 volume and the increased anxiety-like behaviors in the EPM in WD rats. Microglial lacunarity is a measure of heterogeneity that reflects sensitive changes in particular features such as soma size relative to process length (Karperien and Jelinek, 2015). We found that lacunarity was significantly decreased in the right CA1 subfield of rats that consumed an obesogenic WD (**Figure 4C**) (*diet:* [*F*_(1,8)_ = 12.09, *p* <  .008]; complete *F* stats summarized in **Table 5**), suggesting lower complexity with increased microglial transformation (activation).

**Figure 4.**
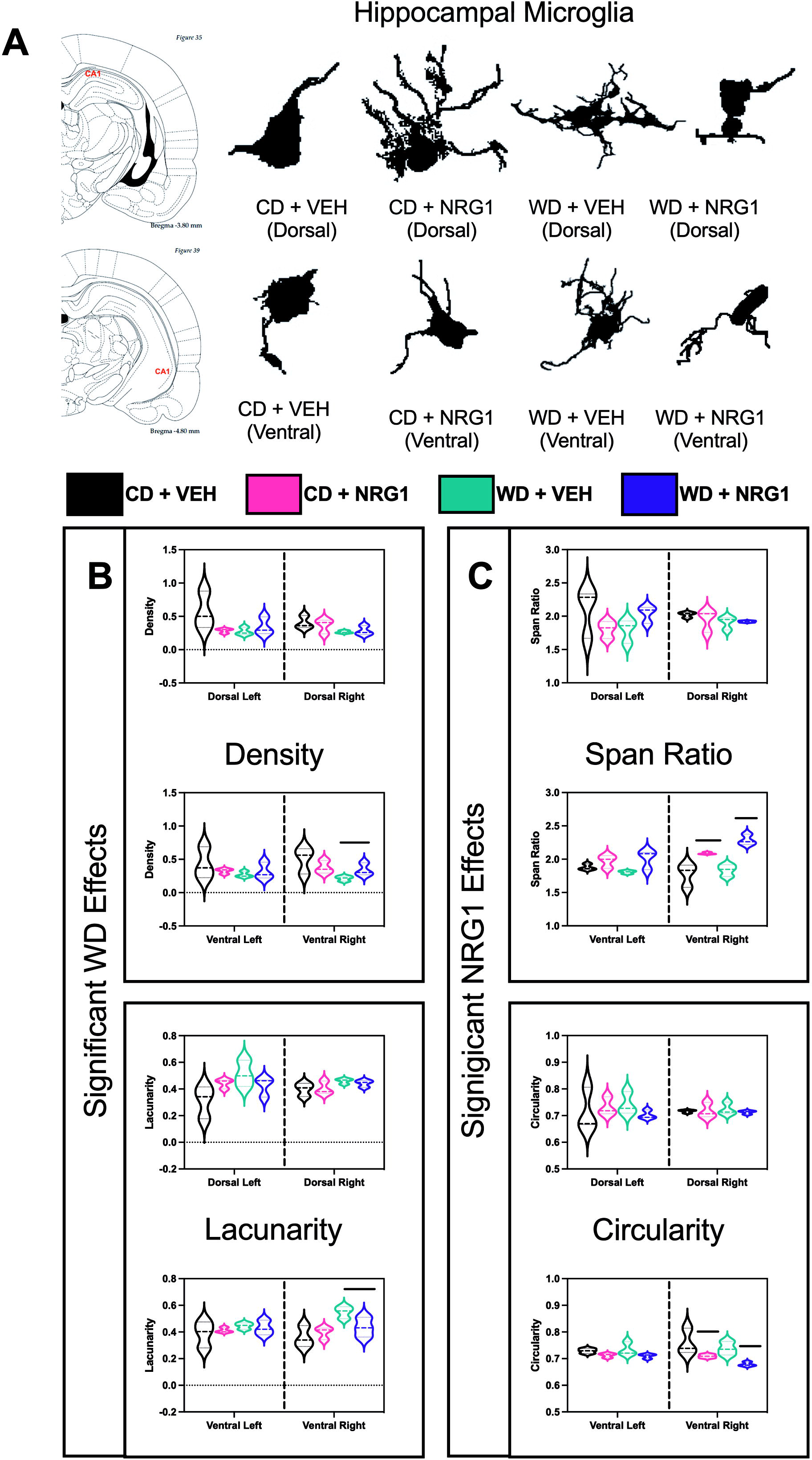
Obesogenic diet and exogenous NRG1 injections alter microglial shape in the hippocampus. Full-focused composite 40x images were acquired from twelve (12) rats. Hippocampal dorsal and ventral CA1 areas were scanned to obtain at least three (3) images per region of interest per hemisphere covering a field of view area of 1.2 mm. Each acquired image was saved as a TIFF file and included at least six (6) Iba-1^+^ cells. **(A)** Representative microglia morphologies from hippocampal dorsal and ventral CA1 regions in each study group. **(B)** Evaluation of microglial density as a measure of cell size based on area demonstrated that the WD significantly decreased microglial density in the right CA1 subfield (diet: [*F*_(1,8)_ = 6.82, *p* < .031]). **(C)** Lacunarity was significantly decreased in the right CA1 subfield of rats that consumed an obesogenic WD (*diet:* [*F*_(1,8)_ = 12.09, *p* <  .008]). The span ratio was significantly increased in the right CA1 region of rats receiving the exogenous NRG1 (*treatment:* [*F*_(1,8)_ = 15.43, *p* < .004]). NRG1 administration reduced circularity in the right CA1 region (*treatment:* [*F*_(1,8)_ = 9.67, *p* < .014]). Sample size = 3 rats / group.

The span ratio is a measure that describes the cell shape (or form factor) and is based on the ratio of length and height of the Convex Hull area occupied by the cell (Fernández-Arjona et al., 2017). The higher the span ratio, the less difference in area between soma and processes, suggesting a more amoeboid shape. The span ratio was significantly increased in the right CA1 region of rats receiving the exogenous NRG1 (**Figure 4C**) (*treatment:* [*F*_(1,8)_ = 15.43, *p* < .004]; complete *F* stats summarized in **Table 5**). Circularity is a measure of shape and size. High circularity values are associated with round sphere shapes, suggesting a more ameboid-like morphology associated with a greater phagocytic activity (Fernández-Arjona et al., 2017). NRG1 administration reduced circularity in the right CA1 region (**Figure 4C**) (*treatment:* [*F*_(1,8)_ = 9.67, *p* < .014]), complete *F* stats summarized in **Table 5**). Detailed microglial parameter values were reported as mean ± SEM for each group (**Table 4**). Complete details regarding microglial morphology descriptors statistics can be found in **Table 5**.

### WD consumption increases cytokine levels while exogenous NRG1 administration decreases selective cytokine levels in the rat hippocampus

Obesity is associated with a robust neuroinflammatory state via increased inflammatory mediators. Our data revealing diet-induced changes in microglial morphology prompted us to examine hippocampal cytokine levels. Using a bead-based flow cytometry immunoassay, we found that the WD exerted a precise cytokine regulation (**Figure 5**). The rats that consumed the WD exhibited a significant increase in pro-inflammatory cytokines in the hippocampus: IL1-α (*p* = .010), TNF-α (*p* = .006), and IL-6 (*p* = .019). Interestingly, the anti-inflammatory cytokine, IL-10, was also significantly increased in WD rats (*p* = .027). Opposite to the main dietary effects in increasing hippocampal cytokine levels, exogenous NRG1 administration significantly reduced IL-33 (*p* = .041), GM-CSF (*p* = .021), CCL-2 (*p* = .037), and IFN-γ (*p* = .031), confirming the potent immunomodulatory effects of NRG1 (Li et al., 2015; Simmons et al., 2016). **Table 3** includes the results and statistics of the cytokines tested in this study.

**Figure 5.**
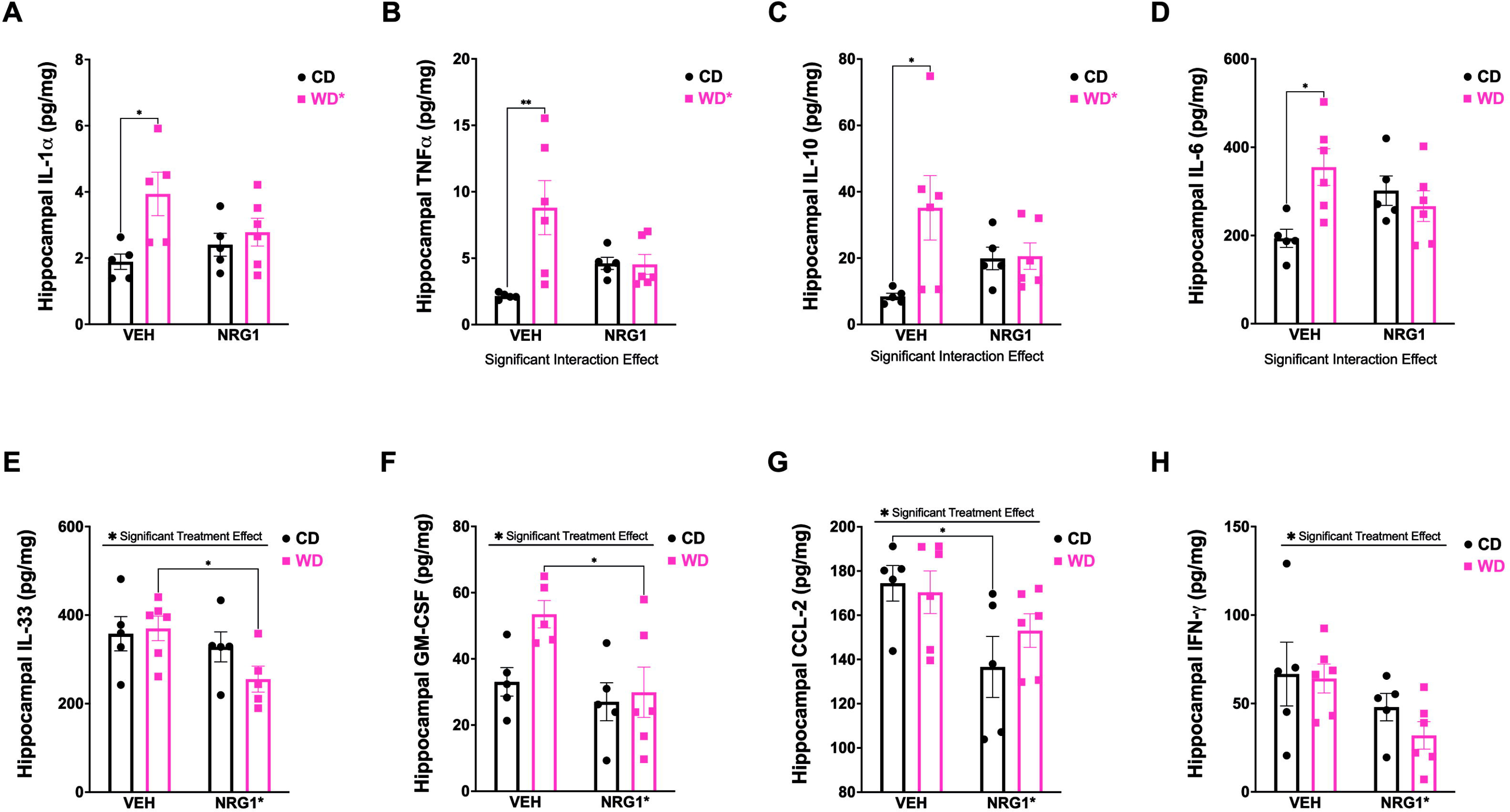
Obesogenic diet consumption and prolonged NRG1 injections alter selective inflammatory mediators in the rat hippocampus. Bead-based flow cytometry immunoassay was used to evaluate cytokine levels. **(A-D)** The WD led to a significant increase in cytokines in the hippocampus: IL1-α (*p* = .010), TNF-α (*p* = .006), IL-10 (*p* = .027), and IL-6 (*p* = .019). **(E-H)** NRG1 administration significantly reduced IL-33 (*p* = .041), GM-CSF (*p* = .021), CCL-2 (*p* = .037), and IFN-γ (*p* = .031). Table 3 includes the detailed statistics of the cytokines listed in Figure 5. Sample size = 5-6 rats / group.

**Table 3.** Detailed cytokine profiles and related statistics.

| Cytokines | Mean ± Standard Error of the Mean (pg/mg) |  |  |  | F | p |
| --- | --- | --- | --- | --- | --- | --- |
|  | Control Diet |  | Western Diet |  |  |  |
|  | Vehicle | NRG-1 | Vehicle | NRG-1 |  |  |
| IL-10 | 8.44 ± 0.96 | 19.90 ± 3.38 | 35.14 ± 9.71 | 20.56 ± 3.99 | Interaction, 4.683 | **Interaction, 0.0441** |
|  |  |  |  |  | Drug, 0.06692 | Drug, 0.7988 |
|  |  |  |  |  | Diet, 5.171 | **Diet, 0.0354** |
| TNF-α | 2.15 ± 0.10 | 4.61 ± 0.46 | 8.81 ± 2.04 | 4.54 ± 0.75 | Interaction, 7.620 | **Interaction, 0.0129** |
|  |  |  |  |  | Drug, 0.5545 | Drug, 0.4661 |
|  |  |  |  |  | Diet, 7.285 | **Diet, 0.0147** |
| IL-1α | 1.89 ± 0.23 | 2.40 ± 0.35 | 3.94 ± 0.66 | 2.78 ± 0.42 | Interaction, 3.542 | Interaction, 0.0771 |
|  |  |  |  |  | Drug, 0.5266 | Drug, 0.4779 |
|  |  |  |  |  | Diet, 7.502 | **Diet, 0.0140** |
| IL-1β | 26.04 ± 3.51 | 33.64 ± 2.17 | 51.36 ± 11.07 | 36.74 ± 6.16 | Interaction, 2.351 | Interaction, 0.1426 |
|  |  |  |  |  | Drug, 0.2340 | Drug, 0.6344 |
|  |  |  |  |  | Diet, 3.846 | Diet, 0.0655 |
| IL-6 | 193.63 ± 20.64 | 301.67 ± 33.35 | 354.98 ± 41.54 | 266.54 ± 34.69 | Interaction, 7.988 | **Interaction, 0.0112** |
|  |  |  |  |  | Drug, 0.07946 | Drug, 0.7812 |
|  |  |  |  |  | Diet, 3.297 | Diet, 0.0861 |
| IFN-γ | 66.65 ± 18.03 | 47.93 ± 7.76 | 64.12 ± 8.19 | 31.96 ± 7.76 | Interaction, 0.3843 | Interaction, 0.5431 |
|  |  |  |  |  | Drug, 5.514 | **Drug, 0.0305** |
|  |  |  |  |  | Diet, 0.7291 | Diet, 0.4044 |
| CXCL1 | 152.59 ± 3.88 | 145.80 ± 9.21 | 165.27 ± 16.81 | 126.12 ± 11.40 | Interaction, 1.788 | Interaction, 0.1978 |
|  |  |  |  |  | Drug, 3.603 | Drug, 0.0738 |
|  |  |  |  |  | Diet, 0.08352 | Diet, 0.7759 |
| CCL-2 | 174.48 ± 8.05 | 136.63 ± 13.80 | 170.40 ± 9.62 | 153.06 ± 7.60 | Interaction, 1.074 | Interaction, 0.3137 |
|  |  |  |  |  | Drug, 7.773 | **Drug, 0.0121** |
|  |  |  |  |  | Diet, 0.3892 | Diet, 0.5405 |
| GM-CSF | 33.02 ± 4.32 | 27.02 ± 5.73 | 53.48 ± 4.13 | 29.90 ± 7.60 | Interaction, 2.209 | Interaction, 0.1555 |
|  |  |  |  |  | Drug, 6.250 | **Drug, 0.0229** |
|  |  |  |  |  | Diet, 3.888 | Diet, 0.0651 |
| IL-18 | 1,056.79 ± 57.36 | 1,074.40 ± 68.86 | 1,102.42 ± 36.09 | 1,106.25 ± 60.97 | Interaction, 0.01497 | Interaction, 0.9040 |
|  |  |  |  |  | Drug, 0.03624 | Drug, 0.8512 |
|  |  |  |  |  | Diet, 0.4736 | Diet, 0.5001 |
| IL-12p70 | 82.11 ± 4.80 | 91.07 ± 7.73 | 105.17 ± 10.55 | 88.31 ± 14.19 | Interaction, 1.482 | Interaction, 0.2392 |
|  |  |  |  |  | Drug, 0.1385 | Drug, 0.7141 |
|  |  |  |  |  | Diet, 0.9161 | Diet, 0.3512 |
| IL-17a | 18.83 ± 0.54 | 19.17 ± 1.54 | 20.69 ± 1.96 | 15.69 ± 2.39 | Interaction, 2.062 | Interaction, 0.1682 |
|  |  |  |  |  | Drug, 1.570 | Drug, 0.2263 |
|  |  |  |  |  | Diet, 0.1897 | Diet, 0.6683 |
| IL-33 | 357.80 ± 38.71 | 328.15 ± 33.89 | 370.01 ± 27.68 | 255.37 ± 29.44 | Interaction, 1.717 | Interaction, 0.2074 |
|  |  |  |  |  | Drug, 4.950 | **Drug, 0.0399** |
|  |  |  |  |  | Diet, 0.8724 | Diet, 0.3634 |

**Table 4.** Detailed microglial morphometric analyses values.

| Parameters | Mean ± Standard Error of the Mean |  |  |  |  |  |  |  |  |  |  |  |  |  |  |  |
| --- | --- | --- | --- | --- | --- | --- | --- | --- | --- | --- | --- | --- | --- | --- | --- | --- |
|  | Control Diet |  |  |  |  |  |  |  | Western Diet |  |  |  |  |  |  |  |
|  | Vehicle |  |  |  | NRG-1 |  |  |  | Vehicle |  |  |  | NRG-1 |  |  |  |
|  | Left |  | Right |  | Left |  | Right |  | Left |  | Right |  | Left |  | Right |  |
|  | Dorsal | Ventral | Dorsal | Ventral | Dorsal | Ventral | Dorsal | Ventral | Dorsal | Ventral | Dorsal | Ventral | Dorsal | Ventral | Dorsal | Ventral |
| Objects | 12.11 ± 0.56 | 13.22 ± 3.11 | 12.44 ± 3.18 | 11.89 ± 4.59 | 16.00 ± 3.61 | 15.00 ± 0.51 | 13.22 ± 1.68 | 12.22 ± 0.62 | 14.22 ± 0.29 | 14.56 ± 1.64 | 17.22 ± 1.49 | 15.44 ± 0.87 | 15.22 ± 2.04 | 17.22 ± 3.35 | 10.22 ± 2.25 | 10.44 ± 0.62 |
| Density | 0.57 ± 0.16 | 0.43 ± 0.14 | 0.40 ± 0.06 | 0.50 ± 0.11 | 0.29 ± 0.02 | 0.32 ± 0.02 | 0.36 ± 0.06 | 0.38 ± 0.06 | 0.27 ± 0.03 | 0.27 ± 0.03 | 0.26 ± 0.01 | 0.21 ± 0.02 | 0.34 ± 0.08 | 0.32 ± 0.07 | 0.28 ± 0.04 | 0.34 ± 0.06 |
| Span Ratio | 2.10 ± 0.21 | 1.88 ± 0.03 | 2.02 ± 0.02 | 1.77 ± 0.10 | 1.80 ± 0.07 | 1.97 ± 0.06 | 1.95 ± 0.10 | 2.09 ± 0.01 | 1.79 ± 0.10 | 1.80 ± 0.01 | 1.91 ± 0.06 | 1.83 ± 0.05 | 2.04 ± 0.07 | 2.01 ± 0.08 | 1.92 ± 0.01 | 2.29 ± 0.05 |
| Maximum Span Across Hull | 86.21 ± 23.79 | 114.09 ± 33.30 | 115.05 ± 16.39 | 94.37 ± 30.57 | 151.39 ± 20.61 | 138.34 ± 18.78 | 130.72 ± 32.07 | 118.28 ± 23.03 | 165.78 ± 30.38 | 159.78 ± 14.18 | 151.94 ± 3.69 | 174.55 ± 14.69 | 131.27 ± 28.21 | 134.50 ± 22.29 | 144.76 ± 14.01 | 130.19 ± 18.35 |
| Convex Hull Area | 3,366 ± 1,658 | 6,443 ± 3,530 | 5,707 ± 1,530 | 4,598 ± 2,894 | 10,482 ± 3,564 | 8,304 ± 2,481 | 8,237 ± 4,523 | 6,366 ± 2,358 | 13,321 ± 5,130 | 12,095 ± 2,678 | 10,262 ± 1,195 | 13,412 ± 2,377 | 7,597 ± 3,189 | 7,566 ± 2,192 | 8,343 ± 1,304 | 6,478 ± 1,616 |
| Perimeter | 212.76 ± 57.46 | 286.98 ± 86.04 | 285.26 ± 41.98 | 239.02 ± 77.45 | 384.77 ± 56.46 | 347.12 ± 47.86 | 328.20 ± 86.28 | 294.37 ± 59.18 | 423.51 ± 81.64 | 407.88 ± 38.47 | 385.42 ± 15.07 | 444.27 ± 42.51 | 326.49 ± 73.00 | 335.30 ± 55.93 | 362.21 ± 35.11 | 316.88 ± 44.33 |
| Circularity | 0.714 ± 0.046 | 0.729 ± 0.005 | 0.716 ± 0.003 | 0.759 ± 0.028 | 0.732 ± 0.020 | 0.713 ± 0.005 | 0.717 ± 0.017 | 0.709 ± 0.004 | 0.742 ± 0.025 | 0.735 ± 0.014 | 0.724 ± 0.013 | 0.738 ± 0.015 | 0.703 ± 0.009 | 0.710 ± 0.005 | 0.712 ± 0.004 | 0.679 ± 0.005 |
| Width of Bounding Rectangle | 70.44 ± 19.03 | 89.10 ± 20.98 | 92.09 ± 13.91 | 71.11 ± 24.65 | 122.23 ± 19.01 | 105.38 ± 14.93 | 107.02 ± 28.05 | 93.41 ± 19.36 | 137.67 ± 27.18 | 126.62 ± 12.79 | 118.88 ± 3.93 | 139.26 ± 10.98 | 101.24 ± 22.97 | 100.18 ± 13.13 | 110.54 ± 9.17 | 94.03 ± 10.09 |
| Height of Bounding Rectangle | 66.05 ± 17.26 | 95.11 ± 33.46 | 91.64 ± 13.17 | 82.32 ± 24.75 | 124.28 ± 16.62 | 116.00 ± 15.29 | 103.58 ± 27.18 | 93.51 ± 17.91 | 133.68 ± 26.34 | 133.54 ± 12.84 | 125.03 ± 7.37 | 144.28 ± 15.30 | 106.17 ± 23.40 | 112.77 ± 20.55 | 120.90 ± 13.66 | 108.77 ± 17.67 |
| Max Radius from Hull's Centre of Mass | 48.56 ± 11.78 | 62.77 ± 17.67 | 63.38 ± 9.09 | 52.45 ± 16.06 | 82.28 ± 10.78 | 75.39 ± 9.62 | 71.70 ± 16.86 | 64.15 ± 11.97 | 89.35 ± 15.57 | 86.65 ± 7.20 | 82.67 ± 2.05 | 94.41 ± 7.75 | 71.73 ± 15.01 | 73.53 ± 11.23 | 79.72 ± 7.89 | 71.11 ± 9.52 |
| Max/Min Radii | 2.60 ± 0.21 | 2.08 ± 0.13 | 2.20 ± 0.08 | 1.99 ± 0.09 | 1.94 ± 0.07 | 2.01 ± 0.03 | 2.03 ± 0.13 | 2.13 ± 0.04 | 1.88 ± 0.09 | 1.94 ± 0.07 | 1.94 ± 0.05 | 1.92 ± 0.13 | 2.02 ± 0.11 | 2.02 ± 0.06 | 2.05 ± 0.06 | 2.38 ± 0.09 |
| Mean Radius | 37.62 ± 9.97 | 50.09 ± 14.62 | 50.04 ± 7.47 | 41.26 ± 13.33 | 65.77 ± 9.33 | 60.49 ± 8.34 | 57.44 ± 14.70 | 51.40 ± 10.08 | 73.03 ± 13.38 | 70.67 ± 6.31 | 66.89 ± 2.28 | 76.49 ± 7.35 | 56.97 ± 12.48 | 58.92 ± 9.47 | 63.11 ± 6.16 | 56.15 ± 7.88 |
| Diameter of Bounding Circle | 86.82 ± 23.97 | 115.05 ± 33.67 | 115.75 ± 16.63 | 95.10 ± 30.98 | 152.96 ± 20.64 | 139.48 ± 18.92 | 131.68 ± 32.30 | 119.32 ± 23.33 | 167.14 ± 30.52 | 161.29 ± 14.16 | 153.11 ± 3.98 | 175.89 ± 15.06 | 132.44 ± 28.45 | 135.55 ± 22.43 | 145.73 ± 13.99 | 131.04 ± 18.30 |
| Maximum Radius | 43.41 ± 11.99 | 57.53 ± 16.84 | 57.87 ± 8.31 | 47.55 ± 15.49 | 76.48 ± 10.32 | 69.74 ± 9.46 | 65.84 ± 16.15 | 59.66 ± 11.66 | 83.57 ± 15.26 | 80.65 ± 7.08 | 76.56 ± 1.99 | 87.94 ± 7.53 | 66.22 ± 14.23 | 67.78 ± 11.21 | 72.87 ± 7.00 | 65.52 ± 9.15 |
| CV for all Radii from Circle's Centre | 0.17 ± 0.01 | 0.16 ± 0.01 | 0.18 ± 0.01 | 0.16 ± 0.00 | 0.17 ± 0.01 | 0.17 ± 0.00 | 0.16 ± 0.01 | 0.18 ± 0.00 | 0.15 ± 0.01 | 0.15 ± 0.01 | 0.16 ± 0.01 | 0.17 ± 0.02 | 0.17 ± 0.01 | 0.16 ± 0.01 | 0.17 ± 0.01 | 0.19 ± 0.01 |
| CV for all Radii | 0.24 ± 0.02 | 0.21 ± 0.01 | 0.21 ± 0.01 | 0.20 ± 0.01 | 0.19 ± 0.01 | 0.19 ± 0.00 | 0.20 ± 0.02 | 0.21 ± 0.01 | 0.17 ± 0.01 | 0.18 ± 0.01 | 0.19 ± 0.01 | 0.18 ± 0.02 | 0.21 ± 0.01 | 0.19 ± 0.01 | 0.20 ± 0.01 | 0.23 ± 0.01 |
| Mean Radii | 37.85 ± 10.57 | 50.35 ± 14.90 | 50.31 ± 7.59 | 41.40 ± 13.53 | 66.43 ± 9.49 | 61.16 ± 8.40 | 57.94 ± 14.92 | 51.80 ± 10.36 | 73.69 ± 13.39 | 71.55 ± 6.44 | 67.83 ± 2.20 | 77.50 ± 7.46 | 57.66 ± 12.71 | 59.75 ± 9.79 | 64.08 ± 6.62 | 56.54 ± 8.13 |
| Lacunarity (Λ) | 0.31 ± 0.07 | 0.39 ± 0.06 | 0.40 ± 0.03 | 0.36 ± 0.05 | 0.44 ± 0.02 | 0.41 ± 0.01 | 0.40 ± 0.03 | 0.40 ± 0.03 | 0.51 ± 0.06 | 0.45 ± 0.01 | 0.46 ± 0.01 | 0.55 ± 0.03 | 0.42 ± 0.04 | 0.43 ± 0.03 | 0.44 ± 0.02 | 0.43 ± 0.04 |

**Table 5.** Detailed microglial morphology descriptors statistics.

| Parameters | *F* |  | *p* |  |
| --- | --- | --- | --- | --- |
|  | Left | Right | Left | Right |
| Objects | ROI, 0.2742<br>Diet, 0.3921<br>Drug, 1.429<br>ROI x Diet, 0.2266<br>ROI x Drug, 0.009065<br>Diet x Drug, 0.06563<br>ROI x Diet x Drug, 0.6550 | ROI, 0.5444<br>Diet, 0.1864<br>Drug, 1.748<br>ROI x Diet, 1.000e-013<br>ROI x Drug, 0.1361<br>Diet x Drug, 2.534<br>ROI x Diet x Drug, 0.3361 | ROI, 0.6147<br>Diet, 0.5486<br>Drug, 0.2661<br>ROI x Diet, 0.6468<br>ROI x Drug, 0.9265<br>Diet x Drug, 0.8043<br>ROI x Diet x Drug, 0.4417 | ROI, 0.4817<br>Diet, 0.6773<br>Drug, 0.2227<br>ROI x Diet, >0.9999<br>ROI x Drug, 0.7218<br>Diet x Drug, 0.1501<br>ROI x Diet x Drug, 0.5780 |
| Density | ROI, 3.696<br>Diet, 1.474<br>Drug, 0.6682<br>ROI x Diet, 1.289<br>ROI x Drug, 5.310<br>Diet x Drug, 2.251<br>ROI x Diet x Drug, 9.357 | ROI, 1.061<br>Diet, 6.824<br>Drug, 008865<br>ROI x Diet, 0.8167<br>ROI x Drug, .04138<br>Diet x Drug, 2.173<br>ROI x Diet x Drug, 2.706 | **ROI, 0.0908**<br>Diet, 0.2593<br>Drug, 0.4374<br>ROI x Diet, 0.2892<br>ROI x Drug, 0.0501<br>Diet x Drug, 0.1720<br>**ROI x Diet x Drug, 0.0156** | ROI, 0.3332<br>Diet, 0.0310<br>Drug, 0.9273<br>ROI x Diet, 0.3926<br>ROI x Drug, 0.8439<br>Diet x Drug, 0.1787<br>ROI x Diet x Drug, 0.1386 |
| Span Ratio | ROI, 0.04611<br>Diet, 0.1785<br>Drug, 1.007<br>ROI x Diet, 0.009374<br>ROI x Drug, 1.260<br>Diet x Drug, 6.741<br>ROI x Diet x Drug, 1.952 | ROI, 1.429<br>Diet, 0.4127<br>Drug, 15.43<br>ROI x Diet, 6.875<br>ROI x Drug, 29.31<br>Diet x Drug, 1.631<br>ROI x Diet x Drug, 0.1072 | ROI, 0.8353<br>Diet, 0.6838<br>Drug, 0.3449<br>ROI x Diet, 0.9252<br>ROI x Drug, 0.2942<br>Diet x Drug, 0.0318<br>ROI x Diet x Drug, 0.1999 | ROI, 0.2662<br>Diet, 0.5386<br>Drug, 0.0044<br>**ROI x Diet, 0.0306**<br>**ROI x Drug, 0.0006**<br>Diet x Drug, 0.2373<br>ROI x Diet x Drug, 0.7517 |
| Maximum Span Across Hull | ROI, 0.1980<br>Diet, 1.139<br>Drug, 0.09756<br>ROI x Diet, 0.4224<br>ROI x Drug, 1.372<br>Diet x Drug, 2.472<br>ROI x Diet x Drug, 3.431 | ROI, 0.3674<br>Diet, 3.832<br>Drug, 0.02684<br>ROI x Diet, 0.9901<br>ROI x Drug, 0.4896<br>Diet x Drug, 1.556<br>ROI x Diet x Drug, 1.207 | ROI, 0.6682<br>Diet, 0.3169<br>Drug, 0.7628<br>ROI x Diet, 0.5340<br>ROI x Drug, 0.2752<br>Diet x Drug, 0.1545<br>ROI x Diet x Drug, 0.1011 | ROI, 0.5612<br>Diet, 0.0860<br>Drug, 0.8739<br>ROI x Diet, 0.3489<br>ROI x Drug, 0.5039<br>Diet x Drug, 0.2476<br>ROI x Diet x Drug, 0.3039 |
| Convex Hull Area | ROI, 0.006925<br>Diet, 0.9803<br>Drug, 0.01110<br>ROI x Diet, 0.2508<br>ROI x Drug, 0.8878<br>Diet x Drug, 2.524<br>ROI x Diet x Drug, 2.241 | ROI, 0.1278<br>Diet, 2.501<br>Drug, 0.2810<br>ROI x Diet, 0.8100<br>ROI x Drug, 1.486<br>Diet x Drug, 2.343<br>ROI x Diet x Drug, 0.8057 | ROI, 0.9357<br>Diet, 0.3511<br>Drug, 0.9187<br>ROI x Diet, 0.6300<br>ROI x Drug, 0.3737<br>Diet x Drug, 0.1508<br>ROI x Diet x Drug, 0.1728 | ROI, 0.7299<br>Diet, 0.1524<br>Drug, 0.6104<br>ROI x Diet, 0.3944<br>ROI x Drug, 0.2575<br>Diet x Drug, 0.1644<br>ROI x Diet x Drug, 0.3956 |
| Perimeter | ROI, 0.1570<br>Diet, 1.142<br>Drug, 0.06528<br>ROI x Diet, 0.3338<br>ROI x Drug, 1.356<br>Diet x Drug, 2.693<br>ROI x Diet x Drug, 3.294 | ROI, 0.4019<br>Diet, 3.555<br>Drug, 0.07430<br>ROI x Diet, 0.7945<br>ROI x Drug, 0.7639<br>Diet x Drug, 1.681<br>ROI x Diet x Drug, 1.233 | ROI, 0.7023<br>Diet, 0.3165<br>Drug, 0.8048<br>ROI x Diet, 0.5793<br>ROI x Drug, 0.2778<br>Diet x Drug, 0.1394<br>ROI x Diet x Drug, 0.1071 | ROI, 0.5438<br>Diet, 0.0961<br>Drug, 0.7921<br>ROI x Diet, 0.3987<br>ROI x Drug, 0.4076<br>Diet x Drug, 0.2309<br>ROI x Diet x Drug, 0.2991 |
| **Circularity** | ROI, 0.01093<br>Diet, 0.0005047<br>Drug, 0.9625<br>ROI x Diet, 0.005566<br>ROI x Drug, 0.1345<br>Diet x Drug, 1.032<br>ROI x Diet x Drug, 0.8468 | ROI, 0.1408<br>Diet, 1.621<br>Drug, 9.667<br>ROI x Diet, 1.808<br>ROI x Drug, 6.005<br>Diet x Drug, 0.3333<br>ROI x Diet x Drug, 0.007073 | ROI, 0.9193<br>Diet, 0.9826<br>Drug, 0.3553<br>ROI x Diet, 0.9424<br>ROI x Drug, 0.7233<br>Diet x Drug, 0.3394<br>ROI x Diet x Drug, 0.3844 | ROI, 0.7173<br>Diet, 0.2386<br>Drug, 0.0145<br>ROI x Diet, 0.2156<br>**ROI x Drug, 0.0399**<br>Diet x Drug, 0.5796<br>ROI x Diet x Drug, 0.9350 |
| **Width of Bounding Rectangle** | ROI, 0.1934<br>Diet, 1.136<br>Drug, 0.004988<br>ROI x Diet, 0.3551<br>ROI x Drug, 1.192<br>Diet x Drug, 3.156<br>ROI x Diet x Drug, 3.784 | ROI, 0.9153<br>Diet, 2.774<br>Drug, 0.07531<br>ROI x Diet, 1.434<br>ROI x Drug, 0.8459<br>Diet x Drug, 2.330<br>ROI x Diet x Drug, 1.900 | ROI, 0.6717<br>Diet, 0.3176<br>Drug, 0.9454<br>ROI x Diet, 0.5677<br>ROI x Drug, 0.3067<br>Diet x Drug, 0.1135<br>ROI x Diet x Drug, 0.0876 | ROI, 0.3667<br>Diet, 0.1344<br>Drug, 0.7907<br>ROI x Diet, 0.2653<br>ROI x Drug, 0.3846<br>Diet x Drug, 0.1654<br>ROI x Diet x Drug, 0.2054 |
| **Height of Bounding Rectangle** | ROI, 0.9757<br>Diet, 1.063<br>Drug, 0.1408<br>ROI x Diet, 0.2694<br>ROI x Drug, 1.232<br>Diet x Drug, 2.404<br>ROI x Diet x Drug, 2.554 | ROI, 0.1285<br>Diet, 3.997<br>Drug, 0.06649<br>ROI x Diet, 0.6001<br>ROI x Drug, 0.8813<br>Diet x Drug, 0.9623<br>ROI x Diet x Drug, 0.8008 | ROI, 0.3522<br>Diet, 0.3327<br>Drug, 0.7172<br>ROI x Diet, 0.6178<br>ROI x Drug, 0.2993<br>Diet x Drug, 0.1597<br>ROI x Diet x Drug, 0.1487 | ROI, 0.7293<br>Diet, 0.0806<br>Drug, 0.8030<br>ROI x Diet, 0.4608<br>ROI x Drug, 0.3753<br>Diet x Drug, 0.3553<br>ROI x Diet x Drug, 0.3970 |
| **Max Radius from Hull's Centre of Mass** | ROI, 0.1926<br>Diet, 1.139<br>Drug, 0.1015<br>ROI x Diet, 0.3153<br>ROI x Drug, 1.286<br>Diet x Drug, 2.480<br>ROI x Diet x Drug, 3.061 | ROI, 0.4640<br>Diet, 3.977<br>Drug, 0.02660<br>ROI x Diet, 0.9183<br>ROI x Drug, 0.5654<br>Diet x Drug, 1.465<br>ROI x Diet x Drug, 1.107 | ROI, 0.6724<br>Diet, 0.3170<br>Drug, 0.7582<br>ROI x Diet, 0.5898<br>ROI x Drug, 0.2896<br>Diet x Drug, 0.1540<br>ROI x Diet x Drug, 0.1183 | ROI, 0.5150<br>Diet, 0.0812<br>Drug, 0.8745<br>ROI x Diet, 0.3660<br>ROI x Drug, 0.4736<br>Diet x Drug, 0.2607<br>ROI x Diet x Drug, 0.3234 |
| **Max/Min Radii** | ROI, 9.990<br>Diet, 3.613<br>Drug, 1.611<br>ROI x Diet, 16.13<br>ROI x Drug, 17.35<br>Diet x Drug, 5.397<br>ROI x Diet x Drug, 25.86 | ROI, 1.230<br>Diet, 0.04899<br>Drug, 3.173<br>ROI x Diet, 5.523<br>ROI x Drug, 13.04<br>Diet x Drug, 3.558<br>ROI x Diet x Drug, 0.03950 | ROI, 0.0134<br>Diet, 0.0938<br>Drug, 0.2400<br>**ROI x Diet, 0.0039**<br>**ROI x Drug, 0.0031**<br>**Diet x Drug, 0.0487**<br>**ROI x Diet x Drug, 0.0009** | ROI, 0.2997<br>Diet, 0.8304<br>Drug, 0.1127<br>**ROI x Diet, 0.0467**<br>**ROI x Drug, 0.0069**<br>**Diet x Drug, 0.0960**<br>ROI x Diet x Drug, 0.8474 |
| **Mean Radius** | ROI, 0.3036<br>Diet, 1.212<br>Drug, 0.06713<br>ROI x Diet, 0.3811<br>ROI x Drug, 1.194<br>Diet x Drug, 2.565<br>ROI x Diet x Drug, 3.219 | ROI, 0.4337<br>Diet, 3.620<br>Drug, .04011<br>ROI x Diet, 0.8918<br>ROI x Drug, 0.5587<br>Diet x Drug, 1.608<br>ROI x Diet x Drug, 1.089 | ROI, 0.5967<br>Diet, 0.3030<br>Drug, 0.8021<br>ROI x Diet, 0.5542<br>ROI x Drug, 0.3064<br>Diet x Drug, 0.1479<br>ROI x Diet x Drug, 0.1105 | ROI, 0.5287<br>Diet, 0.0936<br>Drug, 0.8463<br>ROI x Diet, 0.3726<br>ROI x Drug, 0.4762<br>Diet x Drug, 0.2404<br>ROI x Diet x Drug, 0.3272 |
| **Diameter of Bounding Circle** | ROI, 0.1894<br>Diet, 1.144<br>Drug, 0.09966<br>ROI x Diet, 0.4010<br>ROI x Drug, 1.406 | ROI, 0.3576<br>Diet, 3.796<br>Drug, 0.02675<br>ROI x Diet, 0.9714<br>ROI x Drug, 0.4902 | ROI, 0.6749<br>Diet, 0.3159<br>Drug, 0.7603<br>ROI x Diet, 0.5443<br>ROI x Drug, 0.2698 | ROI, 0.5664<br>Diet, 0.0872<br>Drug, 0.8741<br>ROI x Diet, 0.3532<br>ROI x Drug, 0.5037 |
|  | Diet x Drug, 2.502<br>ROI x Diet x Drug, 3.364 | Diet x Drug, 1.564<br>ROI x Diet x Drug, 1.204 | Diet x Drug, 0.1523<br>ROI x Diet x Drug, 0.1040 | Diet x Drug, 0.2464<br>ROI x Diet x Drug, 0.3044 |
| Maximum Radius | ROI, 0.1894<br>Diet, 1.144<br>Drug, 0.09966<br>ROI x Diet, 0.4010<br>ROI x Drug, 1.406<br>Diet x Drug, 2.502<br>ROI x Diet x Drug, 3.364 | ROI, 0.3576<br>Diet, 3.796<br>Drug, 0.02675<br>ROI x Diet, 0.9714<br>ROI x Drug, 0.4902<br>Diet x Drug, 1.564<br>ROI x Diet x Drug, 1.204 | ROI, 0.6749<br>Diet, 0.3159<br>Drug, 0.7603<br>ROI x Diet, 0.5443<br>ROI x Drug, 0.2698<br>Diet x Drug, 0.1523<br>ROI x Diet x Drug, 0.1040 | ROI, 0.5664<br>Diet, 0.0872<br>Drug, 0.8741<br>ROI x Diet, 0.3532<br>ROI x Drug, 0.5037<br>Diet x Drug, 0.2464<br>ROI x Diet x Drug, 0.3044 |
| CV for all Radii from Cicle's Centre | ROI, 1.051<br>Diet, 3.613<br>Drug, 2.735<br>ROI x Diet, 0.02936<br>ROI x Drug, 0.09261<br>Diet x Drug, 0.5020<br>ROI x Diet x Drug, 1.041 | ROI, 2.158<br>Diet, 0.02939<br>Drug, 0.8968<br>ROI x Diet, 3.574<br>ROI x Drug, 5.949<br>Diet x Drug, 1.527<br>ROI x Diet x Drug, 1.163 | ROI, 0.3353<br>Diet, 0.0939<br>Drug, 0.1368<br>ROI x Diet, 0.8682<br>ROI x Drug, 0.7687<br>Diet x Drug, 0.4987<br>ROI x Diet x Drug, 0.3375 | ROI, 0.1800<br>Diet, 0.8681<br>Drug, 0.3714<br>ROI x Diet, 0.0953<br>**ROI x Drug, 0.0406**<br>Diet x Drug, 0.2517<br>ROI x Diet x Drug, 0.3124 |
| CV for all Radii | ROI, 6.259<br>Diet, 2.403<br>Drug, 0.2050<br>ROI x Diet, 3.428<br>ROI x Drug, 2.438<br>Diet x Drug, 5.630<br>ROI x Diet x Drug, 16.18 | ROI, 1.334<br>Diet, 0.09211<br>Drug, 1.762<br>ROI x Diet, 1.339<br>ROI x Drug, 4.832<br>Diet x Drug, 2.111<br>ROI x Diet x Drug, 0.3416 | ROI, 0.0368<br>Diet, 0.1597<br>Drug, 0.6628<br>ROI x Diet, 0.1012<br>ROI x Drug, 0.1571<br>**Diet x Drug, 0.0450**<br>**ROI x Diet x Drug, 0.0038** | ROI, 0.2815<br>Diet, 0.7693<br>Drug, 0.2210<br>ROI x Diet, 0.2806<br>ROI x Drug, 0.0592<br>Diet x Drug, 0.1843<br>ROI x Diet x Drug, 0.5750 |
| Mean Radii | ROI, 0.3429<br>Diet, 1.223<br>Drug, 0.07455<br>ROI x Diet, 0.3532<br>ROI x Drug, 1.219<br>Diet x Drug, 2.518<br>ROI x Diet x Drug, 3.221 | ROI, 0.4625<br>Diet, 3.716<br>Drug, 0.03975<br>ROI x Diet, 0.8162<br>ROI x Drug, 0.5777<br>Diet x Drug, 1.631<br>ROI x Diet x Drug, 1.103 | ROI, 0.5743<br>Diet, 0.3009<br>Drug, 0.7917<br>ROI x Diet, 0.5687<br>ROI x Drug, 0.3017<br>Diet x Drug, 0.1512<br>ROI x Diet x Drug, 0.1104 | ROI, 0.5156<br>Diet, 0.0901<br>Drug, 0.8470<br>ROI x Diet, 0.3927<br>ROI x Drug, 0.4690<br>Diet x Drug, 0.2374<br>ROI x Diet x Drug, 0.3243 |
| Lacunarity ($\Lambda$) | ROI, 0.03150<br>Diet, 2.737<br>Drug, 0.1107<br>ROI x Diet, 1.871<br>ROI x Drug, 0.1787<br>Diet x Drug, 2.863<br>ROI x Diet x Drug, 5.398 | ROI, 0.2166<br>Diet, 12.09<br>Drug, 0.9348<br>ROI x Diet, 2.772<br>ROI x Drug, 0.7385<br>Diet x Drug, 3.302<br>ROI x Diet x Drug, 2.758 | ROI, 0.8635<br>Diet, 0.1367<br>Drug, 0.7479<br>ROI x Diet, 0.2085<br>ROI x Drug, 0.6837<br>Diet x Drug, 0.1291<br>**ROI x Diet x Drug, 0.0487** | ROI, 0.6541<br>Diet, 0.0084<br>Drug, 0.3619<br>ROI x Diet, 0.1345<br>ROI x Drug, 0.4152<br>Diet x Drug, 0.1067<br>ROI x Diet x Drug, 0.1353 |

### WD consumption and exogenous NRG1 synergize to enhance hippocampal ErbB4 phosphorylation and sheddase protein levels

Dysfunctional NRG1-ErbB4 signaling has been recently implicated in obesity and obesity-related complications (Ennequin et al., 2015a; Salinas et al., 2016; Zeng et al., 2018). In support of these reports, we found that the obesogenic WD decreased total ErbB4 protein levels in the hippocampus (**Figure 6A**). Analyses revealed a significant treatment [*F*_(1, 16)_ = 7.97, *p* = .012] and diet [*F*_(1, 16)_ = 7.34, *p* = .016] effect, while showing no significant interaction [*F*_(1, 16)_ = .96, *p* = .34] effects on ErbB4 protein levels. Post hoc testing revealed that WDN rats exhibited significantly lower ErbB4 protein levels when compared to CDV rats (*p* = .009). Notably, exogenous NRG1 administration caused a hyperphosphorylation of hippocampal ErbB4 (**Figure 6B**). Analyses revealed a significant treatment [*F*_(1, 18)_ = 5.82, *p* = .027] and diet [*F*_(1, 18)_ = 7.36, *p*= .014] effect, while no significant interaction [*F*_(1,18)_ = .13, *p* = .73] effects on protein levels. ErbB4 hyperphosphorylation was particularly evident in WDN rats relative to the CDV group (*p* = .012). Immunohistological evaluation demonstrated pErbB4 expression in hippocampal Cd11b/c cells from rats treated with exogenous NRG1 (**Supplemental Figure 3**). Supporting the expression and roles for NRG1-ErbB4 signaling in microglia cell function (Gerecke et al., 2001; Mencel et al., 2013; Simmons et al., 2016; Chen et al., 2019).

**Figure 6.**
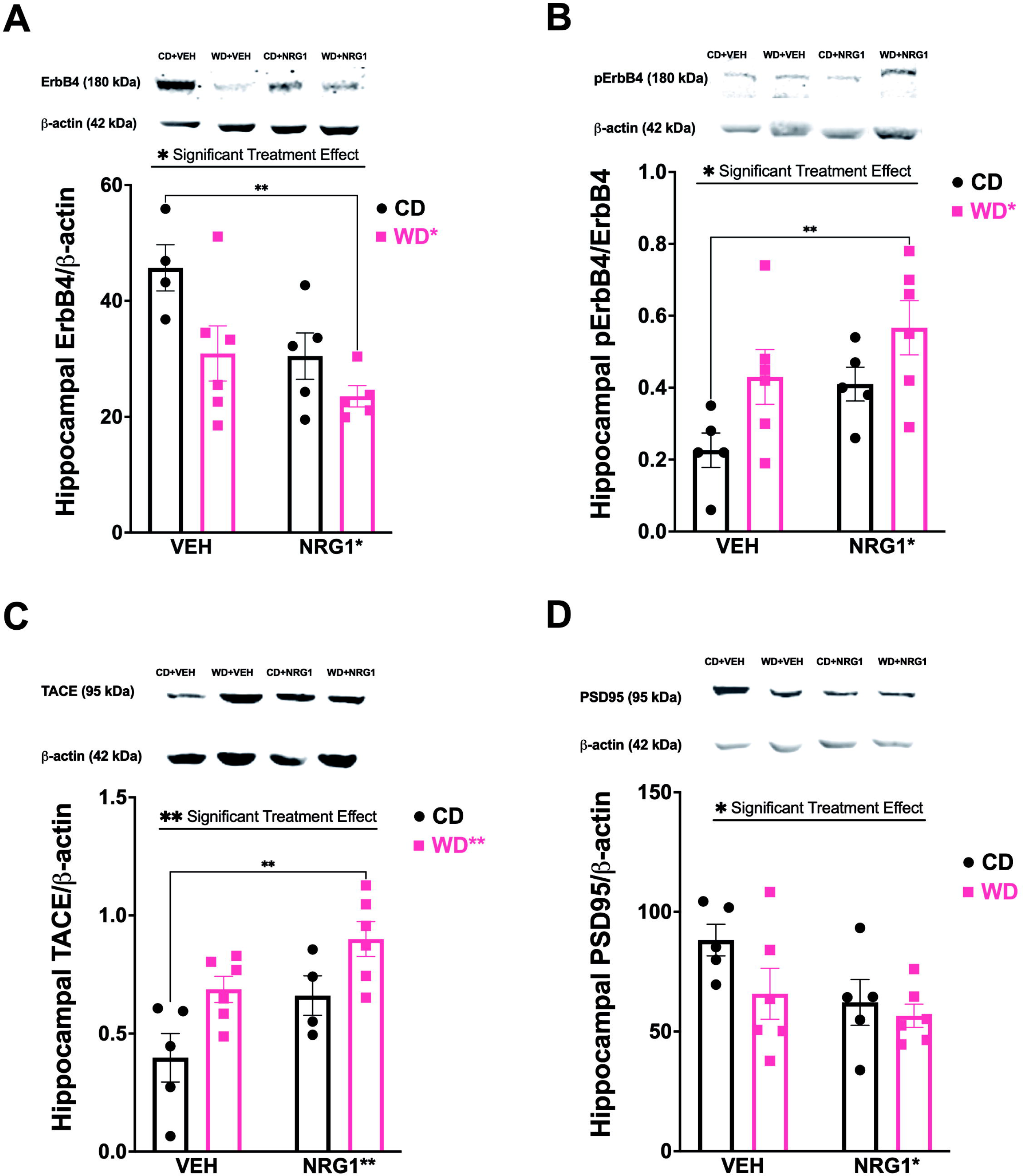
Obesogenic diet intake and subchronic NRG1 administration synergize to promote ErbB4 phosphorylation and TACE/ADAM17 protein levels in the hippocampus. **(A)**Obesogenic WD decreased total ErbB4 protein levels in the hippocampus (diet [*F*_(1, 16)_ = 7.34, *p* = .016]). WDN rats exhibited significantly lower ErbB4 protein levels when compared to CDV rats (*p* = .009). **(B)** Phosphorylation of hippocampal ErbB4 protein levels was affected by the treatment [*F*_(1, 18)_ = 5.82, *p* = .027] and diet [*F*_(1,18)_ = 7.36, *p*= .014]. ErbB4 hyperphosphorylation was particularly evident in WDN rats relative to the CDV group (*p* = .012). **(C)** TACE/ADAM17 protein levels in the hippocampus were affected by the treatment [*F*_(1, 17)_ = 8.93, *p* = .008] and diet [*F*_(1, 17)_ = 11.03, *p* = .004]. WDN rats showed increased TACE/ADAM17 protein levels when compared to CDV rats (*p* = .001). **(D)** NRG1 administration showed a significant decrease for PSD-95 protein levels in the hippocampus (treatment [*F*_(1, 18)_ = 4.47, *p* = .048]). Sample size = 5-6 rats / group.

NRG1 and ErbB4 (trans)activation via proteolytic processing is achieved by the shedding protease TNF-α-Converting Enzyme (*TACE*, also known as *ADAM17*) (Rio et al., 2000; Fleck et al., 2013; Iwakura et al., 2017). We found that the WD increased TACE/ADAM17 protein levels in the hippocampus (**Figure 6C**). We identified a significant treatment [*F*_(1, 17)_ = 8.93, *p* = .008] and diet [*F*_(1, 17)_ = 11.03, *p* = .004] effect, while no significant interaction [*F*_(1, 17)_ = .099, *p* = .76] effects on protein levels. In agreement with the hippocampal pErbB4 protein levels results, we found that WDN rats showed increased TACE/ADAM17 protein levels when compared to CDV rats (*p* = .001).

We measured PSD-95, Akt, and Erk protein levels to investigate the impact of the WD and NRG1 on molecular pathways implicated in ErbB4 signaling. NRG1 administration showed a significant decrease for a postsynaptic regulator of ErbB4 dimerization, PSD-95 (**Figure 6D**). Analyses revealed a significant treatment [*F*_(1, 18)_ = 4.47, *p* = .048], and no significant diet [*F*_(1, 18)_ = 2.83, *p* = .11] or interaction [*F*_(1, 18)_ = 1.02, *p* = .33] effects on PSD-95 protein levels. Analyses for pAkt (normalized to total Akt) showed no significant interaction [*F*_(1, 17)_ = .71, *p* = .41], treatment [*F*_(1, 17)_ = .16, *p*= .69], or diet [*F*_(1, 17)_ = 2.80, *p* = .12] effects on protein levels (**Supplemental Figure 6A**). Similarly, analyses for pErk1/2 (normalized to total Erk) showed no significant interaction [*F*_(1, 18)_ = 1.13, *p* = .30], treatment [*F*_(1, 18)_ = .88, *p* = .36], or diet [*F*_(1, 18)_ = .35, *p* = .56] effects on protein levels (**Supplemental Figure 6B**).

### TACE/ADAM17 protein levels correlate with hippocampal pErbB4

We examined potential mechanisms explaining the reduced ErbB4 expression and increased pErbB4 protein levels in the rats that consumed the obesogenic WD. Twenty-one (21) days of consuming the WD and receiving NRG1 administration was not sufficient to significantly alter NRG1 protein levels in the hippocampus (**Supplemental Figure 7**). Analyses showed no significant interaction [*F*_(1, 18)_ = .14, *p* = .72], treatment [*F*_(1, 18)_ = .031, *p* = .86], or diet [*F*_(1, 18)_ = .38, *p* = .54] effects on hippocampal NRG1 protein levels, as measured by ELISA.

Four (4) structurally and functionally distinct ErbB4 isoforms have been identified. One pair of isoforms differs within their extracellular juxtamembrane domains. These juxtamembrane ErbB4 isoforms are either susceptible (JMa) or resistant (JMb) to proteolytic processing that releases a soluble receptor ectodomain. We used qRT-PCR to examine the effect of the WD and NRG1 on the regulation of hippocampal ErbB4 isoforms. We found lower JMa mRNA levels in the WD rats (relative to CD rats) and higher JMb in the rats that received the exogenous NRG1 (relative to VEH) (**Supplemental Figure 8**). Notably, mixed-effects model analyses revealed a significant interaction between isoform x diet [*F*_(3, 57)_ = 3.13, *p* = .033] and isoform x treatment [*F*_(3, 57)_ = 2.79, *p* = .049].

In summary, our data demonstrate that WD consumption reduces hippocampal ErbB4 protein levels while increasing its phosphorylation. TACE/ADAM17 activities may lead to reduced ErbB4 and also to ErbB4 transactivation (Rio et al., 2000). We performed Pearson’s correlation analyses to test this idea. ADAM17 hippocampal protein levels showed a robust significant correlation with ErbB4 protein levels (*r* = -.59, *p* = .007; **Figure 7A**). ADAM17 protein levels were positively associated with hippocampal pErbB4 protein levels (*r* = .53, *p* = .012; **Figure 7B**). A significant negative relationship was found between ADAM17 protein levels and PSD-95 protein levels in the hippocampus (*r* = -.54, *p* = .012; **Figure 7C**). **Figure 8** describes our working model. We propose that obesogenic diets rich in saturated fatty acids promote TACE/ADAM17-mediated proteolytic cleavage of critical neurodevelopmental and neuroinflammatory processes, and consequently, lead to hippocampal synaptic/structural impairments implicated with anxiety disorders.

**Figure 7.**
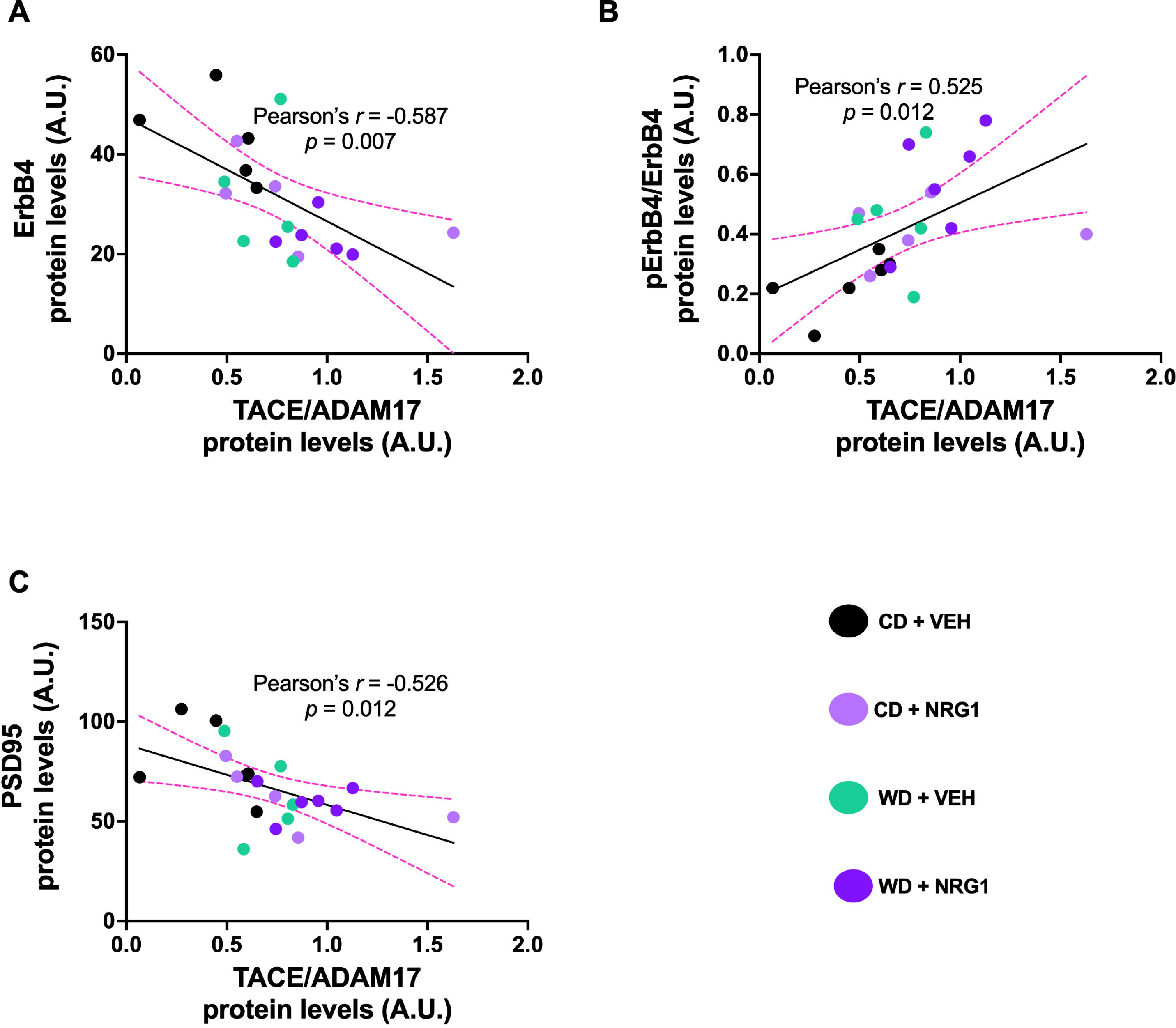
TACE/ADAM17 hippocampal protein levels are associated with ErbB4 activities. **(A)**ADAM17 hippocampal protein levels negatively correlates with ErbB4 protein levels (*r* = -.59, *p* = .007). **(B)** ADAM17 protein levels positively correlates with hippocampal pErbB4 protein levels (*r* = .53, *p* = .012). **(C)** ADAM17 protein levels negatively correlates with PSD-95 protein levels in the hippocampus (*r* = -.54, *p* = .012). Sample size = 5-6 rats / group.

**Figure 8.**
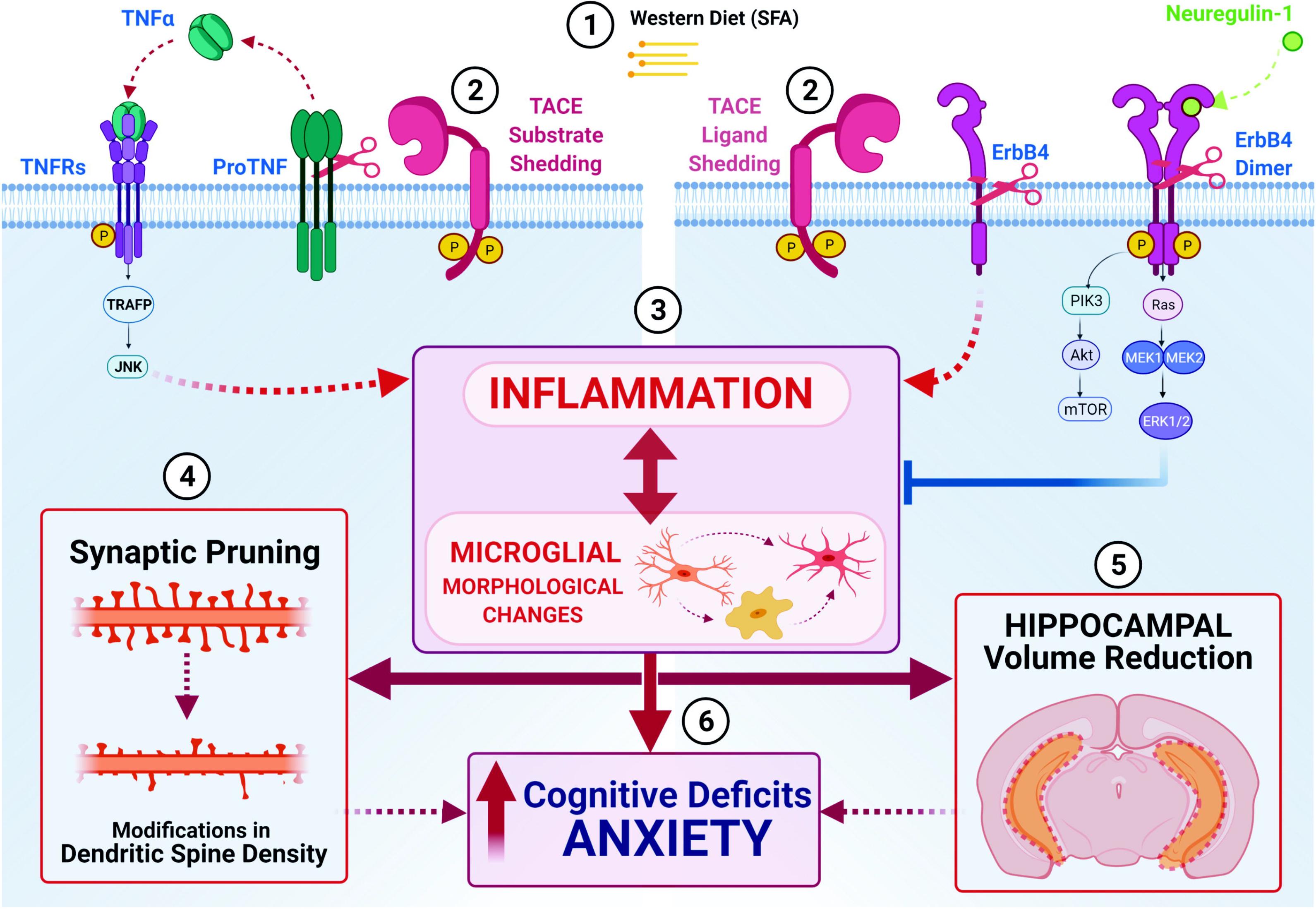
Working model unifying neurodevelopmental and neuroinflammatory hypotheses implicated in obesity-mediated hippocampal dysfunction. (1) It is plausible that exposure to obesogenic diets rich in saturated fatty acids (SFAs) increase TACE/ADAM17 sheddase expression and activities, **(2)** leading to dysregulation of neuroinflammatory (e.g., tumor necrosis factor alfa, TNF-α) and neurodevelopmental (ErbB4) mediators. **(3)** TNF receptor (TNFR) activation and prolonged ErbB4 cleavage tips the inflammatory balance to a pro-inflammatory state associated with microglial activation. **(4)** Neuroinflammation and microglial activation promote synaptic pruning and remodeling. **(5)** Disruption in microglia-synapse interactions contribute to synapse loss and dysfunction, and consequently hippocampal volume atrophy. **(6)** In summary, we postulate that TACE/ADAM17-mediated inflammation, microglial activation, and hippocampal structural impairments contribute to the obesity-induced cognitive and behavioral deficits implicated in anxiety disorders.

## Discussion

This study sought to determine whether altered NRG1-ErbB4 signaling may partly explain hippocampal abnormalities in adolescent rats exposed to an obesogenic Western diet (WD). We demonstrate that both WD consumption and prolonged NRG1 administration during adolescence alter fear and anxiety-like behaviors while reducing hippocampal volume. We also report unique anatomical vulnerabilities to the obesogenic WD, particularly in the CA1 subfield of the hippocampus. These behavioral and anatomical alterations were associated with changes in microglial morphology, unique cytokine profiles, ErbB4 hyperphosphorylation, and increased TACE/ADAM17 protein levels.

### The consumption of an obesogenic diet during early adolescence promotes neuroinflammation and anxiety-like behaviors while increasing hemispheric differences in hippocampal volumes

Obesity and the dietary intake of saturated fats and simple sugars have emerged as risk factors for the development of anxiety and stress-related disorders (Scott et al., 2008; Perkonigg et al., 2009; Johannessen and Berntsen, 2013; Maguen et al., 2013; Mitchell et al., 2013; Kubzansky et al., 2014; Bartoli et al., 2015; Smith et al., 2015). Consistent with studies in humans, we reported that diet-induced obesity (DIO) in rats: **1)** reduces hippocampal volume (Kalyan-Masih et al., 2016), **2)** impairs the maturation of the corticolimbic fear circuits (Vega-Torres et al., 2018), **3)** enhances behavioral vulnerabilities to psychosocial stress (Kalyan-Masih et al., 2016; Vega-Torres et al., 2018, 2019b), **4)** results in profound fear learning and extinction learning deficits (Vega-Torres et al., 2018, 2019b), even in the absence of an obesogenic phenotype (Vega-Torres et al., 2020a), and **5)** leads to alterations in biomarkers implicated in inflammatory responses (Santana et al., 2021).

As valuable as these earlier findings have been in constructing a conceptual framework for the impact of obesogenic environments on hippocampal maturation, our current understanding is primarily based on molecular, cellular, and behavioral analyses of obese versus control animals at the end of dietary manipulations (when rats are considered obese; in our model, approximately at eight weeks after WD exposure). Less is known about the early impact of obesogenic diets on hippocampal structure and function. To address this knowledge gap, we combined volumetric measurements of the hippocampus using magnetic resonance imaging (MRI) with behavioral assays of hippocampus-based tasks after a 21-day exposure to an obesogenic diet during the early adolescence period (from PND 21-41). Using behavioral tasks dependent on optimal hippocampal function, we demonstrated that three weeks on an obesogenic WD increased anxiety-like indices in the elevated plus maze in adolescent rats. The rats exposed to the obesogenic WD also exhibited increased locomotor activity (exploratory behavior) in the open field test. Interestingly, trace fear conditioning, and spatial working memory tasks were not affected in the animals that consumed the WD. MRI-based volumetrics demonstrated that three weeks on the obesogenic diet was sufficient to enhance hippocampal lateralization effects. These behavioral and anatomical alterations were associated with increased pro-inflammatory cytokines and changes in microglial morphology related to phagocytic phenotypes. Together with our previous studies, the findings reported here indicate that exposure to an obesogenic diet during adolescence selectively affects hippocampal structure and specific cognitive domains. Our investigations demonstrate that early adolescence is a critical window during which obesogenic diets exert anxiogenic effects.

### Hyper-NRG1 conditions attenuate trace fear conditioning and reduce hippocampal volume while influencing microglia shape and inflammatory mediators

Neuregulin-1 (NRG1) and its ErbB receptor tyrosine kinases are expressed not only in the developing nervous system but also in the adult brain. ErbB4 is likely to be the primary mediator of NRG1 functions in the brain, and it is mainly expressed in hippocampal GABAergic interneurons, where it is enriched at postsynaptic terminals (Vullhorst et al., 2009). NRG1 signaling participates in several critical neurodevelopmental processes and is implicated in nerve cell differentiation and synapse formation, radial glia formation and neuronal migration, axon navigation, and neurite outgrowth (Mei and Nave, 2014). Perturbations in NRG1/ErbB4 function have been associated with various neuropsychiatric disorders (Mei and Nave, 2014), resilience to stress (Clarke et al., 2018), and obesity/metabolic syndrome (Ennequin et al., 2015a; Salinas et al., 2016; Zeng et al., 2018; Zhang et al., 2018). Emerging evidence suggests that ErbB4 may also be important in the pathogenesis of obesity in humans (Locke et al., 2015; Salinas et al., 2016).

A key finding of this study is that prolonged NRG1 administration during early adolescence resulted in reduced hippocampal volume and fear conditioning deficits. Evidence suggests that a gain of function of NRG1 may contribute to the pathophysiology of schizophrenia (Hashimoto et al., 2004; Chong et al., 2008; Paterson et al., 2014). In agreement, a mouse model overexpressing NRG1 exhibited marked alterations in hippocampal-dependent function and memory while impacting the expression of several genes implicated in inflammation (Deakin et al., 2012). Interestingly, this study reported increased hippocampal volume in the NRG1 transgenic mice (Deakin et al., 2012). The conflicting result on hippocampal volume may be related to the magnitude in NRG1 levels, the developmental stage of the animals, animal species, or NRG1 peripheral effects (e.g., robust leptin and FGF21 secretagogue). Together, it is clear that NRG1 is a potent modulator of hippocampal volume and cognition.

Microglia cells, the resident immune cells in the brain, are responsible for maintaining a dynamic balance between anti-inflammatory and pro-inflammatory mediators. Microglia cells play a significant role in the behavioral outcomes associated with obesity (Sobesky et al., 2014; Guillemot-Legris and Muccioli, 2017; Cope et al., 2018). However, the underlying molecular mechanisms regulating the microglia-dependent inflammatory balance remain poorly understood. Microglia express ErbB2, 3, and 4 receptors (Gerecke et al., 2001; Dimayuga et al., 2003; Calvo et al., 2010) and are highly responsive to NRG1 (Simmons et al., 2016; Alizadeh et al., 2017). Yet, the function of this signaling pathway within microglia is poorly understood. While it is very likely that the observed NRG1 effects are related to changes in local microcircuits (GABAergic; parvalbumin, PV+ interneurons exhibit high ErbB4 expression; (Neddens and Buonanno, 2009)), this study provides additional support to the notion that NRG1-ErbB4 signaling regulates microglial morphology and inflammatory mediators (Calvo et al., 2010; Liew et al., 2016; Simmons et al., 2016; Chen et al., 2019; Shahriary et al., 2019).

### An obesogenic diet influences behavioral, anatomical, and molecular responses to exogenous NRG1 administration

Consistent with studies demonstrating the potential involvement of NRG-ErbB signaling in obesity (Ennequin et al., 2015a; Locke et al., 2015; Cai et al., 2016; Salinas et al., 2016; Zeng et al., 2018; Zhang et al., 2018), we found several significant interactions between the obesogenic diet and exogenous NRG1. We present new evidence that an obesogenic diet influences the effect of prolonged NRG1 administration on the fear-potentiated startle, total hippocampal and CA1 subfield volume, and hippocampal cytokine levels (TNF-α, IL-10, IL-6). Interestingly, exogenous NRG1 synergized with the obesogenic diet to enhance ErbB4 activation and its proteolytic processing (Rio et al., 2000). ErbB4 proteolytic cleavage is mediated by the tumor necrosis factor-α-converting enzyme (TACE) / a Disintegrin And Metalloproteinase 17 (ADAM17)-dependent ectodomain shedding in a protein kinase C (PKC)-dependent manner. Here, we present data suggesting a role for TACE/ADAM17 on ErbB4 activities and synaptic integrity. Future studies will identify the consequences of dysregulated NRG1/ErbB4 ectodomain shedding on hippocampal maturation and cognitive function.

### Limitations

Our results demonstrate significant alterations in neurotrophic and neuroinflammatory signals in adolescent rats that consume an obesogenic diet enriched in saturated fats and simple sugars. However, the temporal window in which these effects take place was not established. To better clarify a mechanism, the spatiotemporal expression of hippocampal ErbB4 and associated cytokine profiles must be determined. Interpretation of the study may be confounded by the effect of daily i.p. injections. While this represents one of the best approaches to achieve therapeutic NRG1 levels in circulation (NRG1 has a short half-life), it is possible that the subchronic daily injection regime affected some endpoint measures. Furthermore, it is not clear from our results whether the NRG1 effects on hippocampal structure and behaviors are direct (and whether ErbB4 is responsible for the alterations). The use of inhibitors (e.g., AG1478) or fusion proteins (e.g., ecto-ErbB4) needs to be considered to dissect mechanisms and determine causality. Future experiments are required to dissociate the chronic vs. acute response to the NRG1 injection on molecular targets. Additionally, the study should be replicated in female rats and other rodent strains.

## Conclusion

Current pathophysiological models explaining how early exposure to obesogenic diets alters brain and behavior have focused on neurodevelopmental or immunological mechanisms. We report a novel interaction between these factors in the maturing hippocampus. Our results indicate that a short-term (21 days) dietary challenge with an obesogenic WD during the critical neurodevelopmental stage of adolescence disrupts ErbB4 activities in the hippocampus. Most importantly, we continue to provide evidence that WD consumption during adolescence leads to neuroanatomical, inflammatory, and molecular alterations related to depression, anxiety disorders, and cognitive impairments in humans. While extrapolating our data must be made carefully, our results indicate that TACE/ADAM17-ErbB4 hyperfunction may contribute to abnormal hippocampal structural and cognitive vulnerabilities in individuals who consume obesogenic diets.

## Acknowledgments

This study was partly supported by the NIH (DK124727, GM060507, MD006988) and the Loma Linda University School of Medicine GRASP Seed Funds to JDF and IDP.

---

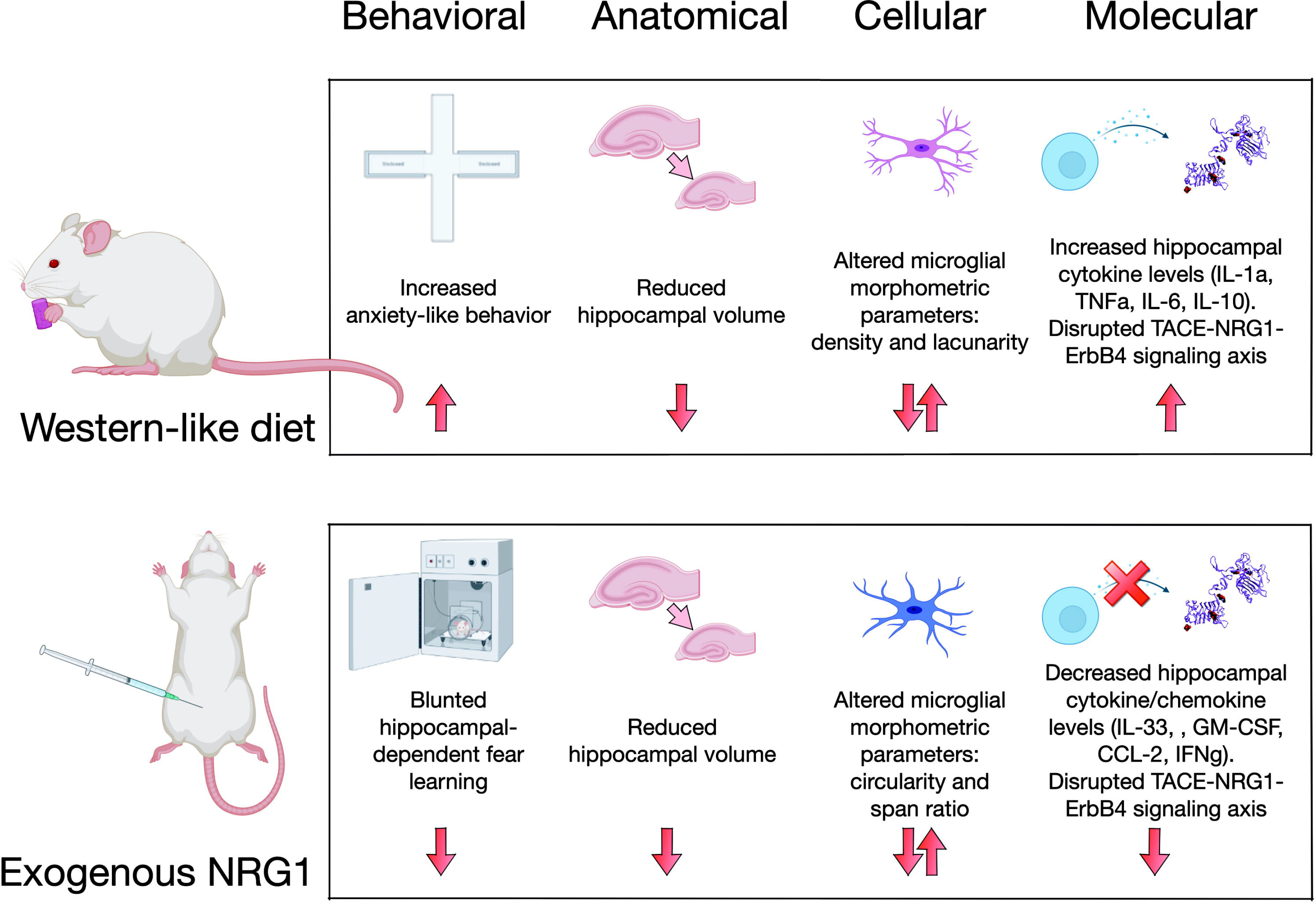

## References

1. Alizadeh A, Dyck SM, Kataria H, Shahriary GM, Nguyen DH, Santhosh KT, Karimi-Abdolrezaee S. 2017. Neuregulin-1 positively modulates glial response and improves neurological recovery following traumatic spinal cord injury. Glia 65:1152–1175.

2. Bartoli F, Crocamo C, Alamia A, Amidani F, Paggi E, Pini E, Clerici M, Carrà G. 2015. Posttraumatic Stress Disorder and Risk of Obesity. J Clin Psychiatry 76:e1253–61.

3. Bauer CCC, Moreno B, González-Santos L, Concha L, Barquera S, Barrios FA. 2015. Child overweight and obesity are associated with reduced executive cognitive performance and brain alterations: a magnetic resonance imaging study in Mexican children. Pediatr Obes 10:196–204.

4. Cai C, Lin M, Xu Y, Li X, Yang S, Zhang H. 2016. Association of circulating neuregulin 4 with metabolic syndrome in obese adults: a cross-sectional study. Bmc Med 14:165.

5. Caillaud K, Boisseau N, Ennequin G, Chavanelle V, Etienne M, Li X, Denis P, Dardevet D, Lacampagne A, Sirvent P. 2016. Neuregulin 1 improves glucose tolerance in adult and old rats. Diabetes Metab 42:96–104.

6. Calvo M, Zhu N, Tsantoulas C, Ma Z, Grist J, Loeb JA, Bennett DLH. 2010. Neuregulin-ErbB Signaling Promotes Microglial Proliferation and Chemotaxis Contributing to Microgliosis and Pain after Peripheral Nerve Injury. J Neurosci 30:5437–5450.

7. Chen J, Zhou X, Ding H, Zhan H, Yang F, Li W, Xie J, Liu X, Xu Y, Su M, Liu B, Zhou X. 2019. Neuregulin-1-ErbB signaling promotes microglia activation contributing to mechanical allodynia of cyclophosphamide-induced cystitis. Neurourol Urodynam 38:1250–1260.

8. Cherbuin N, Sargent-Cox K, Fraser M, Sachdev P, Anstey KJ. 2015. Being overweight is associated with hippocampal atrophy: the PATH Through Life Study. Int J Obesity 39:1509–1514.

9. Chong VZ, Thompson M, Beltaifa S, Webster MJ, Law AJ, Weickert CS. 2008. Elevated neuregulin-1 and ErbB4 protein in the prefrontal cortex of schizophrenic patients. Schizophr Res 100:270–280.

10. Clarke DJ, Chohan TW, Kassem MS, Smith KL, Chesworth R, Karl T, Kuligowski MP, Fok SY, Bennett MR, Arnold JC. 2018. Neuregulin 1 Deficiency Modulates Adolescent Stress-Induced Dendritic Spine Loss in a Brain Region-Specific Manner and Increases Complement 4 Expression in the Hippocampus. Schizophrenia Bull 43:486.

11. Cohen H, Kozlovsky N, Alona C, Matar MA, Joseph Z. 2012. Animal model for PTSD: from clinical concept to translational research. Neuropharmacology 62:715–724.

12. Contreras CM, Rodríguez-Landa JF, García-Ríos RI, Cueto-Escobedo J, Guillen-Ruiz G, Bernal-Morales B. 2014. Myristic Acid Produces Anxiolytic-Like Effects in Wistar Rats in the Elevated Plus Maze. Biomed Res Int 2014:1–8.

13. Cope EC, LaMarca EA, Monari PK, Olson LB, Martinez S, Zych AD, Katchur NJ, Gould E. 2018. Microglia Play an Active Role in Obesity-Associated Cognitive Decline. J Neurosci 38:8889–8904.

14. Dang R, Cai H, Zhang L, Liang D, Lv C, Guo Y, Yang R, Zhu Y, Jiang P. 2016. Dysregulation of Neuregulin-1/ErbB signaling in the prefrontal cortex and hippocampus of rats exposed to chronic unpredictable mild stress. Physiol Behav 154:145–150.

15. Deakin IH, Nissen W, Law AJ, Lane T, Kanso R, Schwab MH, Nave K-A, Lamsa KP, Paulsen O, Bannerman DM, Harrison PJ. 2012. Transgenic Overexpression of the Type I Isoform of Neuregulin 1 Affects Working Memory and Hippocampal Oscillations but not Long-term Potentiation. Cereb Cortex 22:1520–1529.

16. Depboylu C, Rösler TW, Andrade A de, Oertel WH, Höglinger GU. 2015. Systemically administered neuregulin-1β1 rescues nigral dopaminergic neurons via the ErbB4 receptor tyrosine kinase in MPTP mouse models of Parkinson’s disease. J Neurochem 133:590–597.

17. Dimayuga FO, Ding Q, Keller JN, Marchionni MA, Seroogy KB, Bruce-Keller AJ. 2003. The neuregulin GGF2 attenuates free radical release from activated microglial cells. J Neuroimmunol 136:67–74.

18. Ennequin G, Boisseau N, Caillaud K, Chavanelle V, Etienne M, Li X, Montaurier C, Sirvent P. 2015a. Neuregulin 1 affects leptin levels, food intake and weight gain in normal-weight, but not obese, db/db mice. Diabetes Metab 41:168–172.

19. Ennequin G, Boisseau N, Caillaud K, Chavanelle V, Etienne M, Li X, Sirvent P. 2015b. Neuregulin 1 Improves Glucose Tolerance in db/db Mice. Plos One 10:e0130568.

20. Fernández-Arjona M del M, Grondona JM, Granados-Durán P, Fernández-Llebrez P, López-Ávalos MD. 2017. Microglia Morphological Categorization in a Rat Model of Neuroinflammation by Hierarchical Cluster and Principal Components Analysis. Front Cell Neurosci 11:121–22.

21. Fisahn A, Neddens J, Yan L, Buonanno A. 2009. Neuregulin-1 Modulates Hippocampal Gamma Oscillations: Implications for Schizophrenia. Cereb Cortex 19:612–618.

22. Fleck D, Bebber F van, Colombo A, Galante C, Schwenk BM, Rabe L, Hampel H, Novak B, Kremmer E, Tahirovic S, Edbauer D, Lichtenthaler SF, Schmid B, Willem M, Haass C. 2013. Dual Cleavage of Neuregulin 1 Type III by BACE1 and ADAM17 Liberates Its EGF-Like Domain and Allows Paracrine Signaling. J Neurosci 33:7856– 7869.

23. Gao R, Ji M-H, Gao D-P, Yang R-H, Zhang S-G, Yang J-J, Shen J-C. 2016. Neuroinflammation-Induced Downregulation of Hippocampacal Neuregulin 1-ErbB4 Signaling in the Parvalbumin Interneurons Might Contribute to Cognitive Impairment in a Mouse Model of Sepsis-Associated Encephalopathy. Inflammation 40:1–14.

24. Gerecke KM, Wyss JM, Carroll SL. 2004. Neuregulin-1beta induces neurite extension and arborization in cultured hippocampal neurons. Mol Cell Neurosci 27:379–393.

25. Gerecke KM, Wyss JM, Karavanova I, Buonanno A, Carroll SL. 2001. ErbB transmembrane tyrosine kinase receptors are differentially expressed throughout the adult rat central nervous system. The Journal of Comparative Neurology 433:86–100.

26. Guan Y-F, Wu C-Y, Fang Y-Y, Zeng Y-N, Luo Z-Y, Li S-J, Li X-W, Zhu X-H, Mei L, Gao T-M. 2015. Neuregulin 1 protects against ischemic brain injury via ErbB4 receptors by increasing GABAergic transmission. Neuroscience 307:151–159.

27. Guillemot-Legris O, Muccioli GG. 2017. Obesity-Induced Neuroinflammation: Beyond the Hypothalamus. Trends Neurosci 40:237–253.

28. Hao S, Dey A, Yu X, Stranahan AM. 2016. Dietary obesity reversibly induces synaptic stripping by microglia and impairs hippocampal plasticity. Brain Behav Immun 51:230-239.

29. Hashimoto R, Straub RE, Weickert CS, Hyde TM, Kleinman JE, Weinberger DR. 2004. Expression analysis of neuregulin-1 in the dorsolateral prefrontal cortex in schizophrenia. Mol Psychiatr 9:299–307.

30. Holm-Hansen S, Low JK, Zieba J, Gjedde A, Bergersen LH, Karl T. 2016. Behavioural effects of high fat diet in a mutant mouse model for the schizophrenia risk gene neuregulin 1. Genes Brain Behav 15:295–304.

31. Hruby A, Lieberman HR, Smith TJ. 2021. Symptoms of depression, anxiety, and post-traumatic stress disorder and their relationship to health-related behaviors in over 12,000 US military personnel: Bi-directional associations. J Affect Disorders 283:84– 93.

32. Iwakura Y, Wang R, Inamura N, Araki K, Higashiyama S, Takei N, Nawa H. 2017. Glutamate-dependent ectodomain shedding of neuregulin-1 type II precursors in rat forebrain neurons. Plos One 12:e0174780.

33. Jacka FN, Cherbuin N, Anstey KJ, Sachdev P, Butterworth P. 2015. Western diet is associated with a smaller hippocampus: a longitudinal investigation. Bmc Med 13:1–9.

34. Jacobson L, Sapolsky R. 1991. The role of the hippocampus in feedback regulation of the hypothalamic-pituitary-adrenocortical axis. Endocr Rev 12:118–134.

35. Jalilzad M, Jafari A, Babaei P. 2019. Neuregulin1β improves both spatial and associative learning and memory in Alzheimer model of rats possibly through signaling pathways other than Erk1/2. Neuropeptides 78:101963.

36. Jimenez JC, Su K, Goldberg AR, Luna VM, Biane JS, Ordek G, Zhou P, Ong SK, Wright MA, Zweifel L, Paninski L, Hen R, Kheirbek MA. 2018. Anxiety Cells in a Hippocampal-Hypothalamic Circuit. Neuron 97:670–683.e6.

37. Johannessen KB, Berntsen D. 2013. Losing the symptoms: weight loss and decrease in posttraumatic stress disorder symptoms. J Clin Psychol 69:655–660.

38. Kalyan-Masih P, Vega-Torres JD, Miles C, Haddad E, Rainsbury S, Baghchechi M, Obenaus A, Figueroa JD. 2016. Western High-fat Diet Consumption During Adolescence Increases Susceptibility to Traumatic Stress while Selectively Disrupting Hippocampal and Ventricular Volumes. Eneuro 3:ENEURO.0125-16.2016.

39. Kanoski SE, Hayes MR, Greenwald HS, Fortin SM, Gianessi CA, Gilbert JR, Grill HJ. 2011. Hippocampal leptin signaling reduces food intake and modulates food-related memory processing. Neuropsychopharmacol 36:1859–1870.

40. Karperien A, Ahammer H, Jelinek HF. 2013. Quantitating the subtleties of microglial morphology with fractal analysis. Front Cell Neurosci 7:3.

41. Karperien AL, Jelinek HF. 2015. Fractal, Multifractal, and Lacunarity Analysis of Microglia in Tissue Engineering. Frontiers Bioeng Biotechnology 3:51.

42. Kubzansky LD, Bordelois P, Jun HJ, Roberts AL, Cerda M, Bluestone N, Koenen KC. 2014. The Weight of Traumatic Stress: A Prospective Study of Posttraumatic Stress Disorder Symptoms and Weight Status in Women. Jama Psychiat 71:44–51.

43. Kwon O-B, Paredes D, Gonzalez CM, Neddens J, Hernandez L, Vullhorst D, Buonanno A. 2008. Neuregulin-1 regulates LTP at CA1 hippocampal synapses through activation of dopamine D4 receptors. Proc National Acad Sci 105:15587–15592.

44. Li Y, Lein PJ, Ford GD, Liu C, Stovall KC, White TE, Bruun DA, Tewolde T, Gates AS, Distel TJ, Surles-Zeigler MC, Ford BD. 2015. Neuregulin-1 inhibits neuroinflammatory responses in a rat model of organophosphate-nerve agent-induced delayed neuronal injury. J Neuroinflamm 12:64.

45. Li Y, Lein PJ, Liu C, Bruun DA, Giulivi C, Ford GD, Tewolde T, Ross-Inta C, Ford BD. 2012. Neuregulin-1 is neuroprotective in a rat model of organophosphate-induced delayed neuronal injury. Toxicol Appl Pharm 262:194–204.

46. Liew H, Kim Y-M, Choi HS, Jang AR, Churchill D, Lee SH, Suh Y-H. 2016. Soluble Neuregulin-1 from Microglia Enhances Amyloid Beta-induced Neuronal Death. Cns Neurological Disord - Drug Targets 15:918–926.

47. Liu CM, Kanoski SE. 2018. Homeostatic and non-homeostatic controls of feeding behavior: Distinct vs. common neural systems. Physiol Behav 193:223–231.

48. Liu M, Solomon W, Cespedes JC, Wilson NO, Ford B, Stiles JK. 2018. Neuregulin-1 attenuates experimental cerebral malaria (ECM) pathogenesis by regulating ErbB4/AKT/STAT3 signaling. J Neuroinflamm 15:104.

49. Locke AE, Kahali B, Berndt SI, Justice AE, Pers TH, Day FR, Powell C, Vedantam S, Buchkovich ML, Yang J, Croteau-Chonka DC, Esko T, Fall T, Ferreira T, Gustafsson S, Kutalik Z, Luan J, Mägi R, Randall JC, Winkler TW, Wood AR, Workalemahu T, Faul JD, Smith JA, Zhao JH, Zhao W, Chen J, Fehrmann R, Hedman ÅK, Karjalainen J, Schmidt EM, Absher D, Amin N, Anderson D, Beekman M, Bolton JL, Bragg-Gresham JL, Buyske S, Demirkan A, Deng G, Ehret GB, Feenstra B, Feitosa MF, Fischer K, Goel A, Gong J, Jackson AU, Kanoni S, Kleber ME, Kristiansson K, Lim U, Lotay V, Mangino M, Leach IM, Medina-Gomez C, Medland SE, Nalls MA, Palmer CD, Pasko D, Pechlivanis S, Peters MJ, Prokopenko I, Shungin D, Stančáková A, Strawbridge RJ, Sung YJ, Tanaka T, Teumer A, Trompet S, Laan SW van der, Setten J van, Vliet-Ostaptchouk JVV, Wang Z, Yengo L, Zhang W, Isaacs A, Albrecht E, Ärnlöv J, Arscott GM, Attwood AP, Bandinelli S, Barrett A, Bas IN, Bellis C, Bennett AJ, Berne C, Blagieva R, Blüher M, Böhringer S, Bonnycastle LL, Böttcher Y, Boyd HA, Bruinenberg M, Caspersen IH, Chen Y-DI, Clarke R, Daw EW, Craen AJMD, et al. 2015. Genetic studies of body mass index yield new insights for obesity biology. Nature 518:197–206.

50. López-Soldado I, Niisuke K, Veiga C, Adrover A, Manzano A, Martínez-Redondo V, Camps M, Bartrons R, Zorzano A, Gumà A. 2016. Neuregulin improves response to glucose tolerance test in control and diabetic rats. Am J Physiol-endoc M 310:E440–51.

51. Lopresti AL, Drummond PD. 2013. Obesity and psychiatric disorders: commonalities in dysregulated biological pathways and their implications for treatment. Prog Neuro-psychopharmacology Biological Psychiatry 45:92–99.

52. Maguen S, Madden E, Cohen B, Bertenthal D, Neylan T, Talbot L, Grunfeld C, Seal K. 2013. The relationship between body mass index and mental health among Iraq and Afghanistan veterans. J Gen Intern Med 28 Suppl 2:S563–70.

53. Mahar I, Labonte B, Yogendran S, Isingrini E, Perret L, Davoli MA, Rachalski A, Giros B, Turecki G, Mechawar N. 2017. Disrupted hippocampal neuregulin-1/ErbB3 signaling and dentate gyrus granule cell alterations in suicide. Transl Psychiat 7:e1161.

54. Mahar I, MacIsaac A, Kim JJ, Qiang C, Davoli MA, Turecki G, Mechawar N. 2016. Effects of neuregulin-1 administration on neurogenesis in the adult mouse hippocampus, and characterization of immature neurons along the septotemporal axis. Sci Rep-uk 6:30467.

55. Mahar I, Tan S, Davoli MA, Dominguez-Lopez S, Qiang C, Rachalski A, Turecki G, Mechawar N. 2011. Subchronic peripheral neuregulin-1 increases ventral hippocampal neurogenesis and induces antidepressant-like effects. Plos One 6:e26610.

56. McGlinchey-Berroth R, Carrillo MC, Gabrieli JDE, Brawn CM, Disterhoft JF. 1997. Impaired Trace Eyeblink Conditioning in Bilateral, Medial-Temporal Lobe Amnesia. Behav Neurosci 111:873–882.

57. Mei L, Nave K-A. 2014. Neuregulin-ERBB Signaling in the Nervous System and Neuropsychiatric Diseases. Neuron 83:27–49.

58. Mei L, Xiong W-C. 2008. Neuregulin 1 in neural development, synaptic plasticity and schizophrenia. Nat Rev Neurosci 9:437–452.

59. Mencel M, Nash M, Jacobson C. 2013. Neuregulin Upregulates Microglial α7 Nicotinic Acetylcholine Receptor Expression in Immortalized Cell Lines: Implications for Regulating Neuroinflammation. Plos One 8:e70338.

60. Mestre ZL, Bischoff-Grethe A, Eichen DM, Wierenga CE, Strong D, Boutelle KN. 2017. Hippocampal atrophy and altered brain responses to pleasant tastes among obese compared with healthy weight children. Int J Obesity 41:1496–1502.

61. Michopoulos V, Vester A, Neigh G. 2016. Posttraumatic stress disorder: A metabolic disorder in disguise? Exp Neurol 284:220–229.

62. Mitchell KS, Aiello AE, Galea S, Uddin M, Wildman D, Koenen KC. 2013. PTSD and obesity in the Detroit neighborhood health study. Gen Hosp Psychiat 35:671–673.

63. Moreno LA, Rodriguez G, Fleta J, Bueno-Lozano M, Lazaro A, Bueno G. 2010. Trends of dietary habits in adolescents. Crit Rev Food Sci 50:106–112.

64. Neddens J, Buonanno A. 2009. Selective populations of hippocampal interneurons express ErbB4 and their number and distribution is altered in ErbB4 knockout mice. Hippocampus 99:NA NA.

65. Nelson LH, Peketi P, Lenz KM. 2021. Microglia Regulate Cell Genesis in a Sex-dependent Manner in the Neonatal Hippocampus. Neuroscience 453:237–255.

66. Paterson C, Wang Y, Kleinman JE, Law AJ. 2014. Effects of Schizophrenia Risk Variation in the NRG1 Gene on NRG1-IV Splicing During Fetal and Early Postnatal Human Neocortical Development. Am J Psychiat 171:979–989.

67. Perkonigg A, Owashi T, Stein MB, Kirschbaum C, Wittchen H-U. 2009. Posttraumatic stress disorder and obesity: evidence for a risk association. Am J Prev Med 36:1–8.

68. Raji CA, Ho AJ, Parikshak NN, Becker JT, Lopez OL, Kuller LH, Hua X, Leow AD, Toga AW, Thompson PM. 2010. Brain structure and obesity. Hum Brain Mapp 31:353– 364.

69. Rio C, Buxbaum JD, Peschon JJ, Corfas G. 2000. Tumor Necrosis Factor-α-converting Enzyme Is Required for Cleavage of erbB4/HER4*. J Biol Chem 275:10379–10387.

70. Rösler TW, Depboylu C, Arias-Carrión O, Wozny W, Carlsson T, Höllerhage M, Oertel WH, Schrattenholz A, Höglinger GU. 2011. Biodistribution and brain permeability of the extracellular domain of neuregulin-1-β1. Neuropharmacology 61:1413–1418.

71. Ryu J, Hong B-H, Kim Y-J, Yang E-J, Choi M, Kim H, Ahn S, Baik T-K, Woo R-S, Kim H-S. 2016. Neuregulin-1 attenuates cognitive function impairments in a transgenic mouse model of Alzheimer’s disease. Cell Death Dis 7:e2117.

72. Salari-Moghaddam A, Keshteli AH, Afshar H, Esmaillzadeh A, Adibi P. 2018. Association between dietary inflammatory index and psychological profile in adults. Clin Nutr.

73. Salinas YD, Wang L, DeWan AT. 2016. Multiethnic genome-wide association study identifies ethnic-specific associations with body mass index in Hispanics and African Americans. Bmc Genet 17:78.

74. Santana JMS, Vega-Torres JD, Ontiveros-Angel P, Lee JB, Torres YA, Gonzalez AYC, Boria EA, Ortiz DZ, Carmona CA, Figueroa JD. 2021. Oxidative stress and neuroinflammation in a rat model of co-morbid obesity and psychogenic stress. Behav Brain Res 400:112995.

75. Schafer DP, Stevens B. 2013. Phagocytic glial cells: sculpting synaptic circuits in the developing nervous system. Curr Opin Neurobiol 23:1034–1040.

76. Scott KM, McGee MA, Wells JE, Browne MAO. 2008. Obesity and mental disorders in the adult general population. J Psychosom Res 64:97–105.

77. Shahriary GM, Kataria H, Karimi-Abdolrezaee S. 2019. Neuregulin-1 Fosters Supportive Interactions between Microglia and Neural Stem/Progenitor Cells. Stem Cells Int 2019:1–20.

78. Simmons LJ, Surles-Zeigler MC, Li Y, Ford GD, Newman GD, Ford BD. 2016. Regulation of inflammatory responses by neuregulin-1 in brain ischemia and microglial cells in vitro involves the NF-kappa B pathway. J Neuroinflamm 13:237.

79. Smith BN, Tyzik AL, Neylan TC, Cohen BE. 2015. PTSD and obesity in younger and older veterans: Results from the mind your heart study. Psychiat Res 229:895–900.

80. Sobesky JL, Barrientos RM, May HSD, Thompson BM, Weber MD, Watkins LR, Maier SF. 2014. High-fat diet consumption disrupts memory and primes elevations in hippocampal IL-1β, an effect that can be prevented with dietary reversal or IL-1 receptor antagonism. Brain Behav Immun 42:22–32.

81. Stevenson RJ, Francis HM. 2017. The Hippocampus and the Regulation of Human Food Intake. Psychol Bull 143:1011–1032.

82. Tan G-H, Liu Y-Y, Hu X-L, Yin D-M, Mei L, Xiong Z-Q. 2012. Neuregulin 1 represses limbic epileptogenesis through ErbB4 in parvalbumin-expressing interneurons. Nat Neurosci 15:258–266.

83. Trivedi MA, Coover GD. 2006. Neurotoxic lesions of the dorsal and ventral hippocampus impair acquisition and expression of trace-conditioned fear-potentiated startle in rats. Behav Brain Res 168:289–298.

84. Vega-Torres JD, Azadian M, Rios-Orsini R, Reyes-Rivera AL, Ontiveros-Angel P, Figueroa JD. 2020a. Early maturational emergence of adult-like emotional reactivity and anxiety after brief exposure to an obesogenic diet. Biorxiv:2020.03.03.975789.

85. Vega-Torres JD, Azadian M, Rios-Orsini RA, Reyes-Rivera AL, Ontiveros-Angel P, Figueroa JD. 2020b. Adolescent Vulnerability to Heightened Emotional Reactivity and Anxiety After Brief Exposure to an Obesogenic Diet. Front Neurosci-switz 14:562.

86. Vega-Torres JD, Haddad E, Lee JB, Kalyan-Masih P, George WIM, Pérez LL, Vázquez DMP, Torres YA, Santana JMS, Obenaus A, Figueroa JD. 2018. Exposure to an obesogenic diet during adolescence leads to abnormal maturation of neural and behavioral substrates underpinning fear and anxiety. Brain Behav Immun.

87. Vega-Torres JD, Kalyan-Masih P, Argueta DA, DiPatrizio NV, Figueroa JD. 2019a. Endocrine, metabolic, and endocannabinoid correlates of obesity in rats exhibiting high anxiety-related behaviors. Matters Sel.

88. Vega-Torres JD, Reyes-Rivera AL, Figueroa JD. 2019b. Developmental regulation of fear memories by an obesogenic high-saturated fat/high-sugar diet. Biorxiv:748079.

89. Vullhorst D, Neddens J, Karavanova I, Tricoire L, Petralia RS, McBain CJ, Buonanno A. 2009. Selective expression of ErbB4 in interneurons, but not pyramidal cells, of the rodent hippocampus. J Neurosci 29:12255–12264.

90. Zeng F, Wang Y, Kloepfer LA, Wang S, Harris RC. 2018. ErbB4 deletion predisposes to development of metabolic syndrome in mice. Am J Physiol-endoc M 96:1025.

91. Zhang H, Zhang L, Zhou D, He X, Wang D, Pan H, Zhang X, Mei Y, Qian Q, Zheng T, Jones FE, Sun B. 2017. Ablating ErbB4 in PV neurons attenuates synaptic and cognitive deficits in an animal model of Alzheimer’s disease. Neurobiol Dis 106:171–180.

92. Zhang P, Kuang H, He Y, Idiga SO, Li S, Chen Z, Yang Z, Cai X, Zhang K, Potthoff MJ, Xu Y, Lin JD. 2018. NRG1-Fc improves metabolic health via dual hepatic and central action. Jci Insight 3:85.

93. Zieba J, Morris MJ, Karl T. 2019a. Behavioural effects of high fat diet exposure starting in late adolescence in neuregulin 1 transmembrane domain mutant mice. Behav Brain Res 373:112074.

94. Zieba J, Morris MJ, Weickert CS, Karl T. 2019b. Behavioural effects of high fat diet in adult Nrg1 type III transgenic mice. Behav Brain Res 377:112217.

