## Supplemental Methods, Tables, and Figures for "Early neuroadaptations to an obesogenic diet identify the schizophrenia-related ErbB4 receptor in obesity-induced hippocampal abnormalities"

**Supplemental Materials**

**Supplemental Table 1. Macronutrient composition of the custom purified diets.** Detailed composition of the matched low-fat purified control diet (CD, 5-gm% fat, product *#F7463*) and Western-like high-saturated fat diet (WD, 20-gm% fat, product *#F7462*).

**Supplemental Table 2. List of antibodies.**

**Supplemental Table 3. Microglial descriptors and definitions.**

**Supplemental Figure 1. Obesogenic diet and prolonged NRG1 administration attenuate startle reactivity in a tone-dependent manner.** Each tone intensity from the trace conditioning protocol was analyzed (tone alone vs. light + tone; 90 dB [*F*_(1, 40)_ = 64.82, *p* < .0001], 95 dB [*F*_(1, 44)_ = 27.71, *p* < .0001], and 105 dB [*F*_(1, 43)_ = 37.61, *p* < .0001]) and only CDV rats exhibited significant differences between the stimulus type in all three tone intensities. Sample size = 12 rats / group.

**Supplemental Figure 2. Obesogenic diet reduces right hippocampal volume. (A)** Total brain volume was not affect by treatment [*F*_(1, 19)_ = .18, *p* = .67] or diet [*F*_(1, 19)_ = .25, *p* = .62]. **(B)** Hippocampal hemisphere was affected by lateralization [*F*_(1, 19)_ = 6.08, *p* = .023] and diet [*F*_(1, 19)_ = 4.71, *p* = .043]. WD rats exhibited a reduced right hippocampal volume relative to the left hippocampal volume in CD rats (*p* = .019). Sample size = 6 rats / group.

**Supplemental Figure 3. pErbB4 is expressed in macrophages/microglia in the hippocampus of rats treated with NRG1.**

**Supplemental Figure 4. PC analysis reveals distinctive microglial morphological descriptors in each group. (A-D)** Principal Component (PC) analysis on the morphometric parameters to trace the possible differences in microglia driving the changes in hippocampal structure and behavior. These PCs explained more than 80% of the accumulated variance between cells (PC1: ~60%; PC2: ~20%; PC3: ~5%). Sample size approximately 500 microglia / group.

**Supplemental Figure 5. PCA reveals distinctive microglial morphological descriptors contributing to differences between groups.** Seven morphometric parameters describing microglial shape were subjected to PCA and there were distinctive morphological profiles between groups (PC1 = 58, PC2 = 35%). Sample size = 3 rats / group.

**Supplemental Figure 6. ErbB4 downstream mediators are not affected significantly. (A)** pAkt levels (normalized to total Akt) were not affected by the treatment [*F*_(1, 17)_ = .16, *p*= .69] or diet [*F*_(1, 17)_ = 2.80, *p* = .12]. **(B)** pErk1/2 levels (normalized to total Erk) were not affected by the treatment [*F*_(1, 18)_ = .88, *p* = .36] or diet [*F*_(1, 18)_ = .35, *p* = .56]. Sample size = 5-6 rats / group.

**Supplemental Figure 7. NRG1 levels are not significantly altered.** Twenty-one (21) days consuming the WD (diet [*F*_(1, 18)_ = .38, *p* = .54]) and receiving NRG1 (treatment [*F*_(1, 18)_ = .031, *p* = .86]) administration was not sufficient to significantly alter NRG1 protein levels in the hippocampus, as measured by ELISA. Sample size = 5-6 rats / group.

**Supplemental Figure 8. Obesogenic diet and prolonged NRG1 treatment alter ErbB4 isoforms expression in the hippocampus.** A subgroup of rats that did not undergo behavioral manipulations was euthanized with Euthasol (Virbac) and briefly perfused with PBS. The rats were rapidly decapitated, and the hippocampi were isolated. The cDNA was amplified by PCR using the following primer sets (5’-3’):

*Gapdh*, fwd: AGTTCAACGGCACAGTCAAG,

*Gapdh*, rev: GTGGTGAAGACGCCAGTAGA;

*ErbB4-JMa*, fwd: GGACGGGCCATTCCACTTTACC,

*ErbB4-JMa*, rev: CCTCCAATGACTCCGGCTGC;

*ErbB4-cyt1*, fwd: GGAATATTTGGTCCCCCAGGCTTTC,

*ErbB4-cyt1*, rev: GAGGAGGGCTGTGTCCAATTTCAC;

*ErbB4-JMb*, fwd: CATTGAAGACTGCATCGGCCTG,

*ErbB4-JMb*, rev: CCTCCAATGACTCCGGCTGC;

*ErbB4-cyt2*, fwd: GGAATATTTGGTCCCCCAGGCTTTC,

*ErbB4-cyt2*, rev: GTACACAAACTGATTCCTATTGGAGTCAATTC.

The qRT-PCR methods have been detailed in previous publication from our laboratory (Vega-Torres et al., 2020b). Lower JMa mRNA levels in the WD rats (relative to CD rats) and higher JMb in the rats that received the exogenous NRG1 (relative to VEH). Mixed-effects model analyses revealed a significant interaction between isoform x diet [*F*_(3, 57)_ = 3.13, *p* = .033] and isoform x treatment [*F*_(3, 57)_ = 2.79, *p* = .049]. Sample size = 6 rats / group (before detecting outliers).

**Supplemental Table 1**

|  | **CD** | **WD** |
| --- | --- | --- |
| **Macronutrient (main source)** | **% *kcal*** | **% *kcal*** |
| Carbohydrates (corn starch) | 64.7 | 43.1 |
| Protein (casein) | 18.8 | 15.5 |
| Fat (milk fat) | 16.5 | 41.4 |
| ***Total kcal*** | 3.77 | 4.57 |
| **Fatty Acid Class** | **g/kg Diet** | **g/kg Diet** |
| Butyric (C4:0) | 0.72 | 5.10 |
| Caproic (C6:0) | 0.52 | 3.55 |
| Caprylic (C8:0) | 0.33 | 2.21 |
| Capric (C10:0) | 0.78 | 5.35 |
| Lauric (C12:0) | 0.93 | 6.30 |
| Myristic (C14:0) | 3.00 | 20.3 |
| Myristoleic (C14:1) | 0.24 | 1.67 |
| Pentadecanoic (C15:0) | 0.33 | 2.18 |
| Palmitic (C16:0) | 11.6 | 61.6 |
| Palmitoleic (C16:1) | 0.46 | 2.84 |
| Heptadecanoic (17:0) | 0.17 | 1.04 |
| Stearic (C18:0) | 3.52 | 20.2 |
| Oleic (C18:1) | 11.2 | 39.8 |
| Linoleic (C18:2) | 11.5 | 10.5 |
| Alpha Linolenic (C18:3) | 0.31 | 0.83 |
| Arachidic (C20:0) | 0.11 | 0.28 |
| Homogamma Linolenic (C20:3) | <0.07 | 0.20 |
| Arachidonic (C20:4 n-6) | <0.07 | 0.30 |
| EPA (C20:5 n-3) | <0.07 | <0.14 |
| DHA (C22:6 n-3) | <0.07 | <0.14 |
| **Fatty Acid Class** | **g/kg Diet** | **g/kg Diet** |
| Saturated | 20.8 | 121 |
| Monosaturated | 12.7 | 492 |
| Polyunsaturated | 11.3 | 11.5 |
| Omega-3 Fatty Acids | 0.31 | 0.94 |
| Omega-6 Fatty Acids | 11.5 | 10.1 |

**Supplemental Table 2**

| **Target** | **Company (Product)** | **Dilution** |
| --- | --- | --- |
| ADAM17 | Abcam (ab13535) | 1:2500 |
| Akt | Cell Signaling (2920) | 1:200 |
| β-actin | LI-COR (926-42212) | 1:2000 |
| EGFR (ErbB1) | Invitrogen (PA1-1110) | 1:500 |
| ErbB2 | Invitrogen (MA5-13105) | 1.5 𝜇g/mL |
| ErbB3 | Invitrogen (MA5-12675) | 3 𝜇g/mL |
| ErbB4 | Abcam (ab32375) | 1:1000 |
| Iba-1 | Wako (019-19741) | 1:750 |
| NRG1 | Invitrogen (MA5-12896) | 2 𝜇g/mL |
| p44/42 MAPK (Erk1/2) | Cell Signaling (9102) | 1:1000 |
| Phospho-Akt | Cell Signaling (2965) | 1:1000 |
| Phospho-ErbB4 (Y1284) | Abcam (ab61059) | 1:1000 |
| Phospho-p44/42 MAPK (Erk1/2) (Thr202/Tyr204) | Cell Signaling (9101) | 1:1000 |

**Supplemental Table 3. Summary of Microglial Morphological Parameters**

| **Parameters** | **Measure** | **Unit** |
| --- | --- | --- |
| Objects | Cell number | *Cell* |
| Density | $\frac{\# of pixels within cell outline}{Convex Hull Area}$ | $\frac{\# of pixels}{{pixels}^{2}}$ |
| **Span Ratio** | $\frac{\boldsymbol{Convex Hull Elipse Longest Lenght}}{\boldsymbol{Convex Hull Elipse Longest Width}}$ | ***Ratio (a.u)*** |
| **Maximum Span Across Hull** | **Maximum distance between two points across the Convex Hull** | ***pixels*** |
| **Convex Hull Area** | **Area of the polygon containing the whole cell shape** | ***pixels^2^*** |
| Perimeter | Length of the single outline of the cell shape | *pixels* |
| **Circularity** | $\frac{\boldsymbol{4}\boldsymbol{\pi(cell area)}}{\left( \boldsymbol{cell perimeter} \right)^{\boldsymbol{2}}}$ | ***pixels*** |
| Width of Bounding Rectangle | Length of the width of the largest bounding rectangle | *pixels* |
| Height of Bounding Rectangle | Length of the height of the largest bounding rectangle | *pixels* |
| Max Radius from Hull's Centre of Mass | Radius of the largest circle bounding the convex hull area | *pixels* |
| Max/Min Radii | $\frac{Radius of the largest bounding circle}{radius of the smallest bounding circle}$ | *Radio (a.u)* |
| Mean Radius | $\frac{Max Radius+Min radius}{2}$ | *pixels* |
| Diameter of Bounding Circle | Diameter of the largest circle bounding the cell | *pixels* |
| Maximum Radius | Radius of the largest circle bounding the cell | *pixels* |
| CV for all Radii from Cicle's Centre | Coeficient of variability for all the radii of all the circles bounding the cell | *Index of variability (a.u)* |
| CV for all Radii | Coeficient of variability for all the radii of all the circles bounding the convex hull area | *Index of variability (a.u)* |
| Mean Radii | Average of all radii from all bounding circles within the convex hull area | *pixels* |
| Lacunarity (Л) | Coefficient of variation expressed as pixel density per box as a function of box size | *Index of variability (a.u)* |

*a.u = arbitrary units*

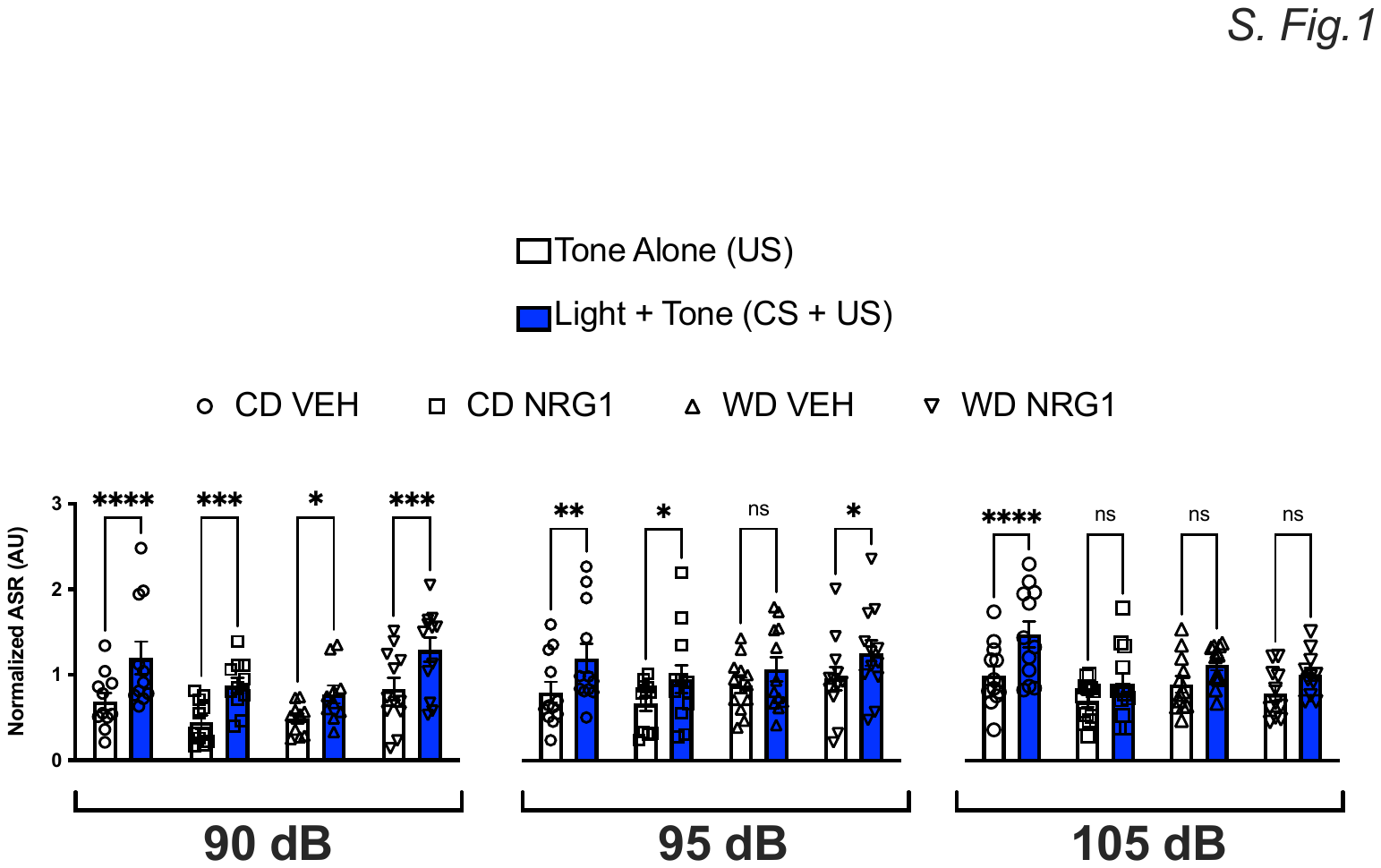

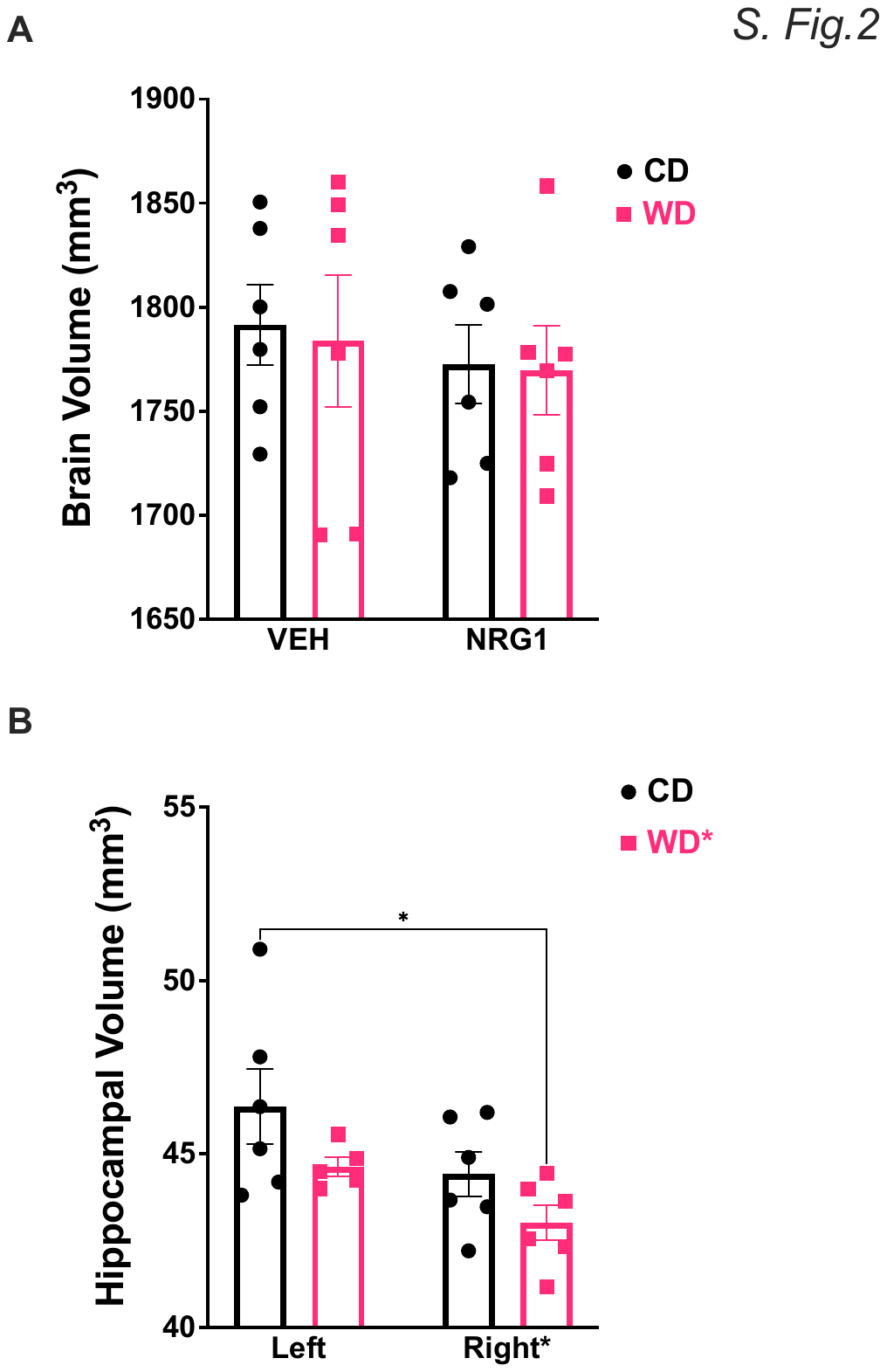

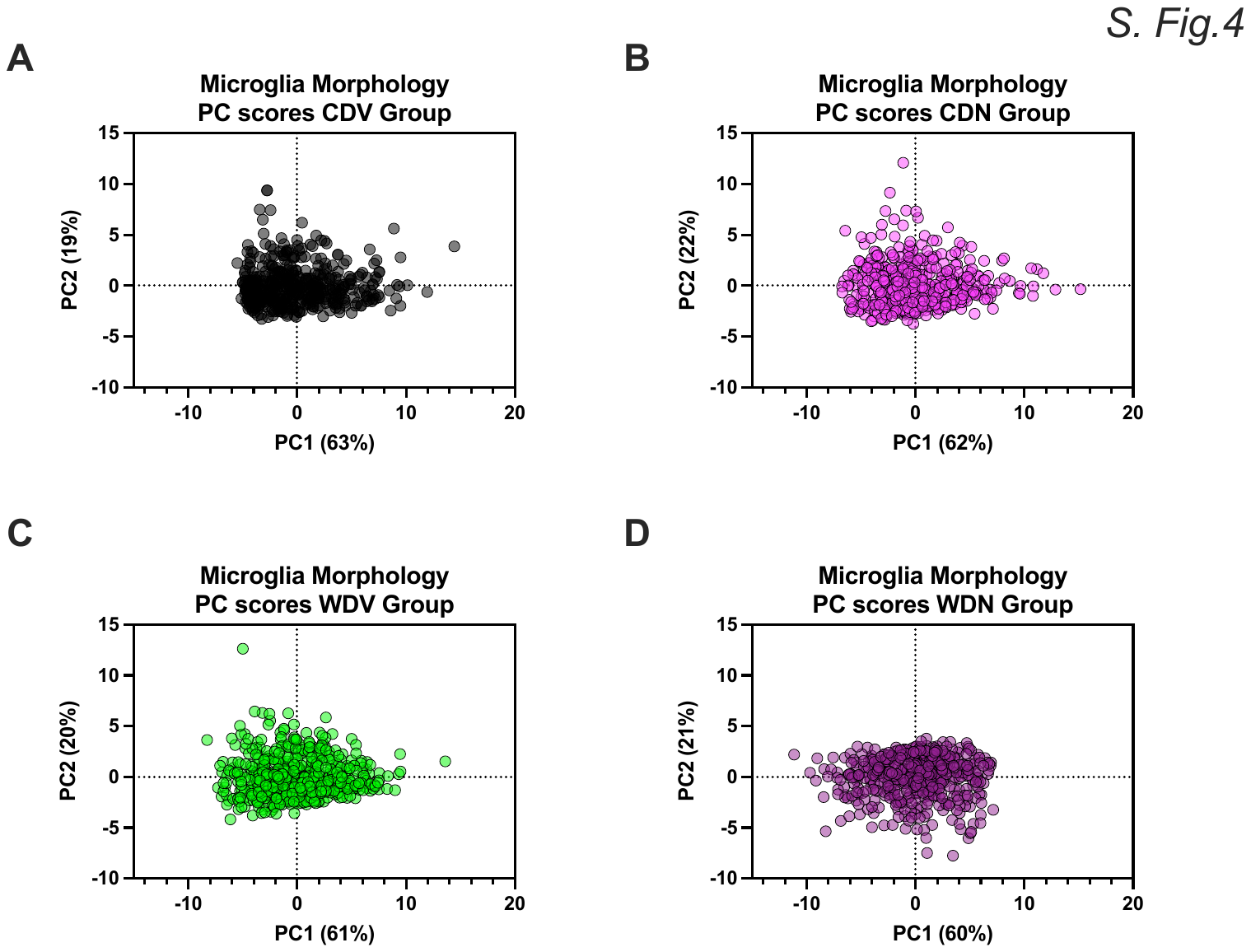

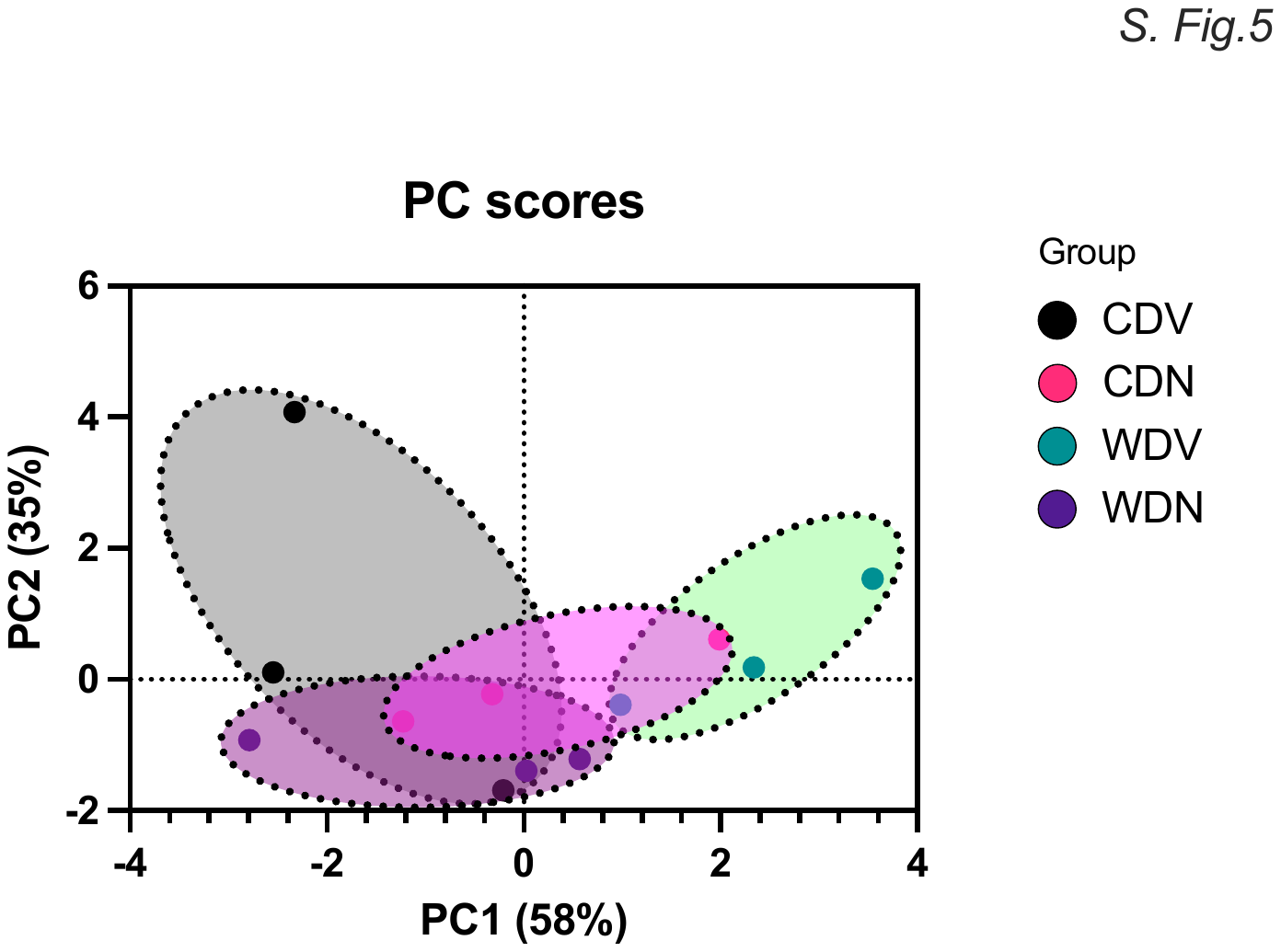

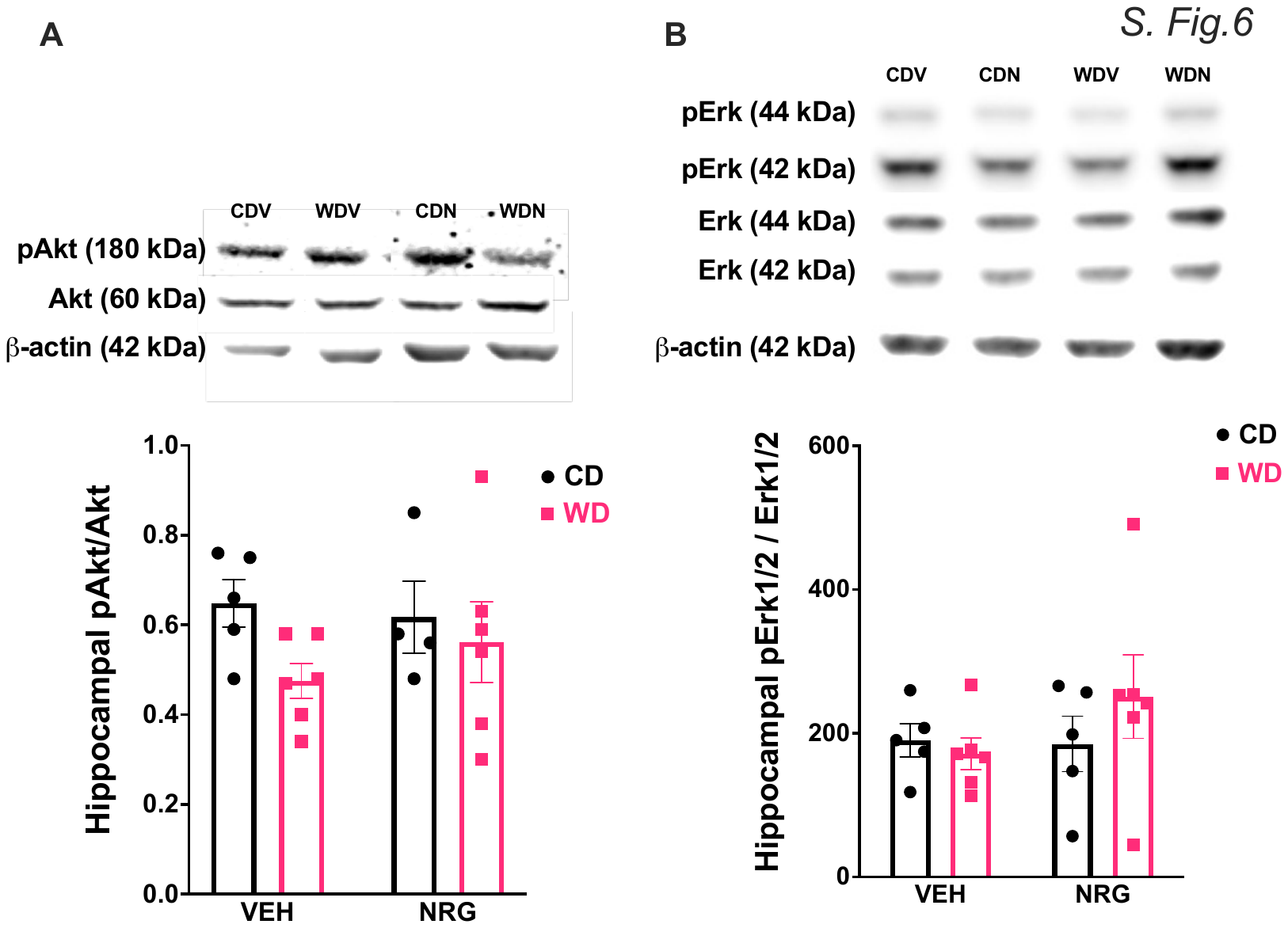

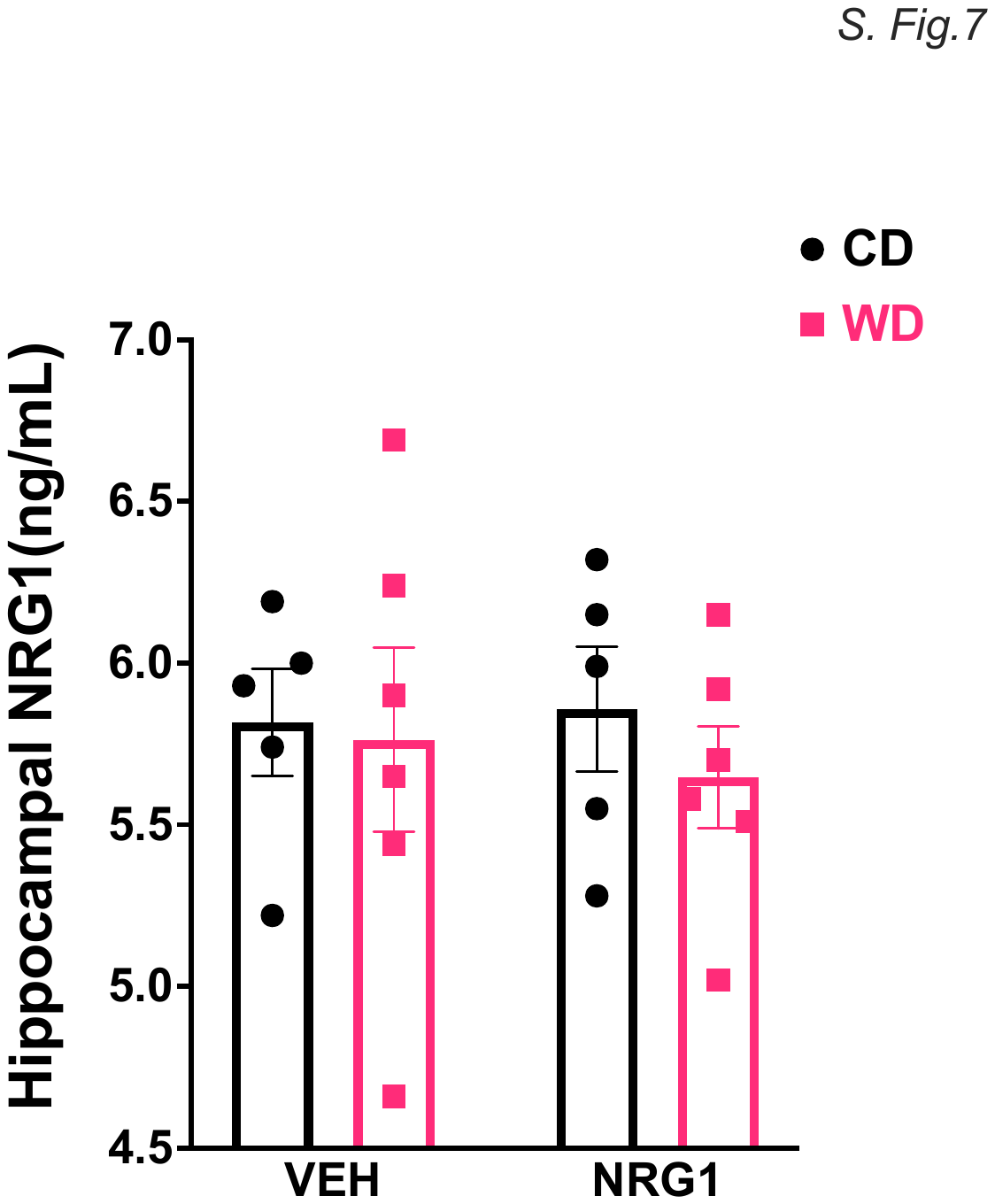

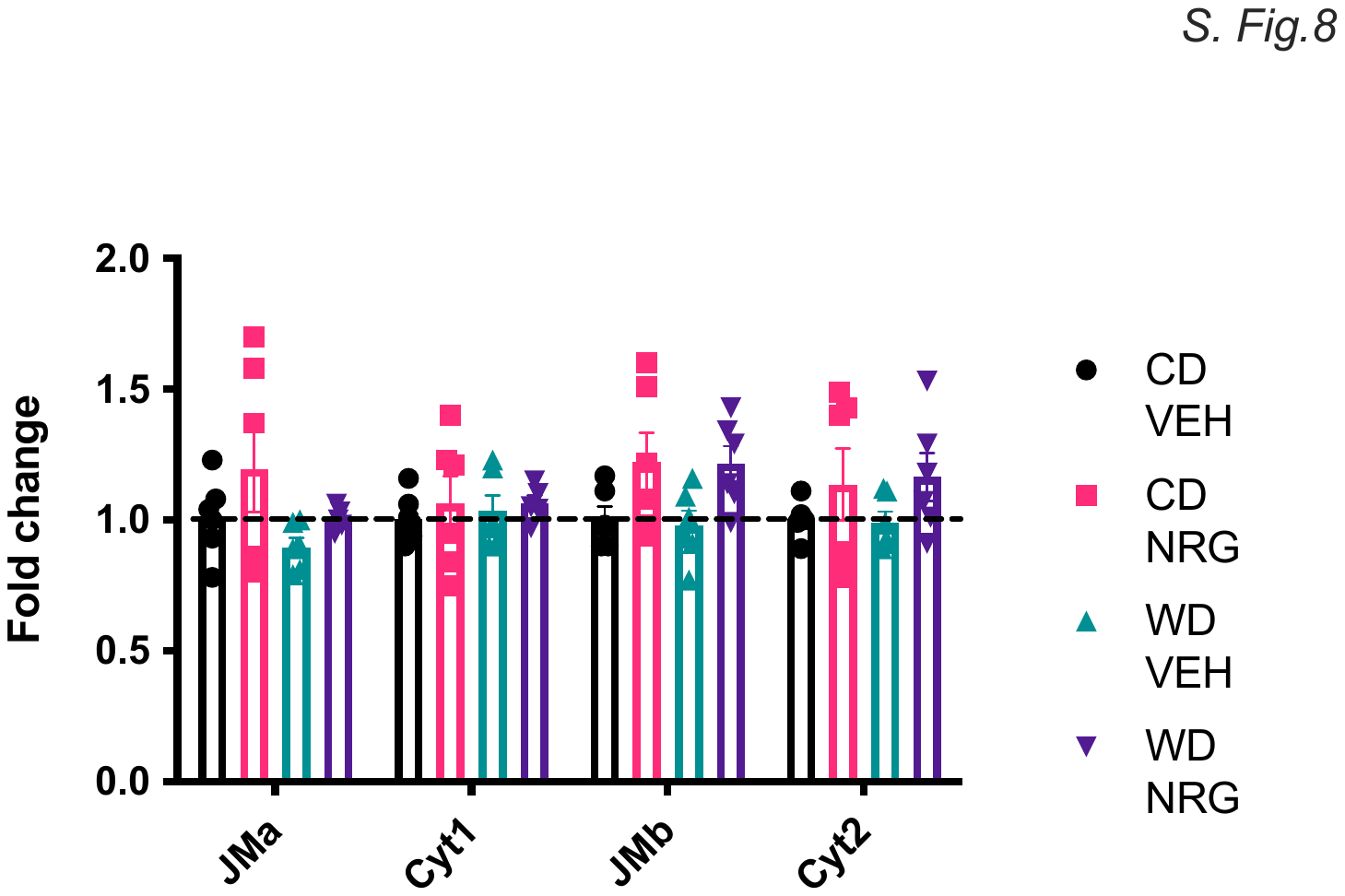
